# COP9 signalosome and PRMT5 methylosome complexes are essential regulators of Lis1-dynein based transport

**DOI:** 10.1101/2025.04.08.647777

**Authors:** Devanshi Gupta, Subbareddy Maddika

**Author notes:** Corresponding author: Dr. Subbareddy Maddika.

## Abstract

Cytoplasmic dynein is a minus-end directed microtubule motor essential for retrograde transport in cells. Dynein is tightly regulated by Lis1 where it is reported to stall dynein movement as well as promote formation of active dynein in different model systems. However, the mechanism behind these contrasting functions of Lis1 in regulating dynein is unknown. Here, we identified COP9 Signalosome (CSN) and PRMT5 complexes as critical regulators of Lis1 dependent dynein function. We demonstrated that Lis1 recruits CSN on dynein to facilitate its deneddylation. We show that neddylation of dynein (DIC1 at K42 residue) is essential for assembly of an active dynein complex and subsequent transport of cargoes in cells. Conversely, PRMT5 is recruited to the Lis1-dynein complex where it methylates Lis1. Methylation of Lis1 at R238 position alters its interaction dynamics, leading to dislodging of CSN from dynein resulting in dynein neddylation and activation. In conclusion, we unravel an interplay between CSN, and PRMT5 that provides a sophisticated regulatory mechanism for Lis1’s opposing modes of regulating the dynamics of dynein.

## Introduction

Cytoplasmic dynein-1 (hereafter dynein) is a minus-end directed microtubule motor essential for transporting cargoes like endosomes, golgi, viruses, mRNA etc ^1, 2^. Dynein complex is a ∼1.5 MDa multi-subunit assembly comprising of a dimer of motor-containing heavy chains (DHC), two intermediate chains (DIC1 and DIC2), two light intermediate chains (LIC1 and LIC2), and three light chains (LC8, LC7, and Tctex1) ^3, 4^. As a key component of the cytoskeleton, dynein is essential for numerous functions including organelle positioning, mitosis, and the maintenance of neuronal integrity ^3, 5, 6^. Its mechanistic activity and regulation are crucial for cellular homeostasis and development. Dynein requires association with dynactin and cargo adaptors, and regulators for its movement ^7–9^. Lis1 (Lissencephaly −1) is a cytoplasmic protein that interacts with dynein and affects its motor activity, thereby influencing dynein-mediated transport and cellular dynamics ^10–12^. Lis1 is conserved from yeast to humans ^13^, and is the only regulator that binds directly to the dynein’s motor domain ^14, 15^. Mutations in Lis1 or dynein give rise to neurodevelopmental defects, such as lissencephaly (a smooth brain disorder), during early developmental processes ^16–19^.

Recent advances in cell biology have underscored the intricate regulatory mechanisms governing dynein function, with Lis1 emerging as a significant modulator. Despite the crucial roles of dynein and Lis1, the precise mechanisms by which Lis1 regulates dynein activity remain complex and not fully understood. Lis1 is essential for recruitment of dynein to various cellular structures, such as nuclear envelope ^20, 21^, plus ends of microtubules ^22, 23^, and cell cortex ^24^ etc. *In vitro* single molecule studies on dynein shows that Lis1 negatively regulates dynein motility by clamping it to microtubules, prolonging its stalling, and increasing the affinity of dynein with microtubules ^15, 25–28^. Moreover, Lis1 has been shown to sterically hinder the mechanochemical cycle of dynein by binding to the stalk and preventing the release of ATP ^15^. Additionally, using *in vivo* model systems such as *Ustilago maydis*, and in Dorsal-Root-Ganglion (DRG) neurons Lis1 has been shown to keep dynein on microtubule plus ends and suppress the motility of dynein ^29, 30^. These studies suggest Lis1 may act as a negative regulator of dynein movement under certain conditions. On the contrary, multiple recent studies have established Lis1 as a positive regulator of dynein function. For instance, Lis1 was shown to relieve dynein from its autoinhibited phi conformation to an open conformation ^31–36^. As phi conformation is unable to bind to the microtubules, Lis1 enhances the localization of dynein on microtubules by opening its closed conformation ^31, 37, 38^. Moreover, recent cryo-EM studies suggests that Lis1 promotes dynein’s mechanochemical cycle in solution ^39^. Besides, Lis1 also promotes the formation of active dynein complexes by pairing it with dynactin and cargo adaptors ^31, 32^. Additionally, Lis1 favours the recruitment of the second dynein dimer for processive movements and fast velocity^31^. These contrasting roles of Lis1 function on dynein pose a question on how is Lis1 able to achieve these opposing functions in a cellular environment ^11, 40, 41^. In this study, we identified two new regulators – COP9 Signalosome and PRMT5 – with which Lis1 is able to maintain its opposing modes of regulating dynein-mediated transport in the cells.

## Results

### CSN associates with Lis1 and dynein

To identify the molecular mechanism on how Lis1 regulates dynein function, we purified Lis1 associated complexes from human cells. Tandem-affinity purification followed by mass spectrometry (TAP-MS) of a triple tagged (SFB)-Lis1 from HEK 293T cells revealed COP9 Signalosome (CSN; red circles) and PRMT5 methylosome components (green circles) as new Lis1 associated complexes along with the known interactors like dynein, dynactin, and Nde1 (blue circles) (Figure 1A & Table S1). COP9 Signalosome (CSN) is a deneddylase (removal of Nedd8) complex of 8 subunits (CSN1-8) which are conserved throughout eukaryotes ^42, 43^. The deneddylating function of CSN exclusively regulates the Cullin-RING E3 ligases (CRLs) by inactivating them and controlling the cellular ubiquitination levels ^44, 45^. However, its association with Lis1 or dynein pathway is unknown. We confirmed the endogenous association of Lis1 with CSN components in cells (Figure 1B). Also, we validated the association of Lis1 and CSN complex using exogenously expressed proteins in HEK 293T cells (Figure S1A). *In vitro* pulldown assay using bacterially purified recombinant proteins suggested that Lis1 directly binds with all the peripheral six PCI domain-containing subunits (CSN 1-4, CSN7, CSN8) but not the two core MPN domain-containing subunits (CSN5 and CSN6) (Figure S1B). Lis1 comprises of a LisH domain at its N-terminus, followed by a short stretch of coiled-coil repeats, and a set of seven WD repeats spanning till the C-terminus of the protein (Figure S1C). We made Lis1 deletion mutants (1′LisH or 1′WD) and tested their interaction with CSN to identify the Lis1 binding domain on CSN. Deletion of either LisH domain or WD repeats significantly reduced Lis1 binding with CSN subunit in cells (Figure S1D), suggesting that both the regions in Lis1 are essential for binding. In fact, our *in vitro* pulldown assay using recombinant proteins of Lis1 domain deletions suggested that Lis1 binds with some CSN subunits (CSN3 and CSN4) via its LisH domain, while it requires WD repeats to bind to other subunits such as CSN2 (Figure S1E)). This data possibly suggests that there could be multiple Lis1 molecules associated with the CSN complex, binding through both of its domains. Since Lis1-dynein exist together, we next tested if CSN associates with dynein too. Along with Lis1, CSN complex also interacts with dynein, represented by dynein intermediate chain (DIC) (Figure 1C & S1F). Interestingly, CSN binds specifically to dynein heavy chain (DHC encoded by DYNC1H1) and dynein intermediate chains (DIC1 and DIC2 encoded by DYNC1I1 and DYNC1I2 respectively) but not with other dynein subunits such as light intermediate chains (LIC1, and LIC2 encoded by DYNC1LI1 and DYNC1LI2 respectively) and light chains (LC8, Tctex1, and LC7 encoded by DYNLL1, DYNLT1 and DYNLRB1 respectively) (Figure 1D, E), possibly suggesting that CSN associates with a premature dynein complex, comprising of heavy chain and intermediate chains. We confirmed the direct association of DIC1 with CSN complex using bacterially purified recombinant proteins which suggested that DIC1 mainly binds with the peripheral subunits (CSN2, CSN3, CSN4, CSN7A and CSN8), but not with core subunits (CSN5 and CSN6) (Figure 1F). Depletion of Lis1 led to significant reduction in the interaction (Figure 1G) between CSN and dynein. However, depletion or inhibition of CSN led to no changes in the association of Lis1 and dynein (Figure S1G, H). Together, this data suggests that Lis1 facilitates CSN association with dynein.

**Figure 1:**
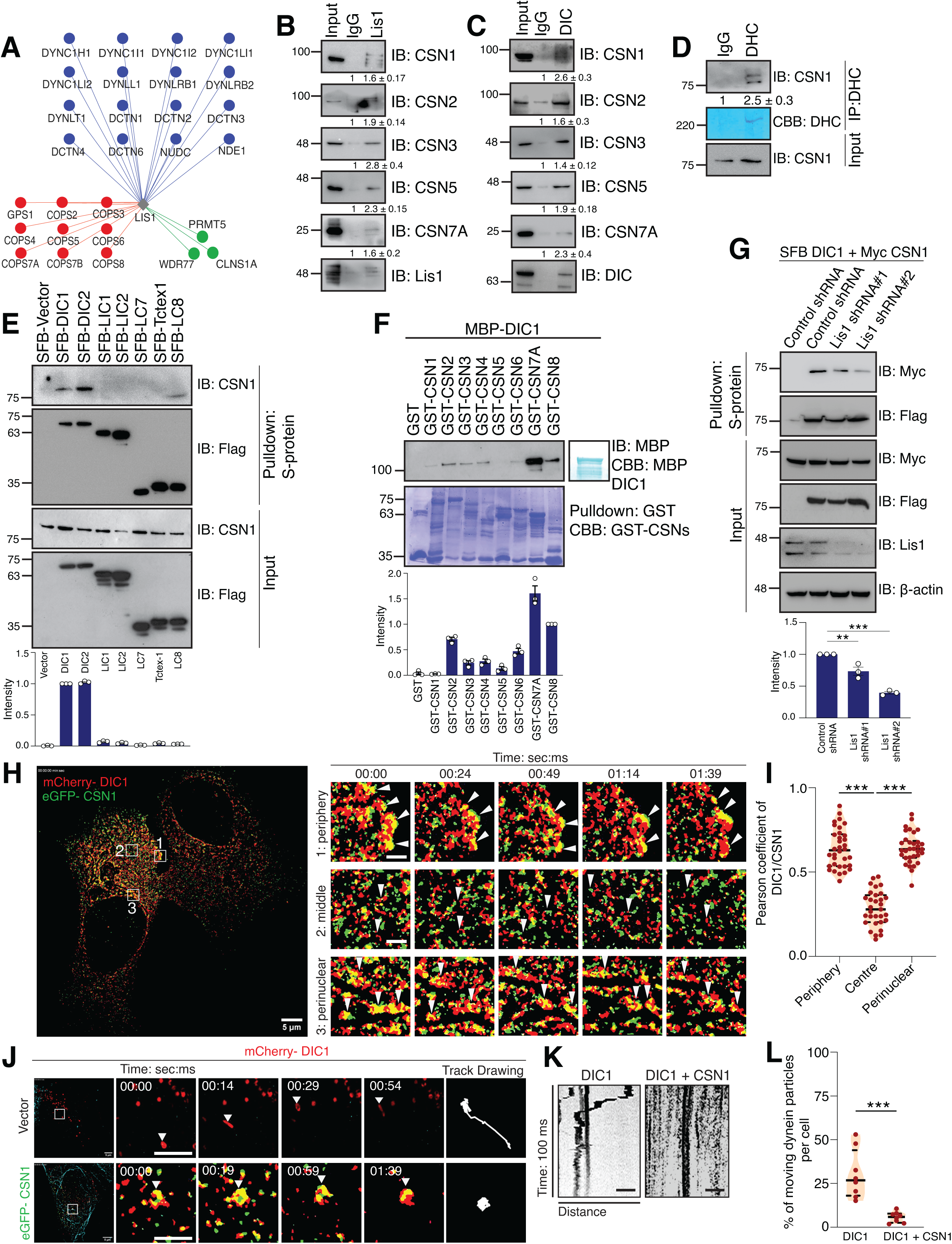
CSN interacts with Lis1 and Dynein. **(A)** Interaction network of Lis1 with its associated proteins and new complexes (COP9 Signalosome and PRMT5 methylosome) derived from mass spectrometry analysis is shown. **(B)** HEK293T cell lysates were subjected to immunoprecipitation with either IgG or Lis1. Presence of CSN subunits in Lis1 immunoprecipitates was detected by western blotting with respective antibodies. The densitometric quantifications from three biological replicates are shown under each blot. **(C)** HEK293T cell lysates were subjected to immunoprecipitation with either IgG or DIC. Presence of CSN subunits in DIC immunoprecipitates was detected by western blotting with respective antibodies. The densitometric quantifications from three biological replicates are shown under each blot. **(D)** HEK293T cell lysates were subjected to immunoprecipitation with either IgG or DHC. Presence of CSN1 in DHC immunoprecipitates was detected by western blotting with the respective antibody. CBB: Coomassie Brilliant Blue. DHC was detected at a lower molecular weight than the full-length protein. The densitometric quantifications from three biological replicates are shown under each blot. **(E)** HEK 293T cells were transfected with the dynein subunits, DIC1, DIC2, LIC1, LIC2, LC7, Tctex1 and LC8 plasmids. Cells were lysed, and incubated with S-protein beads. Interaction was detected by immunoblotting with CSN1 antibody. The densitometric quantifications from three biological replicates are shown in the graph below. **(F)** Bacterially purified recombinant GST-CSN subunits were incubated with purified MBP-DIC1. The eluates were immunoblotted by MBP antibody. CBB: Coomassie Brilliant Blue. The densitometric quantifications from three biological replicates are shown in the graph below. **(G)** Control and Lis1 depleted stable HEK 293T cells were co-transfected with SFB-DIC1 and Myc-CSN1 plasmids. Cells were then lysed, and incubated with S-protein beads. Interaction was detected by immunoblotting with anti-Myc antibody. The densitometric quantifications from three biological replicates are shown in the graph below. (**p = 0.0061, ***p < 0.0001; One way ANOVA followed by Dunnett’s multiple comparisons test). **(H)** U2OS cells were transfected with mCherry-DIC1 and eGFP-CSN1 to visualize their colocalization through time-lapse live-cell super-resolution microscopy with SIM module. Three ROIs were drawn at different areas in the cell – near periphery, in the middle, and near nucleus, and their respective video stills has been shown. **(I)** The quantification of the Pearson coefficient of the colocalization at the three ROIs from (n=30 cells) three biological replicates are shown in the graph. (***p < 0.0001; One way ANOVA followed by Dunnett’s multiple comparisons test) **(J)** U2OS cells were transfected with mCherry-DIC1 and eGFP-CSN1 to track the movement of individual moving dynein using time-lapse live cell super-resolution microscopy with SIM module. The video stills, track drawing, and **(K)** their corresponding kymographs are shown. Scale bar: 1 um. **(L)** The quantification of the percentage of moving dynein particles in each sample (n=8 cells) for the experiment in (J) from two biological replicates are shown in the graph. (***p = 0.0003; Two-tailed t-test)

In cells, dynein can exist in either microtubule bound state or in premature state in cytoplasm. To test if CSN associates with either of these forms of dynein, we manipulated the microtubules by treating the cells with either nocodazole (depolymerizes microtubules leading to immotile dynein), or paclitaxel (prevents the microtubule depolymerization leading to efficient dynein movement). CSN colocalizes with DIC1 in nocodazole treated cells, which is significantly reduced in paclitaxel treated cells (Figure S2A, B). Further, we tested the colocalization of CSN and DIC1 in temperature sensitive conditions that alters microtubule polymerization dynamics. Co-localization of CSN with DIC1 is significantly enhanced upon cold shock where microtubules are dispersed. However, CSN-DIC1 colocalization is reduced upon recovery to ambient temperature with addition of warm media (Figure S2C, D), clearly suggesting that CSN predominantly binds with premature dynein in cytoplasm. Additionally, we tested the colocalization of CSN1 with components like dynactin subunit p150, cargo adaptor BicD2, microtubules subunit α-tubulin, as well as with DHC (Figure S2E). Interestingly, CSN1 efficiently colocalized with DHC, but not with α-tubulin, BicD2 and p150 (Figure S2F). Moreover, CSN1 interaction with p150 and BicD2 was significantly reduced (Figure S2G) thus suggesting that CSN is associated with premature inactive dynein, and not the functional activated dynein complex on microtubules. Additionally, we tested the colocalization of mCherry-DIC1 and eGFP-CSN1 in U2OS cells using live-cell microscopy. The snapshots of the video at different timelines across the three ROIs in the cell clearly suggested that CSN is associated with DIC1 at the cell periphery and perinuclear regions but not in the middle region (Figure 1H, I and Video S1). Moreover, tracking the individual dynein either alone or in the presence of CSN, revealed that CSN restricts the movement of dynein in the cells (Figure 1J-L, S2H and Video S2). In conclusion, this data suggests that CSN binds with premature dynein in cytoplasm and thereby restricts its movement in cells.

### CSN deneddylates dynein subunit, DIC1

CSN is known to regulate substrate ubiquitination and stability via controlling CRL neddylation. Since we found several Cullin proteins in Lis1 purification along with CSN, we first tested if dynein levels are regulated via CSN-CRL axis. Dynein subunit DIC1 binds to different Cullins (Cul1, Cul4A and Cul7) (Figure S3A), While Cullin expression led to Cdt1 (known substrate) degradation ^46–48^ (Figure S3B), neither over expression of CSN1 affected the stability of various dynein subunits (Figure S3C), nor increased concentration of Cullins in cells affected the steady state levels of endogenous DIC (Figure S3D). This suggests that DIC association with CSN is independent of CSN-CRL protein degradation axis. As CSN is a deneddylating enzyme, we next tested if dynein is a new substrate of CSN. Firstly, we tested if dynein components undergo neddylation. Immunoprecipitation using endogenous DIC followed by immunoblotting with Nedd8-specific antibody suggested that DIC gets neddylated (Figure 2A). Next, immunoprecipitation using Nedd8-specific antibody suggested that DIC1 is specifically neddylated in cells (Figure 2B). Subsequently, an *in vitro* neddylation and deneddylation assay on purified GST-DIC1 indicated that CSN readily deneddylates DIC1 (Figure 2C). Further, treatment of cells with CSN5i3 (CSN inhibitor) led to accumulation of neddylated DIC1 (Figure 2D). On the other hand, depletion of CSN5, a catalytic subunit of CSN caused a significant increase in neddylation levels of DIC1 in cells (Figure 2E). Thus, this data suggests that DIC1 gets neddylated in cells and is a deneddylation substrate of CSN. As Lis1 recruited CSN to dynein, we next tested if Lis1 participates in DIC1 deneddylation. *In vitro* deneddylation assay suggested that Lis1 significantly enhanced the deneddylation of DIC1 by CSN (Figure 2F, S3E). Together, this data suggests that Lis1 facilitates CSN dependent deneddylation of dynein subunit DIC1. Next, we sought to find the site of neddylation on DIC1. Using a deep-transfer learning neddylation prediction tool (DTL-NeddSite), we narrowed down to two lysine residues, K42 and K43 on DIC1, which we mutated to arginine residues and tested for the loss of neddylation. We saw a significant decline in neddylation levels with DIC1 K42R mutant (Figure 2G), suggesting that K42 may be the major neddylation site in DIC1.

**Figure 2:**
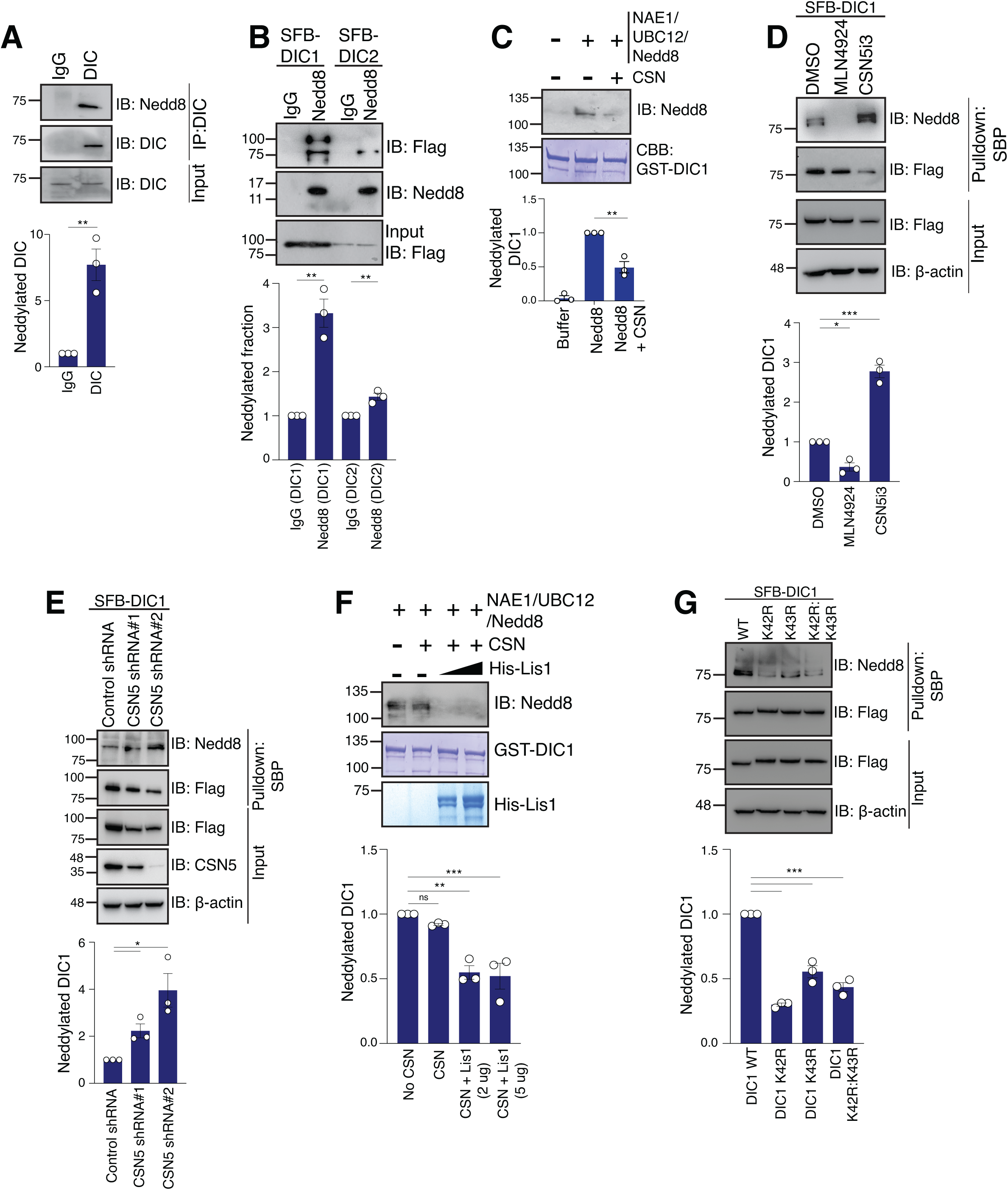
DIC1 gets neddylated and is deneddylated by CSN. **(A)** HEK293T cell lysates were subjected to denaturing immunoprecipitation with either IgG or DIC. Presence of neddylated DIC in the immunoprecipitates was detected by western blotting with Nedd8 antibody. The densitometric quantifications from three biological replicates are shown in the graph below. (**p = 0.0049; Two-tailed t-test). **(B)** HEK 293T cells were transfected with SFB-DIC1 and SFB-DIC2 plasmids. Cells were then lysed under denaturing conditions, and incubated with IgG and Nedd8-bound Protein G beads. Neddylated protein was detected by immunoblotting with anti-Flag antibody. The densitometric quantifications from three biological replicates are shown in the graph below. (**p = 0.0019, **p = 0.0058; Two-tailed t-test). **(C)** Bacterially purified GST-DIC1 was incubated with NAE1, UBC12 and Nedd8 for *in vitro* neddylation for 60 mins, and with purified CSN for *in vitro* deneddylation for 60 mins. Neddylated protein was visualised by immunoblotting with anti-Nedd8 antibody. CBB: Coomassie Brilliant Blue. Quantification is shown in the graph below. (**p = 0.0037; Two-tailed t-test). **(D)** HEK 293T cells transfected with SFB-DIC1 plasmid were treated with DMSO, 1 uM MLN4924 and 1 uM CSN5i3 for 12 hours. 24 hours after transfection, cells were lysed under denaturing conditions, and further incubated with streptavidin-sepharose beads. Neddylated proteins were detected by immunoblotting with anti-Nedd8 antibody. The densitometric quantifications from three biological replicates are shown in the graph below. (*p = 0.0114, ***p = 0.0001; One way ANOVA followed by Dunnett’s multiple comparisons test). **(E)** Control and CSN5 depleted HEK 293T cells were transfected with SFB-DIC1 plasmid. Cells were then lysed under denaturing conditions, and further incubated with streptavidin-sepharose beads. Neddylated proteins were detected by immunoblotting with anti-Nedd8 antibody. The densitometric quantifications from three biological replicates are shown in the graph below. (*p = 0.0141; Two-tailed t-test). **(F)** Bacterially purified GST-DIC1 was incubated with NAE1, UBC12 and Nedd8 for *in vitro* neddylation for 60 mins, and with purified CSN for *in vitro* deneddylation for 30 mins. Increasing amounts (2 ug, 5 ug) of His-Lis1 was added during deneddylation to see the kinetics of the reaction. Neddylated protein was visualized by immunoblotting with anti-Nedd8 antibody. CBB: Coomassie Brilliant Blue. The densitometric quantifications from three biological replicates are shown in the graph below. (ns = not significant, **p = 0.0013, ***p = 0.0008; One way ANOVA followed by Dunnett’s multiple comparisons test). **(G)** HEK 293T cells were transfected with SFB-DIC1 WT, K42R, K43R and K42:43R plasmids. Cells were then lysed, and incubated with streptavidin-sepharose beads. Neddylated protein was detected by immunoblotting with anti-Nedd8 antibody. The densitometric quantifications from three biological replicates are shown in the graph below. (***p < 0.0001; One way ANOVA followed by Dunnett’s multiple comparisons test).

### Neddylation of DIC1 is essential for Dynein movement

Dynein mediates minus-end directed movement of cargoes, such as retrograde movement of endosomes, lysosomes and is essential for maintaining golgi architecture and dynamics in the cell ^6, 49^. In non-polarized cells, dynein activity results in organelle movement from the periphery to the cell center (i.e. centripetal) ^2, 50^. Therefore, to test the role of neddylation/deneddylation of DIC1, we next sought to assess the organelle positioning as a readout for dynein activity (Figure S4A, B). As expected, while control cells display predominantly perinuclear positioning of lysosomes, endosomes and Golgi apparatus due to active dynein, depletion of Lis1 led to redistribution of lysosomes and endosomes along with Golgi architectural defects (Figure S4C, D). Similar to defects in organelle positioning caused by Lis1 depletion, treatment of U2OS cells with a neddylation inhibitor (MLN4924) caused significant increase in the distribution of organelles away from the perinuclear region in the cells (Figure 3A, B). Similar organelle position defects were observed in another cell line IMR32 with the loss of Lis1 (Figure S4E, F) or inhibition of neddylation (Figure S5A, B). Further, to understand the dynein movement in these conditions, we tested the localization of DIC1 in MLN4924 and CSN5i3 treated conditions. We found that, while DIC1 is located throughout the cytoplasm in the control cells, neddylation inhibition strictly kept DIC1 near periphery, but deneddylation inhibition concentrated it in the perinuclear space (Figure 3C, D), thus supporting our data on distribution of organelles under the same conditions. Next to understand the effect of neddylation and deneddylation on dynamicity of dynein, we employed a FKBP-FRB system-based peroxisome motility assay by utilizing two constructs Pex3-mEmerald-FKBP and FRB-BicD2S along with rapamycin, as described earlier to assess dynein movement in cells ^32, 51^. Upon treatment with MLN4924 there was a sharp decline in the speed of the peroxisomes and their net displacement (Figure 3E-G, S5 C, D and Video S3). Conversely, exogenous expression of CSN1 has significantly reduced the movement, the speed of peroxisomes, and their net displacement in cells (Figure S5E-H and Video S4).

**Figure 3:**
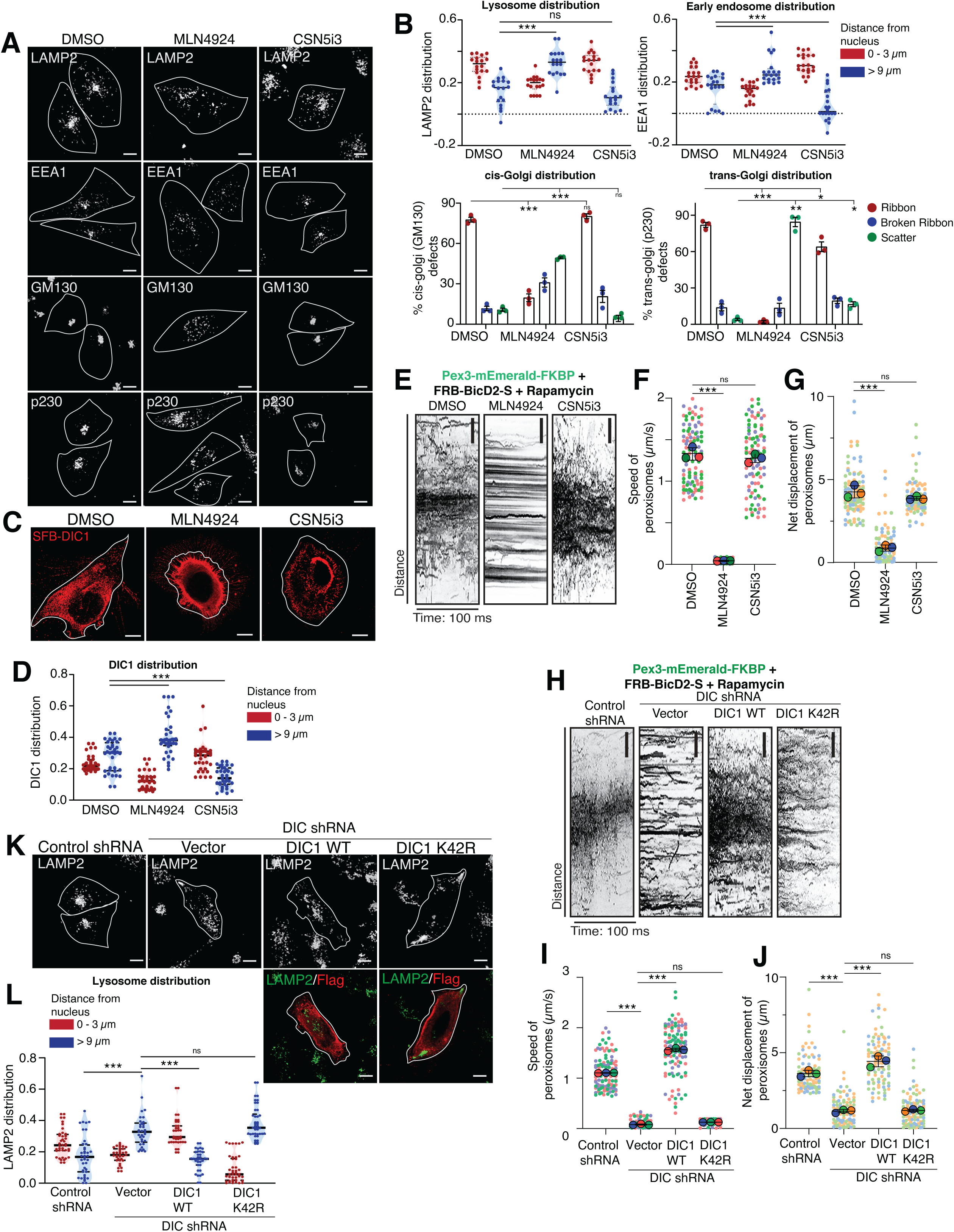
Neddylation of DIC1 is necessary for dynein movement and for proper organelle positioning. **(A)** U2OS cells were treated with DMSO, 3 uM MLN4924, and 3 uM CSN5i3 for 4 hours before fixation, and labelled with LAMP2 (lysosomes), EEA1 (early endosome), GM130 (cis-Golgi) and p230 (trans-Golgi) antibodies to visualize their distribution under confocal microscope. Representative images are shown. Scale bars: 10 µm. **(B)** Quantification of the distribution of LAMP2-positive compartments (n=18 cells), EEA1-positive compartments (n=20 cells) across three biological replicates for the experiments shown in (A). (***p < 0.0004; ns not significant; One way ANOVA followed by Dunnett’s multiple comparisons test). The golgi distribution has been categorized into 3 types: ribbon, broken ribbon and scatter. The values plotted are the mean ± SE from three independent experiments. (***p < 0.0004; ** p 0.0021; *p 0.0205; ns not significant; Two-way ANOVA followed by Dunnett’s multiple comparisons test). **(C)** U2OS cells were transfected with SFB-DIC1 and treated with DMSO, 3 uM MLN4924, and 3 uM CSN5i3 for 4 hours before fixation, and subsequently staining with anti-flag antibody to visualize the localization. Scale bars: 10 µm. **(D)** Quantification of the distribution of SFB-DIC1 (n = 36 cells) across three biological replicates for the experiments shown in (C). Data are median ± interquartile range. (\*\*\**p* < 0.0001; One way ANOVA followed by Dunnett’s multiple comparisons test). **(E)** U2OS cells were transfected with Pex3-mEmerald-FKBP and FRB-BicD2-S. After 20 hours of transfection, cells were treated with DMSO, 3 uM MLN4924, and 3 uM CSN5i3 for 4 hours, followed by 1 uM Rapamycin just before live-cell imaging. Whole cell kymographs of peroxisomes representative of each condition are shown. Scale: 1 um. **(F)** Quantification of the speed of peroxisomes in U2OS cells for the experiment shown in (E). Each color represents a replicate. Bigger circle is the mean of the data points represented as smaller circles from each biological replicate (n = 35, 25, 40 cells). Data are mean ± SE. (***p < 0.0001; One way ANOVA). **(G)** Quantification of net displacement of peroxisomes in U2OS cells for the experiment shown in (E). Each color represents a replicate. Bigger circle is the mean of the data points represented as smaller circles from each biological replicate (n = 35, 25, 40 cells). Data are mean ± SE. (***p < 0.0001; One way ANOVA). **(H)** Control and DIC depleted U2OS cells were transfected with Pex3-mEmerald-FKBP and FRB-BicD2-S along with DIC1 WT and DIC1 K42R plasmids. After 24 hours of transfection, cells were treated with 1 uM Rapamycin just before live-cell imaging. Whole cell kymographs of peroxisomes representative of each condition are shown. Scale: 1 um. **(I)** Quantification of the speed of peroxisome in U2OS cells for the experiment shown in (H). Each color represents a replicate. Bigger circle is the mean of the data points represented as smaller circles from each biological replicate (n = 40, 30, 42 cells). Data are mean ± SE. (***p < 0.0001; ns not significant, two-tailed Student’s *t*-test). **(J)** Quantification of the net displacement of peroxisome in U2OS cells for the experiment shown in (H). Each color represents a replicate. Bigger circle is the mean of the data points represented as smaller circles from each biological replicate (n = 40, 30, 42 cells). Data are mean ± SE. (***p < 0.0001; ns not significant, two-tailed Student’s *t*-test). **(K)** Control and DIC depleted U2OS cells were transfected with DIC1 WT and DIC1 K42 plasmids. Cells were fixed and labelled with LAMP2 (lysosomes) antibody to visualize the lysosome distribution under confocal microscope. Representative images are shown. Scale bars: 10 µm. **(L)** Quantification of the distribution of LAMP2-positive compartments (n = 40, 34, 36, 42 cells) across three biological replicates for the experiments shown in (K). Data are median ± interquartile range. (***p < 0.0001; *p < 0.0396; two-tailed Student’s *t*-test).

Further, to substantiate the role of neddylation on dynein movement, we overexpressed DIC1 WT and DIC1 K42R mutant in DIC depleted U2OS cells (Figure S6A) and observed the motility of peroxisomes. Wild type DIC1, but not neddylation defective K42R mutant, could rescue the movement, speed and displacement of cargoes lost in DIC depleted cells (Figure 3H-J, S6B, C and Video S5). In addition, the organelle position defects seen upon depletion of DIC1 (Figure S6D, E) could be rescued by expression of wild type DIC1 but not K42R mutant (Figure 3K, L and S7A, B). Together, this data suggests that neddylation of DIC1 at K42 residue is required for activity of dynein and Lis1-CSN keeps dynein in an inactive state by deneddylating DIC1.

### Loss of neddylation constitutes a defective dynein motor assembly

As we now know loss of neddylation impairs the dynein mediated transport of cargoes, we next sought to understand the underlying mechanism of how neddylation contributes to dynein function. Proper assembly of dynein with different subunits, and cargo adaptors is essential for a functional dynein dependent transport along the microtubules. We assessed if neddylation is important for proper dynein assembly. In order to establish the defective assembly of dynein complex in cells upon loss of neddylation, we purified the complexes associated with SFB-tagged DIC1 proteins under different conditions and identified the complexes using mass spectrometry. Treatment of cells with neddylation inhibitor or expression of K42R mutant led to significant loss of spectral counts of dynactin complex (p150, p50, p22 and p62 encoded by DCTN1-4 respectively) and LIC2, with no changes in the abundance of dynein heavy chain and light chains (Figure 4A, B). Our biochemical interaction studies showed neddylation has no effect on binding of dynein heavy chain (DHC), light intermediate chain 1 (LIC1), light chain components (LC8, Tctex1, LC7), with DIC1 (Figure S8A-C). However, association of exogenously expressed dynein light intermediate chain 2 (LIC2) with DIC1 was severely compromised in cells treated with neddylation inhibitor (Figure 4C). Also, neddylation defective K42R mutant of DIC1 display defective binding with LIC2 (Figure 4D). In addition, binding of exogenously expressed cargo adaptor (BicD2) with DIC1 was reduced under non-neddylation conditions (Figure 4E, F). Further, we confirmed the defective association of endogenous BicD2 and LIC2 with DIC1 upon neddylation inhibition (Figure 4G) or by using neddylation defective K42R mutant (Figure 4H). Defective dynein assembly in DIC1 K42R mutant conditions is due to defective neddylation but not due to general effect of K42 mutation per se as K42R mutant retains its ability to interact with known partner such as Nde1 under *in vitro* conditions (Figure S8D), with no changes in its protein stability (Figure S8E) as well as its dimerization ability (Figure S8F) in cells. This data clearly suggests that neddylation of DIC1 is essential for proper assembly of dynein complex and therefore Lis1 recruited CSN inhibits dynein mediated transport via DIC1 deneddylation.

**Figure 4:**
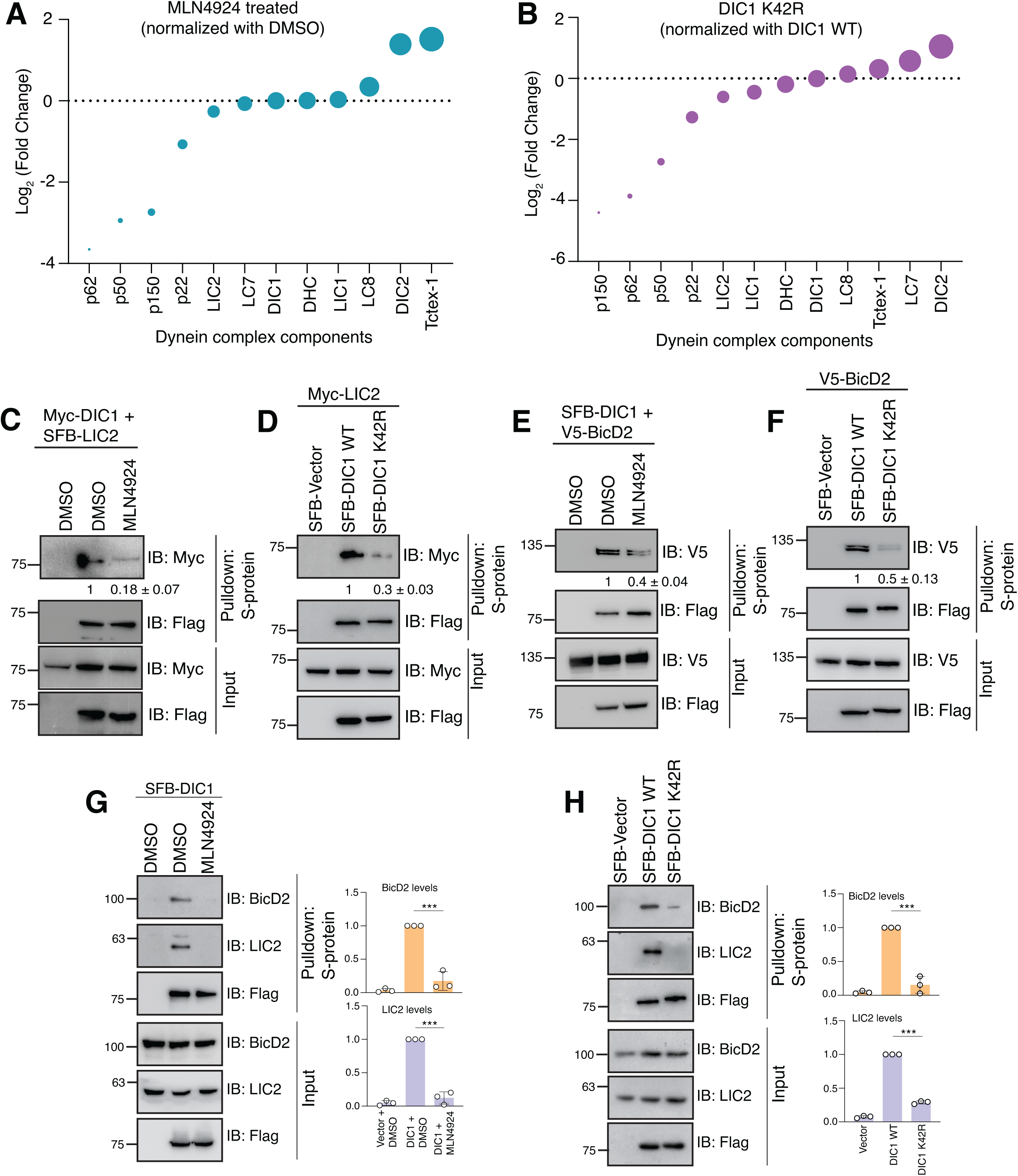
DIC1 neddylation is necessary for proper Dynein complex assembly. **(A)** Quantitation of Log_2_ Fold change of the spectral counts of each protein identified by mass spectrometry in SFB-DIC1 purified from MLN4924 treated cells and normalized with DMSO. **(B)** Quantitation of Log_2_ Fold change of the spectral counts of each protein identified by mass spectrometry in SFB-DIC1 K42R purified from cells and normalized with SFB-DIC1 WT **(C)** HEK 293T cells were co-transfected with SFB-LIC2 and Myc-DIC1 plasmids. Cells were treated with 1 uM MLN4924 (NAE1 inhibitor) or DMSO for 12 hours. Cells were lysed and further incubated with S-protein beads. Interaction was detected by immunoblotting with anti-Myc antibody. The densitometric quantifications from three biological replicates are shown under each blot. **(D)** HEK 293T cells were co-transfected with SFB-DIC1 WT and SFB-DIC1 K42R with Myc-LIC2 plasmids. Cells were then lysed, and incubated with S-protein beads. Interaction was detected by immunoblotting with anti-Myc antibody. The densitometric quantifications from three biological replicates are shown under each blot. **(E)** HEK 293T cells were co-transfected with V5-BicD2 and SFB-DIC1 plasmids. Cells were treated with 1 uM MLN4924 (NAE1 inhibitor) or DMSO for 12 hours. Cells were lysed and further incubated with S-protein beads. Interaction was detected by immunoblotting with anti-V5 antibody. The densitometric quantifications from three biological replicates are shown under each blot. **(F)** HEK 293T cells were co-transfected with SFB-DIC1 WT and SFB-DIC1 K42R with V5-BicD2 plasmids. Cells were then lysed, and incubated with S-protein beads. Interaction was detected by immunoblotting with anti-V5 antibody. The densitometric quantifications from three biological replicates are shown under each blot. **(G)** HEK 293T cells were transfected with SFB-DIC1 plasmid. Cells were treated with 1 uM MLN4924 (NAE1 inhibitor) or DMSO for 12 hours. Cells were lysed and further incubated with S-protein beads. Interaction was detected by immunoblotting with anti-BicD2 and anti-LIC2 antibodies. Quantification from three biological replicates is shown in the graph. (***p = 0.0005, ***p < 0.0001; Two tailed t-test). **(H)** HEK 293T cells were transfected with SFB-DIC1 WT and SFB-DIC1 K42R plasmid. Cells were then lysed, and incubated with S-protein beads. Interaction was detected by immunoblotting with anti-BicD2 and anti-LIC2 antibodies. Quantification from three biological replicates is shown in the graph. (***p = 0.0003, ***p < 0.0001; Two tailed t-test)..

### PRMT5 interacts with Dynein and promotes its movement

As CSN is ‘switching off’ the dynein movement, we then searched for a ‘switch on’ regulator for the dynein dependent movement. In our TAP-MS of Lis1, we found another enzymatic complex, PRMT5 methylosome complex. Protein Arginine Methyltransferase 5 (PRMT5) is a type-II symmetric di-methyltransferase ^52^, which methylates arginine residues on target proteins ^52, 53^. PRMT5 constitutes a methylosome complex comprising of a cofactor methylosome protein 50 (MEP50/WDR77) ^54^, and a substrate recognition component, like pICln ^55^, RioK1 ^56^, and COPR5 ^57^ to methylate specific substrates driving a cellular function. We tested the possibility of existence of a methylation-based switch on Lis1 that promotes dynein movement by releasing CSN from Lis1-dynein. We validated the interactions of Lis1 with each of the components of methylosome complex (Figure S9A). We also confirmed the endogenous association of Lis1 and PRMT5 in HEK 293T cells (Figure 5A) as well as in IMR32 cells (Figure S9B). Along with Lis1, endogenous PRMT5 also interacts with dynein subunit DIC (Figure 5B). Immunoprecipitation with exogenously expressed proteins additionally confirms interaction of DIC1 and DIC2 with PRMT5 methylosome complex (Figure S9C). Further, Lis1 recruits PRMT5 to Lis1-dynein complex as depletion of Lis1 led to reduced interaction (Figure 5C) with DIC1.

**Figure 5:**
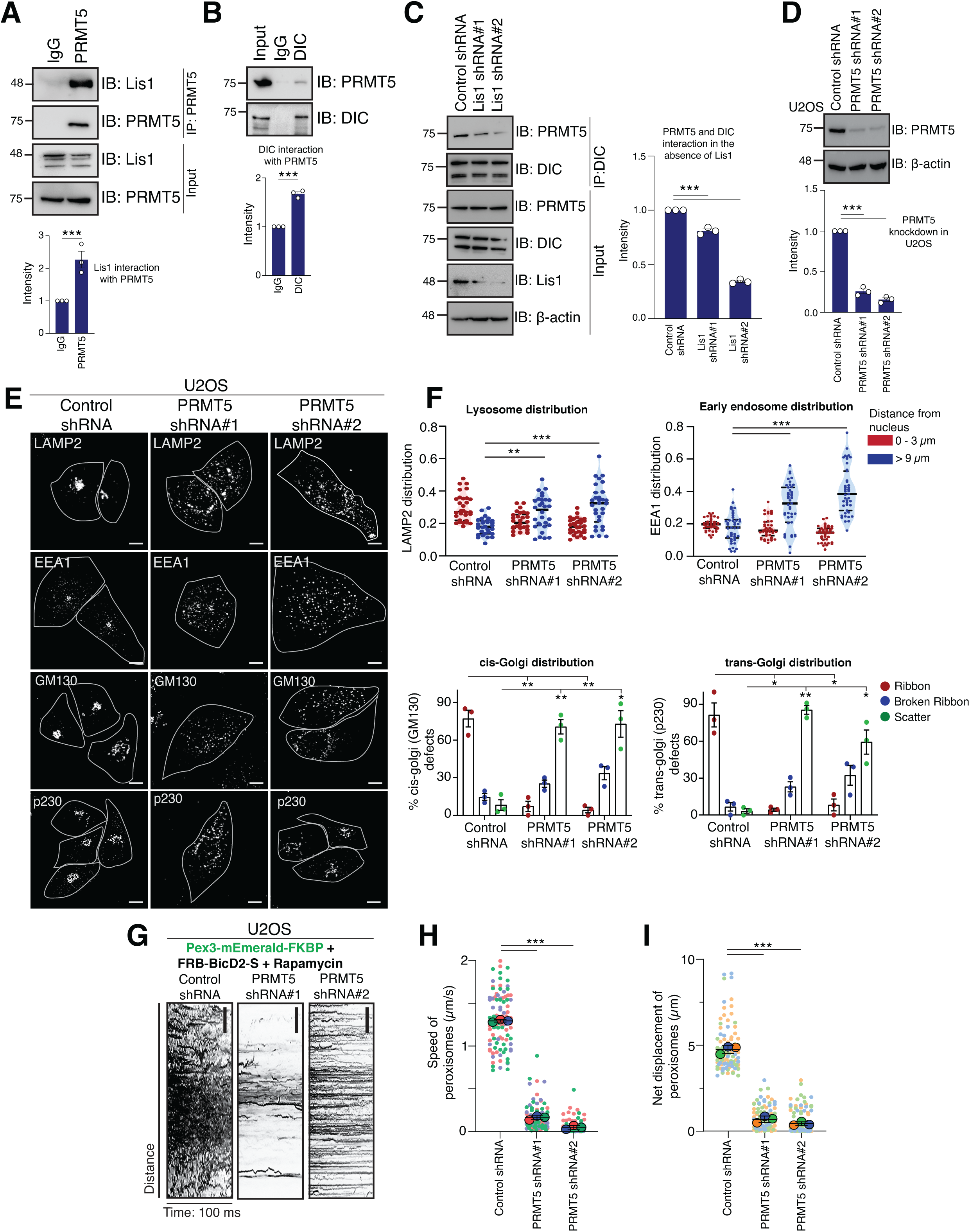
PRMT5 interacts with Lis1 and Dynein and promotes Dynein movement. **(A)** HEK293T cell lysates were subjected to immunoprecipitation with either IgG or PRMT5 antibody. Presence of Lis1 in PRMT5 immunoprecipitates was detected by western blotting with respective antibodies. The densitometric quantifications from three biological replicates are shown in the graph below. (**p = 0.0078; Two-tailed t-test). **(B)** HEK293T cell lysates were subjected to immunoprecipitation with either IgG or DIC antibody. Presence of PRMT5 in DIC immunoprecipitates was detected by western blotting with respective antibodies. The densitometric quantifications from three biological replicates are shown in the graph below. (**p = 0.0001; Two-tailed t-test). **(C)** Control and Lis1 depleted stable HEK 293T cells were immunoprecipitated with DIC antibody. Interaction was detected by immunoblotting with anti-PRMT5 antibody. The densitometric quantifications from three biological replicates are shown in the graph below. (***p < 0.0001; One way ANOVA followed by Dunnett’s multiple comparisons test). **(D)** Control and PRMT5 depleted stable U2OS cells were selected and knockdown was tested by incubating with respective antibodies. The densitometric quantifications from three biological replicates are shown in the graph below. (***p < 0.0001; One way ANOVA followed by Dunnett’s multiple comparisons test). **(E)** Control and PRMT5 depleted stable U2OS cells were fixed, and labelled with LAMP2 (lysosomes), and EEA1 (early endosome), GM130 (cis-Golgi) and p230 (trans-Golgi) antibodies to visualize their distribution under confocal microscope. Representative images are shown. Scale bars: 10 µm. **(F)** Quantification of the distribution of LAMP2-positive compartments (n=30 cells), and EEA1-positive compartments (n=43 cells) across three biological replicates for the experiments shown in (E). Data are median ± interquartile range (***p < 0.0001; **p .0020; One way ANOVA followed by Dunnett’s multiple comparisons test). The golgi distribution has been categorized into 3 types: ribbon, broken ribbon and scatter. The values plotted are the mean ± SE from three independent experiments shown in (E). (**p 0.0036, *p 0.0361, ns not significant, Two-way ANOVA followed by Dunnett’s multiple comparisons test). **(G)** Control and PRMT5 depleted U2OS cells were transfected with Pex3-mEmerald-FKBP and FRB-BicD2-S plasmids. After 24 hours of transfection, cells were treated with 1 uM Rapamycin just before live-cell imaging. Whole cell kymographs of peroxisomes representative of each condition are shown. Scale: 1 um. **(H)** Quantification of the speed of peroxisome in U2OS cells for the experiment shown in (G) Each color represents a replicate. Bigger circle is the mean of the data points represented as smaller circles from each biological replicate (n = 40, 26, 34 cells). Data are mean ± SE range. (***p < 0.0001; One way ANOVA followed by Dunnett’s multiple comparisons test). **(I)** Quantification of the net displacement of peroxisome in U2OS cells for the experiment shown in (G). Each color represents a replicate. Bigger circle is the mean of the data points represented as smaller circles from each biological replicate (n = 40, 26, 34 cells). Data are mean ± SE. (***p < 0.0001; One way ANOVA followed by Dunnett’s multiple comparisons test).

To understand the functional relevance of PRMT5 in dynein dependent functions, we next tested for the organelle positioning defects in PRMT5 depleted U2OS cells (Figure 5D). Phenocopying Lis1 loss, depletion of PRMT5 caused severe defects in the positioning of, lysosomes, endosomes, and Golgi architecture in cells (Figure 5E, F). Similar organelle position defects were observed in another cell line IMR32 with the loss of PRMT5 (Figure S9D, E). Moreover, shRNA mediated depletion of PRMT5 has significantly reduced the cargo movement, the speed and their net displacement in cells (Figure 5G-I, S9F, G, and Video S6). Also, depletion of WDR77 (Figure S10A), a cofactor of PRMT5 essential for the methyltransferase activity led to organelle position defects (Figure S10B, C). Treatment of U2OS cells with a PRMT5 inhibitor (EPZ015666) caused significant increase in the distribution of organelles away from the perinuclear region in the cells (Figure S10D, E), suggesting that PRMT5 methyltransferase activity is essential for dynein movement.

### PRMT5 methylates Lis1 and positively regulates dynein functions

Next to test if PRMT5 catalytic activity is required for dynein dependent organelle positioning, we made shRNA-resistant constructs of PRMT5 WT and its catalytic inactive mutant PRMT5 G367A:R368A, and expressed them in PRMT5-depleted cells. Wild type PRMT5, but not its catalytic inactive mutant, rescues the lysosomes and endosomes to perinuclear space and restoring Golgi architecture (Figure 6A, B, and S11A, B), clearly suggesting that PRMT5 methyltransferase activity is crucial for dynein-mediated organelle positioning. To understand how PRMT5 is involved in this process, we next sought to find the substrate of PRMT5. Since Lis1 recruited CSN as an off-switch onto dynein, we hypothesized that Lis1 may be methylated to switch from its role as a negative regulator to a positive regulator of dynein movement. To test our hypothesis, we analyzed if Lis1 gets methylated in cells. By pulling down with arginine methylation-specific antibody, we found that endogenous Lis1 gets methylated (Figure 6C). Methylation on Lis1 is enhanced by wild type PRMT5 but not a catalytic inactive G367A:R368A mutant (Figure 6D). Also, *in vitro* methyltransferase assay using bacterially purified GST-Lis1 indicated that PRMT5 wild type, but not its catalytic inactive mutant, methylates Lis1 (Figure 6E). Treatment of cells with PRMT5 inhibitor EPZ015666, as well as depleting the cofactor WDR77, reduced methylation levels of Lis1 in cells (Figure 6F and S12A), again suggesting that Lis1 is a PRMT5 substrate. Next, to find the site of methylation, we used a methylation prediction tool, PRmePred and picked two potential arginine residues on Lis1, R238 and R342. We found R238 site as a possible PRMT5 methylation site on Lis1 as mutation of R238A significantly lost methylation but no methylation changes were observed in R342A mutant (Figure 6G). Subsequently, to assess the effect of Lis1 R238 methylation on dynein movement, we made shRNA resistant mutants for these sites and tested for the rescue of organelle positioning in Lis1 depleted cells. Wild type Lis1 and R342A mutant, but not R238A mutant, could rescue the lysosome position defects in Lis1 depleted cells (Figure S12B), confirming that R238 is the site of methylation on Lis1. Interestingly, a previous study ^32^ reported that R238 residue is one of the contact sites of Lis1 on dynein heavy chain, although subsequent higher resolution structures ^13^ suggested that this residue may not be directly involved in Lis1-dynein contacts. Nonetheless, we tested if Lis1 R238 mutation has any effect on DHC interaction. No changes in the interaction of Lis1 R238A with DHC (Figure S12C) was found. Moreover, R238A mutant neither alters Lis1 protein stability (Figure S12D) nor Lis1-Lis1 dimerization (Figure S12E), clearly suggesting that organelle position defects found in Lis1 R238 mutant is due to defective methylation and not due to loss of Lis1-DHC interaction. To further analyze the importance of R238 methylation, we made another mutation, R238K (structurally similar but methylation deficient), in addition to R238F (methylation mimic) on Lis1, and tested for the organelle positioning with these mutants. We found that while Lis1 WT and Lis1 methylation mimic R238F mutant was able to rescue the lysosome position back to perinuclear region in Lis1 depleted conditions, the Lis1 R238K mutant remain defective (Figure 6H, I). Similarly, Lis1 R238K, but not R238F, was defective in rescuing the early endosome positioning, as well as the Golgi architecture in Lis1 depleted cells (Figure S13A, B). Additionally, Wild type Lis1 and R238F methylation mimic, but not methylation defective R238K mutant, could rescue the movement, speed and displacement of cargoes lost in Lis1 depleted cells (Figure 6J-L, S13C, D, and Video S7). Together this data suggests that methylation on Lis1 R238 site by PRMT5 is essential to maintain the dynein movement, failure of which leads to severe transport defects.

**Figure 6:**
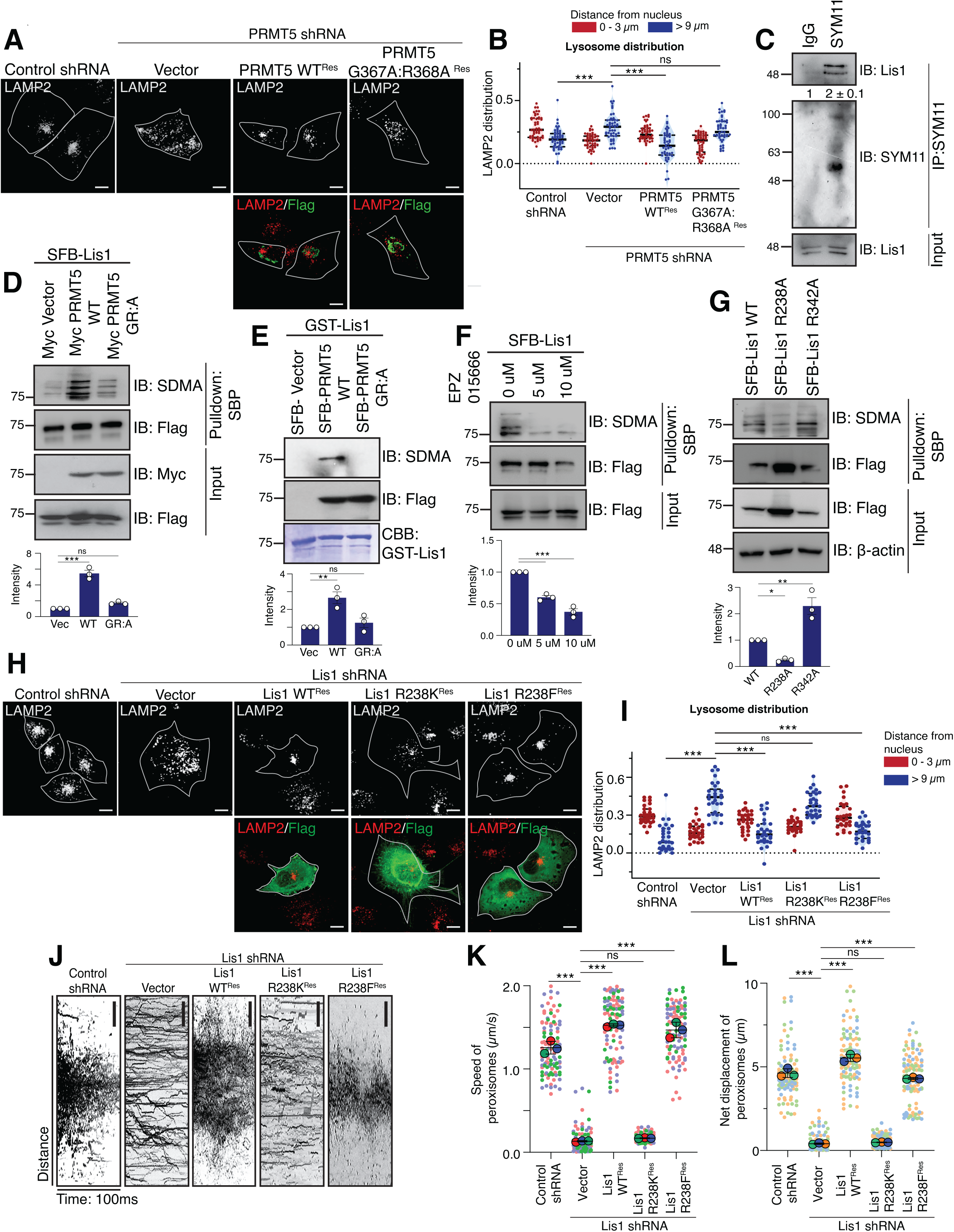
PRMT5 mediated methylation on Lis1 is necessary for Dynein function. **(A)** Control and PRMT5 depleted stable U2OS cells were transfected with shRNA resistant PRMT5 WT and its catalytic dead G367A:R368A plasmids. Cells were fixed and labelled with LAMP2 (lysosomes) antibody to visualize the lysosome distribution under confocal microscope. Scale bars: 10 µm. **(B)** Quantification of the distribution of LAMP2-positive compartments (n = 50 cells) across three biological replicates for the experiments shown in (A). Data are median ± interquartile range. (***p < 0.0001; ns not significant; two-tailed Student’s *t*-test). **(C)** HEK 293T cells were subjected to a denatured immunoprecipitation with SYM11 antibody. The eluates were immunoblotted with Lis1 antibody. The densitometric quantifications from three biological replicates are shown under each blot. **(D)** HEK 293T cells transfected with SFB-Lis1 plasmid along with Myc vector, Myc-PRMT5 WT and Myc-PRMT5 catalytic mutant (G367A:R368A). 24 hours after transfection, cells were lysed under denaturing conditions, and further incubated with streptavidin-sepharose beads. Methylated proteins were detected by immunoblotting with anti-symmetric dimethyl arginine antibody. The densitometric quantifications from three biological replicates are shown in the graph below. (ns = not significant, ***p < 0.0001; One way ANOVA followed by Dunnett’s multiple comparisons test). **(E)** Bacterially purified GST-Lis1 was incubated with PRMT5 WT and PRMT5 G367A:R368A purified from HEK 293T cells for *in vitro* methylation, along with S-adenosyl methionine (SAM). Methylated protein was visualized by immunoblotting with anti-symmetric dimethyl arginine antibody. CBB: Coomassie Brilliant Blue. The densitometric quantifications from three biological replicates are shown in the graph below. (ns = not significant, **p = 0.0054; One way ANOVA followed by Dunnett’s multiple comparisons test). **(F)** HEK 293T cells transfected with SFB-Lis1 plasmid were treated with EPZ015666 (PRMT5 inhibitor) at indicated concentrations for 12 hours. 24 hours after transfection, cells were lysed under denaturing conditions, and further incubated with streptavidin-sepharose beads. Methylated proteins were detected by immunoblotting with anti-symmetric dimethyl arginine antibody. The densitometric quantifications from three biological replicates are shown in the graph below. (***p < 0.0001; One way ANOVA followed by Dunnett’s multiple comparisons test). **(G)** HEK 293T cells transfected with SFB-Lis1 WT, R238A and R342A plasmid. 24 hours after transfection, cells were lysed under denaturing conditions, and further incubated with streptavidin-sepharose beads. Methylated proteins were detected by immunoblotting with anti-symmetric dimethyl arginine antibody. The densitometric quantifications from three biological replicates are shown in the graph below. (*p = 0.0476, **p < 0.0044; One way ANOVA followed by Dunnett’s multiple comparisons test). **(H)** Control and Lis1 depleted stable U2OS cells were transfected with shRNA resistant Lis1 WT, R238K, and R238F plasmids. Cells were fixed and labelled with LAMP2 (lysosomes) antibody to visualize the lysosome distribution under confocal microscope. Scale bars: 10 µm. **(I)** Quantification of the distribution of LAMP2-positive compartments (n = 30 cells) across three biological replicates for the experiments shown in (H). Data are median ± interquartile range. (\*\*\**p* < 0.0001, ns = not significant; two-tailed Student’s *t*-test). **(J)** Control and Lis1 depleted stable U2OS cells were transfected with Pex3-mEmerald-FKBP and FRB-BicD2-S along with shRNA resistant Lis1 WT, R238K, and R238F plasmids. After 24 hours of transfection, cells were treated with 1 uM Rapamycin just before live-cell imaging. Whole cell kymographs of peroxisomes representative of each condition are shown. Scale: 1 um. **(K)** Quantification of the speed of peroxisome in U2OS cells for the experiment shown in (J). Each color represents a replicate. Bigger circle is the mean of the data points represented as smaller circles from each biological replicate (n = 39, 35, 31 cells). Data are median ± interquartile range. (***p < 0.0001; ns not significant, two-tailed Student’s *t*-test). **(L)** Quantification of the net displacement of peroxisome in U2OS cells for the experiment shown in (J). Each color represents a replicate. Bigger circle is the mean of the data points represented as smaller circles from each biological replicate (n = 39, 35, 31 cells). Data are median ± interquartile range. (***p < 0.0001; ns not significant, two-tailed Student’s *t*-test).

### PRMT5 relieves dynein from CSN inhibition

Since, we found two new modifications important for dynein dependent cargo transport, we next wanted to test if these modifications occur on microtubule resident dynein or free cytoplasmic dynein complex. Firstly, Lis1 was found to be associated with moving dynein (Figure 7A, B and Video S8). Next, biochemical fractionation revealed that along with Lis1 and dynein, PRMT5 is also associated with cytoskeletal fraction (Figure 7C). Moreover, neddylation of DIC1 (Figure 7D) and methylation of Lis1 (Figure 7E) is exclusively found to occur on the microtubule associated Lis1-dynein complex. Together, this data suggests that PRMT5 complex is recruited to microtubule bound Lis1-dynein complex to mediate their modifications. We found both CSN and PRMT5 as important regulators of dynein movement that are recruited by Lis1. While CSN inhibits dynein movement, PRMT5 promotes dynein transport. To understand their interplay towards regulating dynein function, we hypothesized that PRMT5 mediated methylation on Lis1 acts as a positive switch by dislodging CSN from Lis1-dynein complex. Indeed, expression of PRMT5 led to reduction in binding of CSN (Figure 7F) with DIC1 whereas treatment of cells with PRMT5 inhibitor enhanced CSN1 binding with dynein complex (Figure 7G). In addition, in the presence of methylation mimic mutant of Lis1 (R238F) association of CSN1 and DIC1 is severely compromised (Figure7H), clearly demonstrating that PRMT5 mediated Lis1 methylation displaces CSN from Lis1-dynein complex.

**Figure 7:**
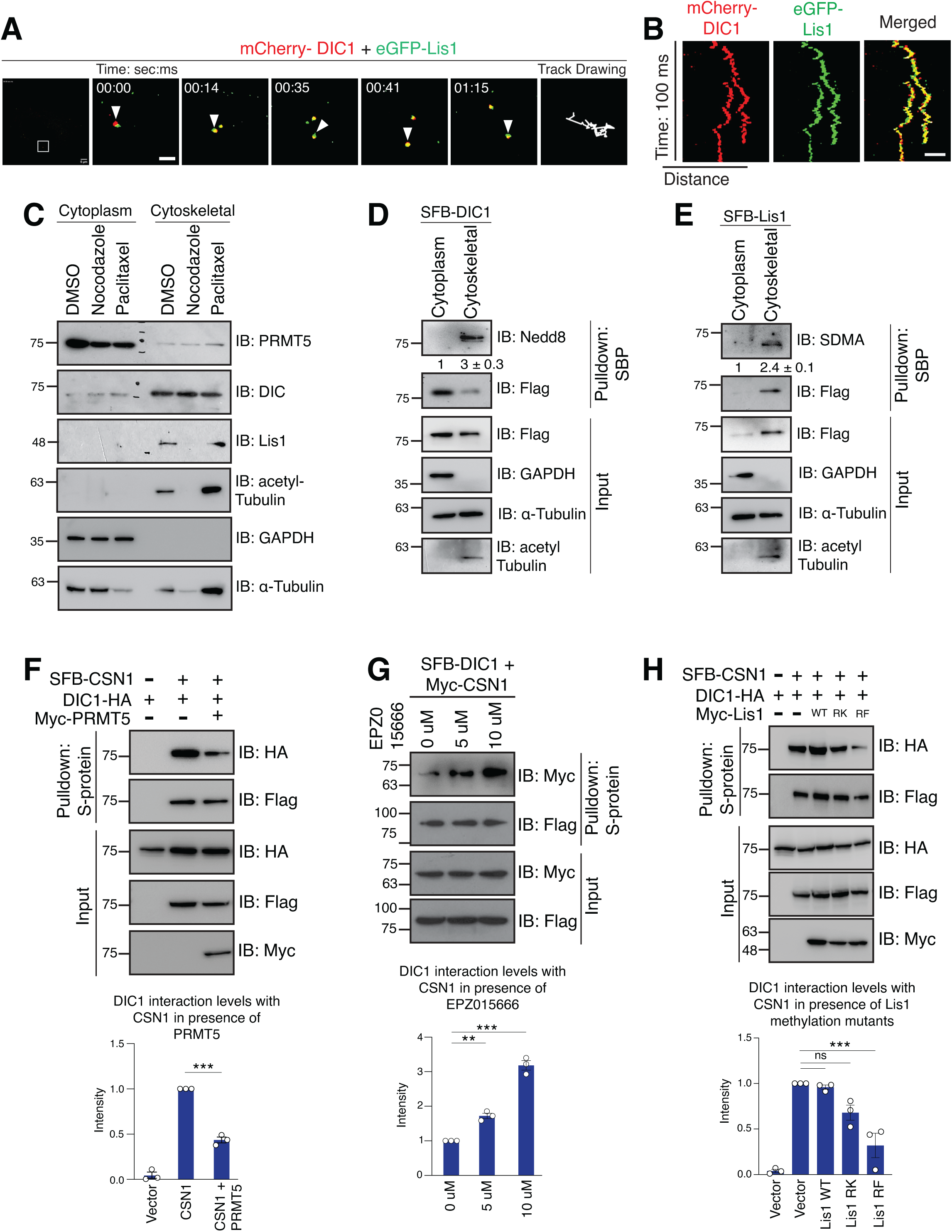
PRMT5 displaces CSN from dynein to facilitate neddylation of DIC1. **(A)** U2OS cells were transfected with mCherry-DIC1 and eGFP-Lis1 and were imaged using live-cell super-resolution microscopy with SIM module. The video stills, track drawing and the **(B)** respective kymographs of the movement are shown. Scale bars: 1 µm. **(C)** U2OS cells were fractionated into cytoplasmic and cytoskeletal fractions. The presence of proteins was detected by immunoblotting with indicated antibodies. **(D)** U2OS cells were transfected with SFB DIC1, and were fractionated into cytoplasmic and cytoskeletal fractions. The fractions were denatured and incubated with streptavidin-sepharose beads. Neddylated proteins were detected by immunoblotting with anti-Nedd8 antibody. The densitometric quantifications from three biological replicates are shown under each blot. **(E**) U2OS cells were transfected with SFB Lis1, and were fractionated into cytoplasmic and cytoskeletal fractions. The fractions were denatured and incubated with streptavidin-sepharose beads. Methylated proteins were detected by immunoblotting with anti-SDMA antibody. The densitometric quantifications from three biological replicates are shown under each blot. **(F)** HEK 293T cells were co-transfected with SFB-CSN1 and DIC1-HA. Their interaction was tested in the presence of Myc-PRMT5. Cells were then lysed, and incubated with S-protein beads. Interaction was detected by immunoblotting with anti-HA antibody. The densitometric quantifications from three biological replicates are shown in the graph below. (***p < 0.0001; Two tailed t test). **(G)** HEK 293T cells were co-transfected with SFB-DIC1 and Myc-CSN1 plasmids. Cells were treated with EPZ015666 (PRMT5 inhibitor) at indicated concentrations for 12 hours. Cells were then lysed, and incubated with S-protein beads. Interaction was detected by immunoblotting with anti-Myc antibody. The densitometric quantifications from three biological replicates are shown in the graph below. (**p = 0.0025, ***p < 0.0001; One way ANOVA followed by Dunnett’s multiple comparisons test). **(H)** HEK 293T cells were co-transfected with SFB-CSN1 and DIC1-HA. Their interaction was tested in the presence of Myc-Lis1 WT, Myc Lis1 R238K and Myc-Lis1 R238F. Cells were then lysed, and incubated with S-protein beads. Interaction was detected by immunoblotting with anti-HA antibody. Densitometric quantification from three biological replicates are shown in the graph. (ns = not significant, ***p = 0.0008; One way ANOVA followed by Dunnett’s multiple comparisons test).

Subsequently, in support of our hypothesis that PRMT5-Lis1 association displaces CSN to facilitate DIC1 neddylation, we found that loss of either Lis1 or PRMT5 led to reduction in DIC1 neddylation (Figure 8A, B). Further, expression of wild type PRMT5, but not a catalytic inactive mutant, enhanced DIC1 neddylation (Figure 8C). On contrary, inhibition of PRMT5 catalytic activity with a specific inhibitor reduced DIC1 neddylation (Figure 8D). On the other hand, we also tested the effect of neddylation inhibition on Lis1 methylation. We found that Lis1 methylation was in fact slightly enhanced upon neddylation inhibition (Figure S13E). This is possibly due to accumulation of methylated Lis1 species on dynein complex as dynein motor is unable to move in the absence of neddylation as shown above in figure 3.

**Figure 8:**
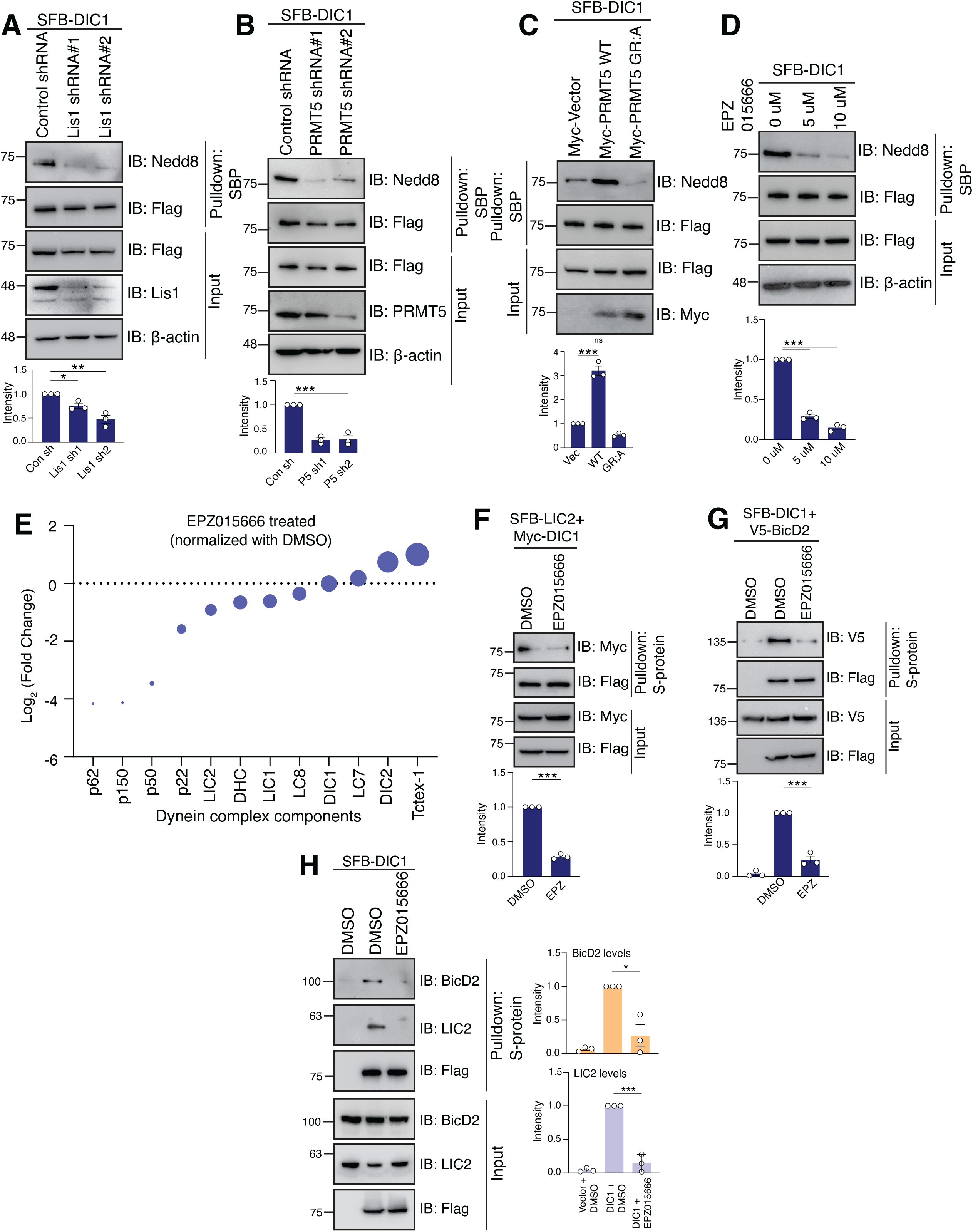
Loss of methylation leads to defective dynein complex assembly. **(A)** Control and Lis1 depleted stable HEK 293T cells were transfected with SFB-DIC1 plasmid. Cells were then lysed under denaturing conditions, and further incubated with streptavidin-sepharose beads. Neddylated proteins were detected by immunoblotting with anti-Nedd8 antibody. The densitometric quantifications from three biological replicates are shown in the graph below. (*p = 0.0412, **p = 0.0011; One way ANOVA followed by Dunnett’s multiple comparisons test). **(B)** Control and PRMT5 depleted stable HEK 293T cells were transfected with SFB-DIC1 plasmid. Cells were then lysed under denaturing conditions, and further incubated with streptavidin-sepharose beads. Neddylated proteins were detected by immunoblotting with anti-Nedd8 antibody. The densitometric quantifications from three biological replicates are shown in the graph below. (***p = 0.0001; One way ANOVA followed by Dunnett’s multiple comparisons test). **(C)** HEK 293T cells were co-transfected with SFB-DIC1 and Myc-PRMT5 WT and Myc-PRMT5 catalytic mutant (G367A:R368A) plasmids. Cells were lysed under denaturing conditions, and further incubated with streptavidin-sepharose beads. Neddylated proteins were detected by immunoblotting with anti-Nedd8 antibody. The densitometric quantifications from three biological replicates are shown in the graph below. (ns = not significant, ***p < 0.0001; One way ANOVA followed by Dunnett’s multiple comparisons test). **(D)** HEK 293T cells were transfected with SFB-DIC1 plasmid. Cells were treated with EPZ015666 (PRMT5 inhibitor) at indicated concentrations for 12 hours. Cells were lysed under denaturing conditions, and further incubated with streptavidin-sepharose beads. Neddylated proteins were detected by immunoblotting with anti-Nedd8 antibody. The densitometric quantifications from three biological replicates are shown in the graph below. (***p < 0.0001; One way ANOVA followed by Dunnett’s multiple comparisons test). **(E)** Quantitation of Log_2_ Fold change of the spectral counts of each protein identified in mass spectrometry in SFB-DIC1 purified from EPZ015666 treated cells and normalized with DMSO. **(F)** HEK 293T cells were co-transfected with SFB-LIC2 and Myc-DIC1 plasmids. Cells were treated with 5 uM EPZ015666 (PRMT5 inhibitor) or DMSO for 12 hours. Cells were lysed and further incubated with S-protein beads. Interaction was detected by immunoblotting with anti-Myc antibody. The densitometric quantifications from three biological replicates are shown in the graph below. (***p < 0.0001; Two tailed t test). **(G)** HEK 293T cells were co-transfected with SFB-DIC1 and V5-BicD2 plasmids. Cells were treated with 5 uM EPZ015666 (PRMT5 inhibitor) or DMSO for 12 hours. Cells were lysed and further incubated with S-protein beads. Interaction was detected by immunoblotting with anti-V5 antibody. The densitometric quantifications from three biological replicates are shown in the graph below. (***p < 0.0001; Two tailed t test). **(H)** HEK 293T cells were transfected with SFB-DIC1 plasmid. Cells were treated with 1 uM MLN4924 (NAE1 inhibitor) or DMSO for 12 hours. Cells were lysed and further incubated with S-protein beads. Interaction was detected by immunoblotting with anti-BicD2 and anti-LIC2 antibodies. Quantification from three biological replicates is shown in the graph. (*p = 0.0115, ***p = 0.0003; Two-tailed t-test).

Since we demonstrated that Lis1 methylation removes CSN and promotes dynein neddylation and at the same time neddylation was shown to be essential for active dynein assembly, we next tested the effect of Lis1 methylation on dynein assembly. In order to establish the defective assembly of dynein complex in cells upon loss of Lis1 methylation, we purified the complexes associated with SFB-tagged DIC1 proteins in the presence or absence of methylation inhibitor and identified the complexes using mass spectrometry. Treatment of cells with methylation inhibitor led to significant loss of spectral counts of dynactin complex and LIC2, with no changes in the abundance of dynein light chains (Figure 8E). Phenocopying the loss of neddylation, inhibition of PRMT5 in cells also led to defective binding of LIC2 and BicD2 (Figure 8F-H) with DIC1 with no changes in LIC1 binding (Figure S13F). Together, these findings establish that PRMT5-dependent methylation on Lis1 acts as a positive regulatory switch to remove CSN and drive the neddylation on dynein and subsequent assembly of dynein complex.

### Lis1 association with regulators is clinically relevant in lissencephaly

Lissencephaly (or smooth brain disorder) is caused by the failure of neuronal migration during early developmental stages ^16, 17, 58^. Loss of Lis1 or truncated Lis1 protein or mutations in Lis1 are some of the causes behind lissencephaly ^18, 19, 59^. Since we identified a functional connection between Lis1-CSN-PRMT5 association in dynein mediated cargo transport, we speculated that some of these pathogenic lissencephaly mutations may disrupt the association of Lis1-CSN or Lis1-PRMT5 complex and thus may contribute to disease. We searched for mutations of Lis1 identified in lissencephaly patients from HGMD (Human Gene Mutation Database) ^60^, where the gene lesions responsible for human inherited diseases are listed ^61^. As expected, we found several mutations in Lis1 such as F31S, H149R and G162S that displayed defective binding with dynein subunit, DIC1 (Figure 9A). But interestingly, we also found Lis1 mutations that are defective in binding with either CSN or PRMT5 irrespective of their ability to bind dynein. Mutations at G162S, S169P and D317H of Lis1 were defective in binding to CSN (Figure 9B), whereas H149R, G162S and D317H mutants were unable to bind to PRMT5 (Figure 9C). Functionally, while Lis1 wild type could rescue the lysosome position defects, DIC1 binding defective mutant (H149R) of Lis1 could not do so in Lis1 depleted cells (Figure 9D). Importantly, Lis1 S169P that retained the ability to bind to dynein and PRMT5 but defective in binding to CSN is hyperactive in dynein mediated transport (Figure 9D, E). On the other hand, D317H mutant with intact dynein binding but lost PRMT5 association is defective in rescuing the organelle position defects caused by Lis1 depletion. Together, this Lis1 clinical mutation data fully support our findings that Lis1 recruits CSN to put a brake on dynein movement and subsequently PRMT5 recruitment followed by methylation of Lis1 dislodges CSN to activate dynein. With this data, we provided possible mechanistic insights behind lissencephaly where either a positive or a negative regulation of dynein is lost, giving rise to a diseased phenotype.

**Figure 9:**
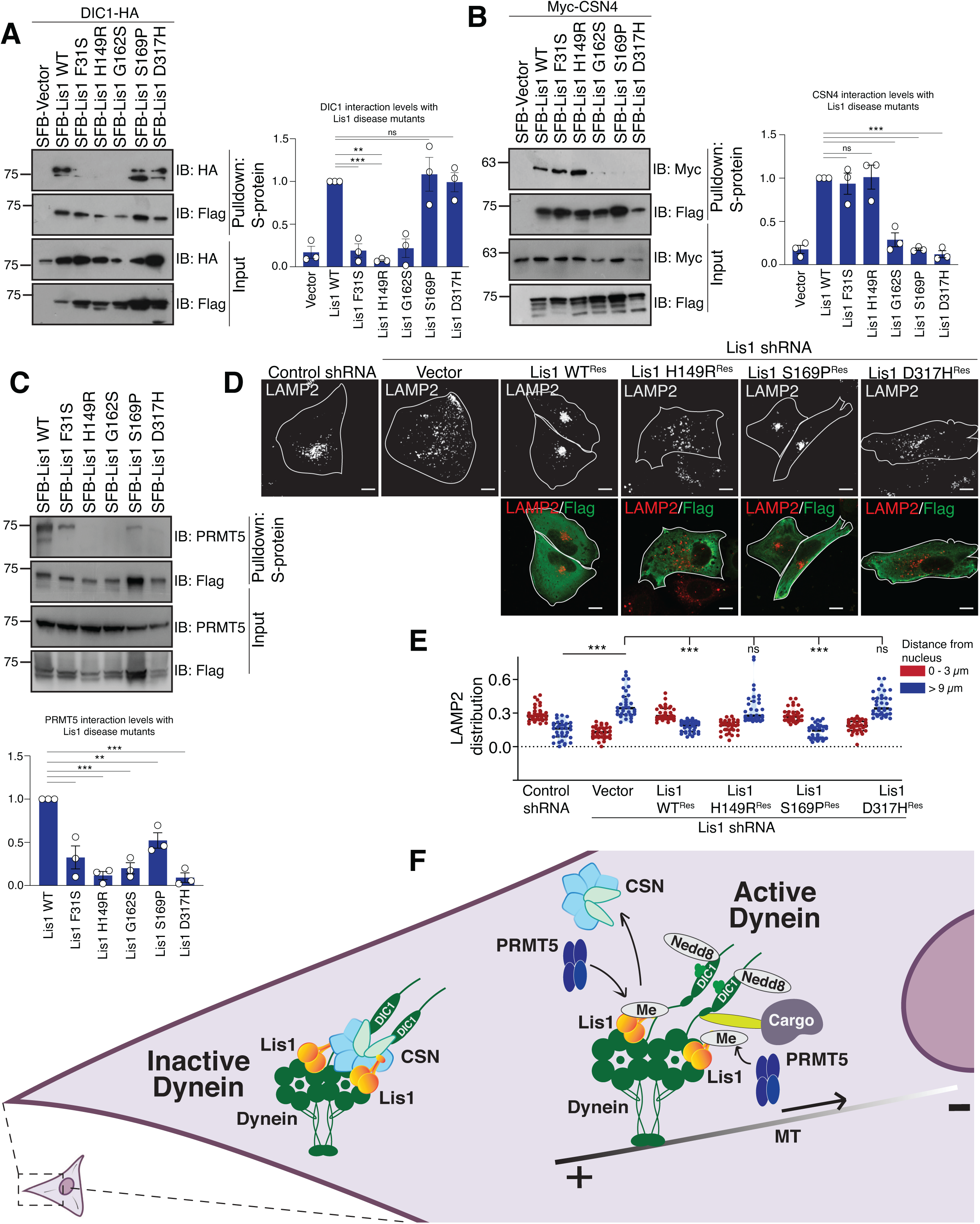
Lissencephaly disease mutations are defective in binding with CSN and PRMT5. **(A)** HEK 293T cells were co-transfected with SFB-Lis1 WT, SFB-Lis1 F31S, SFB-Lis1 H149R, SFB-Lis1 G162S, SFB-Lis1 S169P, and SFB-Lis1 D317H in combination with DIC1-HA plasmids. Cells were then lysed, and incubated with S-protein beads. Interaction was detected by immunoblotting with anti-HA antibody. Quantification from three biological replicates is shown in the graph. (***p = 0.0008, **p = 0.0011, ns = not significant; One way ANOVA followed by Dunnett’s multiple comparisons test). **(B)** HEK 293T cells were co-transfected with SFB-Lis1 WT, SFB-Lis1 F31S, SFB-Lis1 H149R, SFB-Lis1 G162S, SFB-Lis1 S169P, and SFB-Lis1 D317H in combination with Myc-CSN4 plasmids. Cells were then lysed, and incubated with S-protein beads. Interaction was detected by immunoblotting with anti-Myc antibody. Quantification from three biological replicates is shown in the graph. (***p = 0.0003, ns = not significant; One way ANOVA followed by Dunnett’s multiple comparisons test). **(C)** HEK 293T cells were transfected with SFB-Lis1 WT, SFB-Lis1 F31S, SFB-Lis1 H149R, SFB-Lis1 G162S, SFB-Lis1 S169P, and SFB-Lis1 D317H plasmids. Cells were then lysed, and incubated with S-protein beads. Interaction was detected by immunoblotting with anti-PRMT5 antibody. Quantification from three biological replicates is shown in the graph. (***p = 0.0002, **p = 0.0034; One way ANOVA followed by Dunnett’s multiple comparisons test). **(D)** Control and Lis1 depleted U2OS cells were transfected with shRNA resistant SFB-Lis1 WT, SFB-Lis1 H149R, SFB-Lis1 S169P, and SFB-Lis1 D317H plasmids. Cells were fixed, and labelled with LAMP2 (lysosomes) antibody to visualize their distribution under confocal microscope. Scale bars: 10 µm. **(E)** Quantification of the distribution of LAMP2-positive compartments (n = 35 cells) across three biological replicates for the experiments shown in (D) Data are median ± interquartile range. (***p < 0.0001; **p 0.0048; ns not significant; two-tailed Student’s *t*-test). **(F)** A model for regulation of dynein by Lis1 where CSN keeps dynein inactive by deneddylating it, while PRMT5 activates dynein by methylating Lis1, which in turn, facilitates DIC1 neddylation, and thus makes the dynein motor active.

## Discussion

In conclusion, we identified two regulators, CSN and PRMT5 complexes, that contribute to opposing behaviour of Lis1 on dynein. We found that Lis1 recruits CSN to deneddylate dynein and keep it inactive. For activation of dynein, PRMT5 methylates Lis1 and displaces CSN, which facilitates the neddylation of DIC1, thereby marking the activation and movement of dynein (model shown in Figure 9F). Dynein undergoes numerous conformational changes in the cell. It takes a phi conformation when it is auto-inhibited, and an open conformation when it is active ^3, 34^. Even when it is active, it undergoes several conformational changes upon loading to microtubules, binding with cargoes and dynactin ^6, 62, 63^. Each of these events may perhaps be governed by modification-based switches. However, only a handful of studies showed post-translational modifications (such as phosphorylation and pyrophosphorylation) on dynein ^64–66^. Similarly, Lis1 has been shown to regulate dynein in various ways, but PTMs on Lis1 are limited. Lis1 was shown to be phosphorylated ^67^, but the underlying mechanism, and the fate of the modified protein was unknown. Here, we have reported two new modifications regulating the activation of Lis1-dynein function in cells.

Several interesting points arise from our work. Our study concurs with other studies where Lis1 was shown to stall dynein for longer periods of time and slowing down dynein ^25, 68^. Our work highlights the involvement of a deneddylase complex CSN, which deneddylates dynein and keeps it inactive. Lis1 governing this association supports earlier studies that showed Lis1 as a negative regulator of dynein movement ^29^. Nevertheless, neddylation has largely been studied in context with ubiquitin systems ^69, 70^; this is the first report where this modification has been explored in context with dynein and its movement. Additionally, dynein intermediate chain 1 (DIC1) is the first substrate of CSN outside the realms of ubiquitin biology, which opens new possibilities of regulation for this complex. It is also noteworthy, that most of the PTMs on dynein reported hitherto cause impaired functions ^65^, however, neddylation on DIC1 shown in this study is necessary not only for its assembly, but also for the movement of dynein.

On the other hand, our work also supports the studies where Lis1 was demonstrated to form active dynein complexes, and responsible for processive dynein movements ^10, 12, 14, 31–33^. This study underscores the involvement of a new regulator, PRMT5 methyltransferase, which methylates Lis1 and promotes dynein movement. Not only that, PRMT5 methylates Lis1, which helps in relieving dynein from the inhibition of CSN, and facilitates the neddylation of DIC1. This crosstalk of post-translational modifications are the hallmarks for a proper assembly of dynein complex, and hence leads to movement of dynein. Thus, this study reports a new duo of methylation facilitating neddylation to regulate dynein movement, which opens new directions for exploring modifications on this interface.

Our findings reveal a dual regulatory system where Lis1 interacts with both the CSN and PRMT5, influencing dynein’s function through distinct mechanisms, possibly explaining the opposing roles of Lis1 on dynein ^14, 40, 41^. While, we denoted PRMT5 as a switch-ON mechanism for Lis1-dynein activation, it would be interesting in future studies to further explore when and how demethylation of Lis1 occurs and the identity of the demethylating enzymatic machinery. It is worth noting that we observed co-localization of CSN with DIC1 near perinuclear space in addition to cell periphery suggesting a possible re-recruitment of CSN to inactivate dynein. Therefore, identifying the demethylases and their possible connection in re-recruitment of CSN would further help in understanding the complete active-inactive cycles of dynein. On the disease front, lissencephaly is caused by failure of neuronal migration during early developmental processes ^18^. The major reason behind this disease are the mutations on Lis1 ^19, 59^, with limited molecular mechanism. This study focuses on the key events that gets lopsided in lissencephaly, such as defective associations with CSN and PRMT5 that may give rise to uncoordinated movement of dynein, leading to disease phenotypes. The implications of these interactions extend to understanding and potentially addressing disorders like lissencephaly, where disruptions in this finely-tuned system have significant developmental consequences. Modulating the interaction between Lis1 and its regulatory partners or correcting specific post-translational modifications could offer potential new strategies for treating lissencephaly and other neurodevelopmental disorders associated with dynein dysfunction.

## Materials and Methods

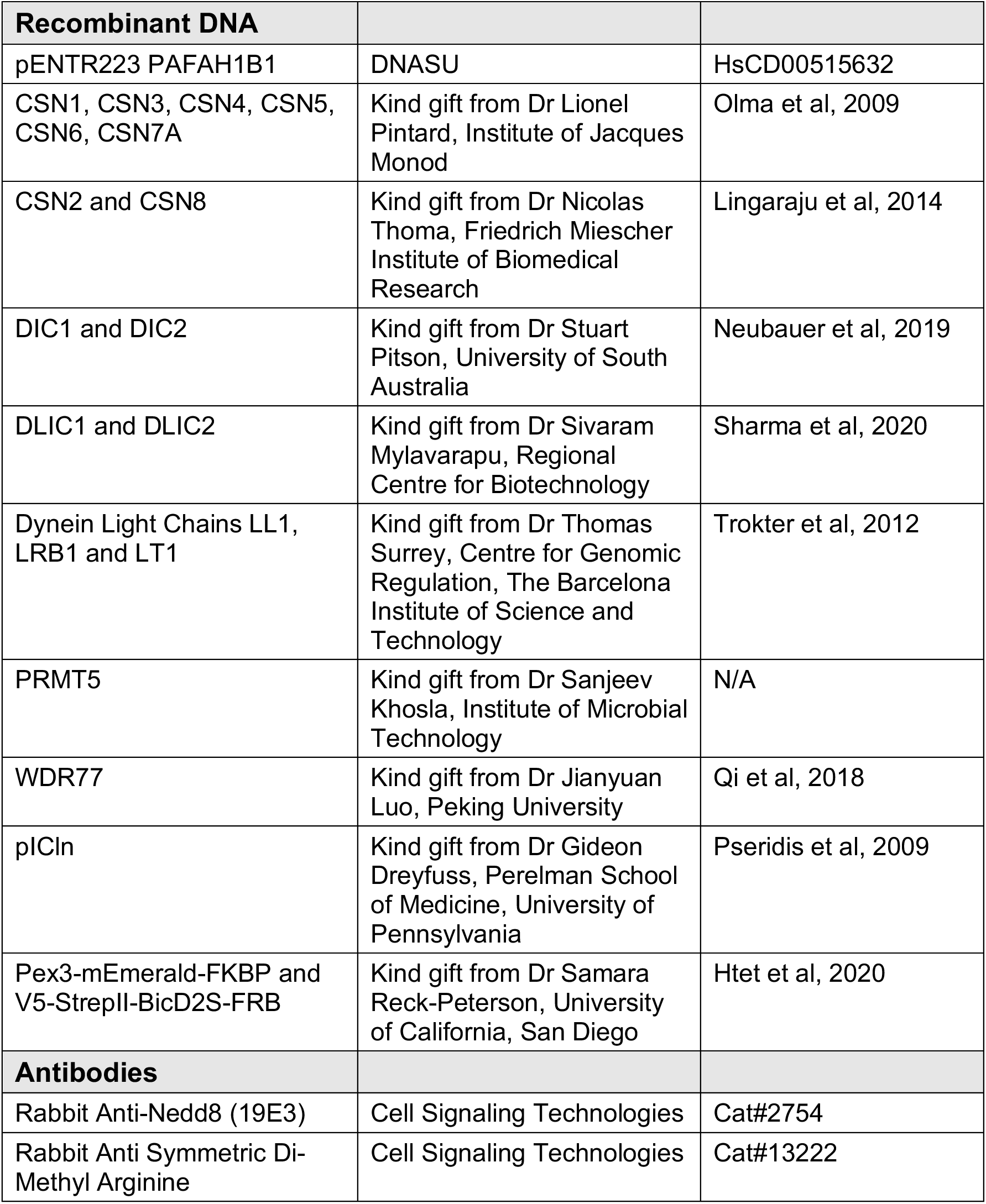
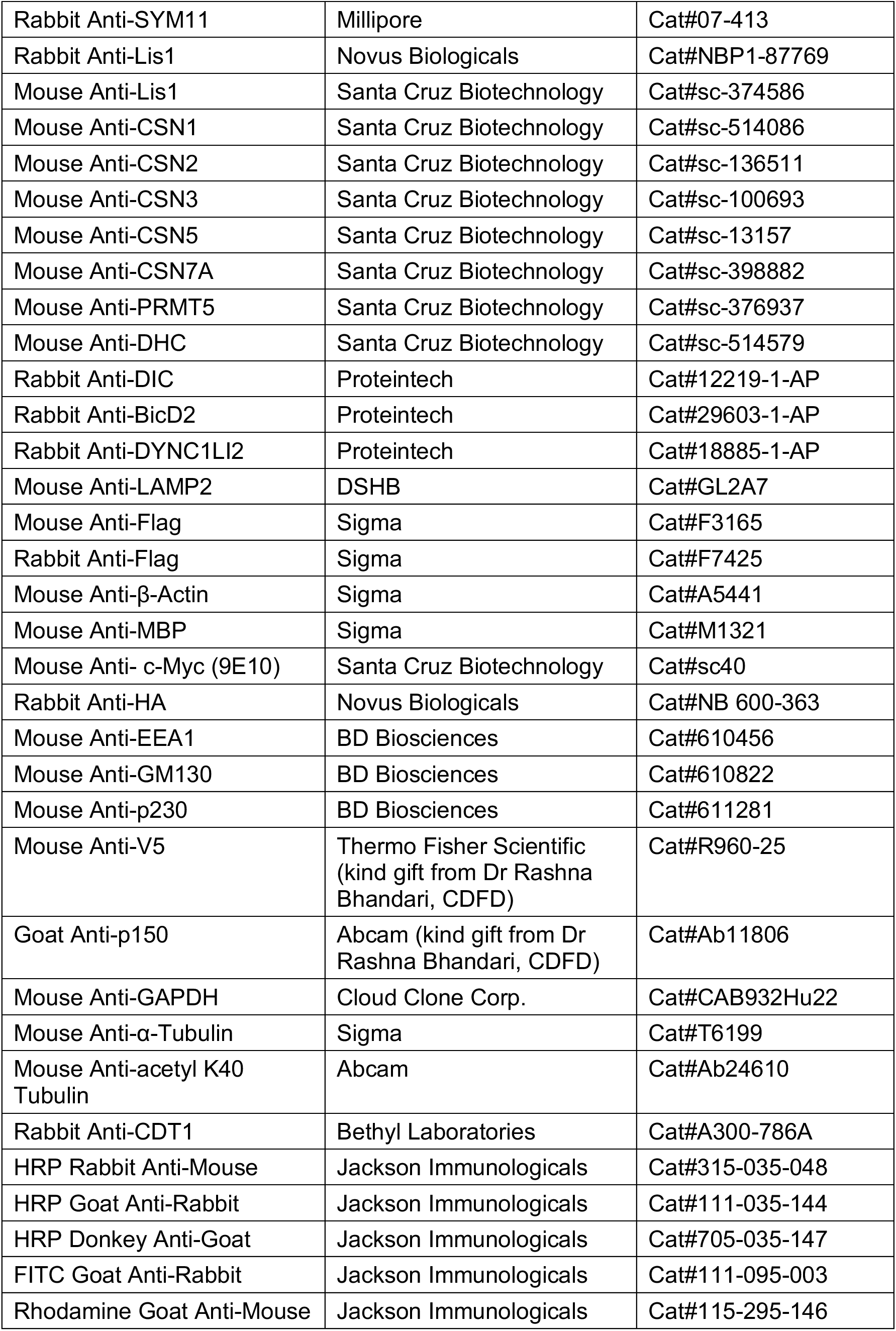
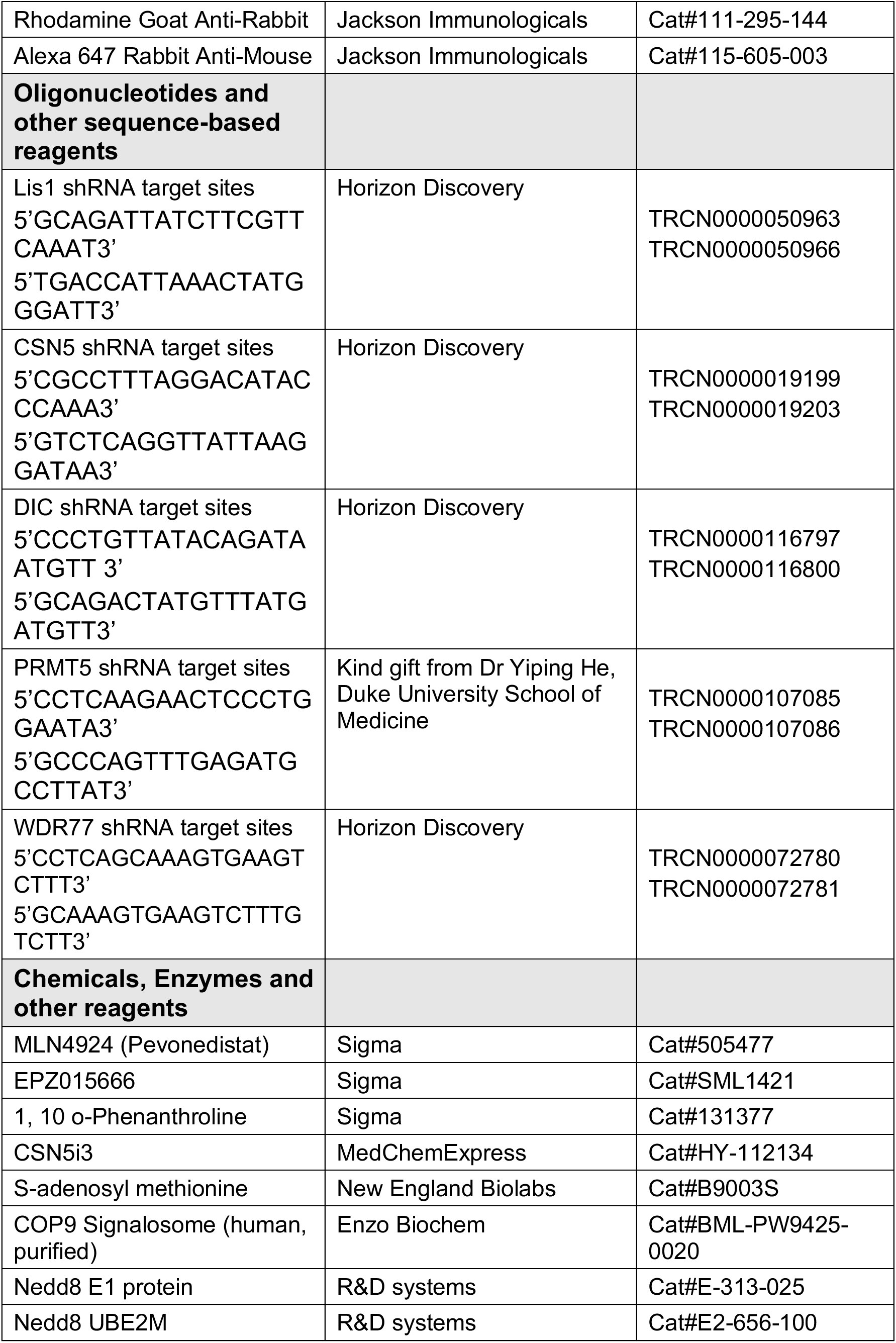
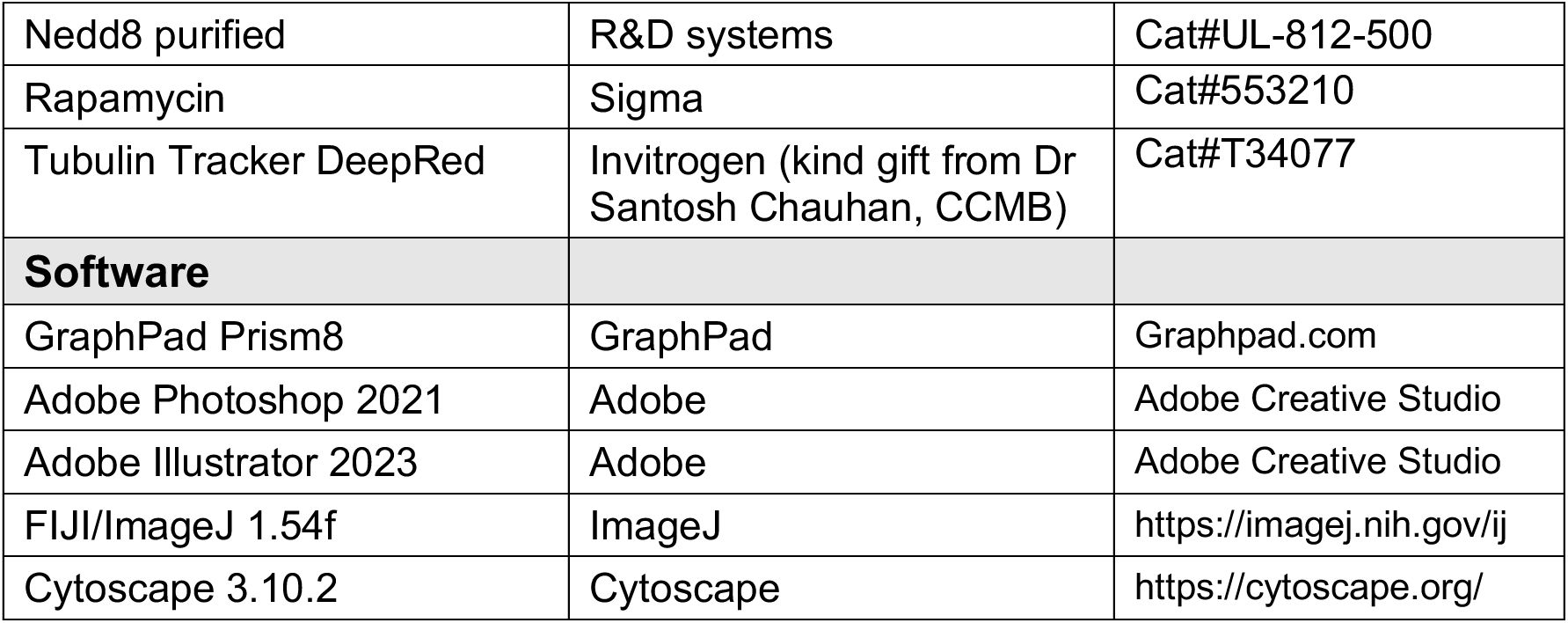

### Plasmids

The plasmids used in this study were cloned using Gateway Cloning (Invitrogen). Expression plasmids were moved into (S-protein/Flag/Streptavidin-binding-protein) SFB, Myc, MBP, His and mCherry-tagged vectors for expression. All clones were verified by Sanger sequencing. All the point mutations were generated by PCR-based site directed mutagenesis and further cloned into Gateway vectors.

### Transfections and Lentiviral transduction

HEK293T and BOSC23 cells were grown in RPMI, while U2OS and IMR32 cells were grown in DMEM, containing 10% FBS and 1% penicillin and streptomycin. All cell lines were purchased from American Type Culture Collection (ATCC), which were tested and authenticated by the cell bank using their standard short tandem repeat (STR)-based techniques.

Cell lines were transfected with various plasmids using PEI (Polysciences) according to the manufacturer’s protocol. Briefly, the plasmid was diluted in serum-free medium mixed with PEI (1 µg/µl) in 1:3 ratios. After incubating the DNA and PEI mixture at room temperature (RT) for 20 min, the complexes were added to cells to allow the transfection of plasmid.

Lentivirus-based respective shRNA coding plasmids were transfected transiently using PEI (Invitrogen) in BOSC23 packaging cells along with packaging vectors (psPAX2 and pMD2.G). At 48 h post transfection, the viral medium was collected, filtered through a 0.45 µM filter and added to the target cells along with polybrene (8 µg/ml). 48 h post infection, cells were collected and processed for various assays and immunoblotting was performed with the specific antibodies to check the efficiency of knockdown.

### Immunoprecipitation and pull-down assays

Cells were lysed with NETN buffer (20 mM Tris/HCl, pH 8.0, 100 mM NaCl, 1 mM EDTA, and 0.5% Nonidet P-40) containing 1 mg·mL^−1^ of each Pepstatin A, Aprotinin, and 100 mM PMSF (phenylmethylsulphonyl fluoride) on ice for 20 minutes. The whole-cell lysates were incubated with either Protein G Sepharose beads conjugated with respective antibody, or Streptavidin– Sepharose beads (Amersham Biosciences), or S-protein–agarose beads (Novagen) for 2 h at 4 °C. The protein complexes were then washed with NETN buffer and subjected to SDS-PAGE. Immunoblotting was performed using standard protocols.

For detecting neddylation and methylation in cells, denaturing immunoprecipitation was performed by boiling the HEK 293T cells in denaturing lysis buffer (50 mM Tris-HCl pH 7.5, 100 mM β-mercaptoethanol, 1% SDS and 5 mM EDTA) for 10 min followed by sonication. The SDS concentration was adjusted to 0.1% by adding 1× NETN buffer and incubated on ice for 20 min. The whole-cell lysates obtained by centrifugation were incubated with streptavidin– sepharose for overnight at 4°C. The immunocomplexes were then washed with NETN buffer and subjected to SDS–PAGE. For detecting neddylation, 2 uM of 1,10 OPT was added to the lysis buffer.

### Tandem Affinity Purification

Lis1-associated proteins were isolated by using tandem affinity purification as described before^71^. Briefly, HEK 293T cells expressing SFB-Lis1 were lysed with NETN lysis buffer containing protease inhibitors on ice for 20 min. The cell lysates were added on to streptavidin–sepharose beads and incubated for 2 h at 4 °C. Then, beads were washed with NETN lysis buffer and the associated proteins were eluted using 2 mg·mL^−1^ Biotin (Sigma) for 2 h at 4 °C. The eluates from the first step of purification were then incubated with S-protein–agarose beads (Novagen) for 2 h at 4 °C. After clearing the unbound proteins by washing, the proteins were eluted by boiling in SDS-loading buffer for 5 min at 95 °C. Eluted protein lysate was loaded on SDS/PAGE. The associated proteins were identified by in-gel trypsin digestion followed by LC-MS/MS analysis at Taplin Biological Mass Spectrometry Facility (Harvard University). SFB-tagged GFP was used as a control for purification. For filtering the interactome data, we compared the dataset of SFB-Lis1 with SFB-GFP and only the unique proteins were attributed as the interacting partners of Lis1. For dynein complex assembly dataset, the spectral counts of each protein were first normalized with DIC1 within the sample. Later the common interactors were compared against the control dataset and the fold change was obtained. The log_2_ (fold change) was plotted for each interactor using the GraphPad Prism 8.0.

### Data availability

The mass spectrometry proteomics data have been deposited to the ProteomeXchange Consortium via the PRIDE ^72^ partner repository with the dataset identifier PXD058912 and 10.6019/PXD058912.

Submission details:

Project Name: Lis1 dynein interactome dataset

Project accession: PXD058912

Project DOI: 10.6019/PXD058912

Reviewer access details

Reviewer can access the dataset by logging in to the PRIDE website using the following account details:

Username

Password: 8ItrNizlNsWJ

### Recombinant protein purification and enzymatic assays

GST, MBP and His-tagged respective proteins were transformed into *Escherichia coli* BL21 (DE3) cells. Cultures were grown to an optical density (OD) at 600 nm of ∼0.6 and induced with 0.5 mM isopropyl β-D-1-thiogalactopyranoside (IPTG) at 18°C for overnight. The cell pellets were lysed in lysis buffer (50 mM Tris-HCl pH 7.5, 150 mM NaCl, and 0.01% NP–40 IGEPAL), and 20 mM Imidazole pH 8.0 with 0.1% NP-40 IGEPAL (for His-tagged proteins), and protease inhibitors (Aprotinin, Pepstatin, PMSF) and sonicated. Cell lysates were pulled down with Dextrin–Sepharose or Glutathione Sepharose or Nickel NTA beads for 2 h at 4°C. Then, beads were washed five times with wash buffer (50 mM Tris-HCl pH 7.5, 300 mM NaCl, 0.01% NP– 40 IGEPAL, 1 mM DTT and protease inhibitors) or and bound proteins were eluted with the elution buffer containing 20 mM Tris-HCl pH 7.5, 200 mM NaCl, 1 mM EDTA and 10 mM maltose, or 50 mM Tris-HCl pH 8, 150 mM NaCl and 10 mM reduced glutathione, or 50 mM Tris-HCl pH 7.5, 150 mM NaCl, 300 mM Imidazole pH 8.0 with 0.1% NP-40 Igepal. The proteins after purification were used for further enzymatic assay.

For *in vitro* methylation assay, GST-Lis1 was purified on beads (as mentioned above), followed by incubation with purified SFB-PRMT5 from HEK 293T cells in methylation buffer (50 mM Tris pH 8.8, 5 mM MgCl2, 4 mM DTT) and 2 mM S-adenosylmethionine (SAM). The reaction was incubated at 30°C for 1 h. After incubation, reaction was stopped by adding the SDS loading buffer and subjected to SDS-PAGE.

For *in vitro* neddylation assay, 1 µM NAE1, 2 µM UBE2M, and 15 µM Nedd8 were added to reactions with neddylation buffer (50 mM Tris, pH 7.6, 150 mM NaCl, 10 mM MgCl2, 1 mM DTT and 5 mM ATP). Reactions were incubated at 30°C for 1 h, terminated with SDS loading buffer and analysed by SDS-PAGE. For deneddylation reactions, the initial neddylation reaction was washed with neddylation buffer twice, and incubated for another 1 h at 30°C with purified CSN complex (0.3 µg), a modified protocol ^73^.

### Cytoskeletal fractionation

The fractionation was performed using Semi-retentive cytoskeletal fractionation (SERCYF) as described in ^74^. Briefly, cells were washed with pre-warmed PIPES-EGTA-MgCl_2_ (PEM) Buffer (100 mM PIPES-NaOH (pH 6.8), 1 mM EGTA, 2 mM MgCl_2_). For extraction of cytoplasmic fraction, pre-warmed PEMTT buffer (0.2% Triton X-100, 100 nM paclitaxel in PEM) was added on top of the cells and incubated at 37°C for 1 min without shaking. This fraction was collected in a micro-centrifuge tube. The cells were again rinsed with PEMTT buffer to collect the residual cytoplasmic fraction and added to the previous tube, to which 2X SDS buffer was added. To collect cytoskeletal fraction, an equal volume of 2X SDS buffer was added on the plate, and the fraction was collected by scraping the cells. The samples were further denatured by boiling at 95°C for 5 min. The cells used for this experiment were U2OS cells, as they are adherent and do not come out easily, compared to the HEK 293T cells. GAPDH was used as the cytoplasmic marker, while acetyl α-tubulin was used as the cytoskeletal marker for the respective fractions.

### Immunofluorescence and Live cell microscopy

Cells grown on coverslips were fixed with 4% paraformaldehyde for 15 min at 37°C and permeabilized with 0.2% Triton X-100 for 10 min at RT. For visualization of microtubules, cells were first fixed with 4% paraformaldehyde for 15 min at 37°C, washed with 1× PBS thrice, and then fixed with chilled (kept at −20°C) methanol for 4 min. This step eliminates the need for Triton X-100 mediated permeabilization, so after washing the traces of methanol with 1× PBS, the samples can be directly proceeded for blocking. Samples were blocked with 5% BSA at RT for 30 min and incubation with primary antibodies for overnight at 4°C.

After incubation, cells were washed three times with 1× PBS and then incubated with FITC, Rhodamine, or Alexa Fluor 647 conjugated secondary antibodies at RT for 60 min followed by three washes with 1× PBS. Cells were washed with 1× PBS, and coverslips were mounted with glycerol containing paraphenylenediamine and imaged using a confocal microscope (Leica TCS SP8) or Carl Zeiss ELYRA Super-Resolution Microscope with Plan apochromat 63X oil immersion lens with N.A 1.46. For super resolution images, a lattice pattern structured and 15 phases shifted raw images were acquired for every Z plane with a slice size of 110 nm. The raw images were acquired with Zen Black and processed through Structured Illumination Microscopy (SIM) module, quantified using Zen blue and Fiji software.

For live cell imaging of peroxisome motility assay, cells were seeded in 20 mm glass bottom dishes (ThermoFisher) for 24 hours. Cells were then transfected with eGFP or mCherrry-tagged plasmids using PEI according to the manufacturer’s protocol. Cells were imaged after 30-36 hours after transfection. For peroxisome motility assay, 24 hours after transfection, media was replenished and 1 uM Rapamycin was added 20 min before imaging using Carl Zeiss ELYRA Super-Resolution Microscope. Time lapse imaging was performed with exposure for 50 ms, at an interval of 1 ms for 100 cycles. Raw videos were captured and then subjected to SIM processing and analysed using Fiji. The videos run at 5 frames per second.

### Image analysis and quantification

Western blot quantifications were performed using Fiji. Briefly, a rectangular ROI was drawn around the band of interest, which results in a curve, selecting the area under the curve gives the intensity of the band. These measurements were exported to excel, and normalized with respect with the control samples. For quantitation of the blots in interaction studies, the immunoprecipitated fractions were first normalized with their respective inputs, and then compared with the amount of protein in pull-down. Any blobs on or around the band of interest are also taken into consideration in the densitometric quantifications.

Quantification of the distribution of lysosomes and endosomes based on LAMP2 and EEA1 signal intensity was performed as described in ^75^. Briefly, a boundary was drawn along the periphery of each selected cell using the freehand selection tool in Fiji. To avoid the bias of signals from the neighbouring cells, ‘clear outside’ function was used. Next, an ROI was drawn around the nucleus, and LAMP2/EEA1 signal intensity was measured for that section. The same ROI was then incremented by 3 µm till the cell periphery, and LAMP2/EEA1 intensity was measured for each incremented ROI. Finally, LAMP2/EEA1 intensity was calculated for perinuclear (0–3 µm; by subtracting the intensity of the first ROI from second) and periphery (>9 µm; by subtracting the intensity of the fourth ROI from total cell intensity) region of cell. LAMP2 distribution was plotted by dividing each section’s intensity (perinuclear and periphery) with total LAMP2 intensity of each cell.

The golgi morphology was described by observing visual changes in its distribution for each cell across the samples. The standard for each category of ribbon, broken ribbon and scatter phenotypes has been shown in relevant studies ^49^.

The colocalization analysis was done using inbuilt plugin Coloc in Zen Black or using JACoP plugin of Fiji software and the corresponding Pearson’s correlation coefficient was determined. Signal intensity within the cell was considered for colocalization. Briefly, multiple ROIs (usually 3 – 4) were selected within a cell, which were averaged out later for each cell and their respective data points for each cell across biological replicates were plotted on the graph.

For tracking single-particle peroxisomes motility in cells, we used TrackMate plugin of Fiji. The parameters used are as follows: Vesicle diameter, 1 µm; Detector, DoG; Initial thresholding, none; Tracker, Simple LAP tracker; Linking max distance, 2 µm; Gap-closing max distance, 2 µm; Gap-closing max frame gap, 2; Filters, none. This data shows the movement of each particle from a particular x,y coordinate to another, the duration for which it moved, and number of stops it made during the movement. Considering these points, we imaged around 15-20 cells in each replicate and picked at least 5 moving peroxisomes from each cell. The tracks and spots data were exported as csv files and respective speed and displacement were plotted on graph using GraphPad Prism 8.0.

For assessing dynein movement in cells, we measured the movement of dynein particles using TrackMate with the same setting as described above. After assessing the motility patterns of dynein particles, we assigned particles moving above 0.1 µm/sec as moving dynein, while the rest as non-moving dynein. Their respective speeds in different samples were plotted using GraphPad prism 8.0.

The kymographs were made using Kymograph Builder plugin of Fiji. The track drawing of live cell videos was performed using MTrackJ plugin in Fiji. Briefly, the point to be tracked was selected, then ‘add’ button activates multipoint tool which draws points on the track. After manually clicking on the moving particle within each frame, the track drawing is formed, which can be exported either as a video or an image format.

### Statistics

All the graphs in this study represent either mean ± SD, or mean ± SE, or median ± interquartile range and *p* values were calculated using two-tailed Student’s *t*-test, One-way ANOVA, or Two-way ANOVA (GraphPad Prism 8.0). Differences between groups were considered statistically significant for *p* values < 0.05. All immunofluorescence images, were analyzed from at least three independent experiments or at least eight cells.

## Acknowledgments

This work was supported by DBT grant (BT/PR44301/BRB/10/2003/2021 to S.M), DBT National Bioscience Award for Career Development (BT/HRD/NBA/39/08/2018-19 to S.M) and CDFD core funds. D.G. acknowledges the support of research fellowship from the Council of Scientific and Industrial Research (CSIR). We thank Gaurav Kumar Khunger and Dr Amit Tuli for their valuable insights in imaging data analysis and quantification. We thank all members of LCDCS for their critical inputs. The authors thank Nanci Rani for providing technical assistance.

## Conflict of interest

The authors declare that they have no conflict of interest.

## Author contributions

SM conceptualized and managed the project. SM and DG designed the experiments, analysed the data and wrote the manuscript. DG performed all the experiments.

## Notes

### Competing Interest Statement

The authors have declared no competing interest.

## References

1. Canty, J.T. & Yildiz, A. Activation and Regulation of Cytoplasmic Dynein. Trends Biochem Sci 45, 440–453 (2020).

2. Paschal, B.M. & Vallee, R.B. Retrograde transport by the microtubule-associated protein MAP 1C. Nature 330, 181–183 (1987).

3. Canty, J.T., Tan, R., Kusakci, E., Fernandes, J. & Yildiz, A. Structure and Mechanics of Dynein Motors. Annu Rev Biophys 50, 549–574 (2021).

4. King, S.J., Bonilla, M., Rodgers, M.E. & Schroer, T.A. Subunit organization in cytoplasmic dynein subcomplexes. Protein Sci 11, 1239–1250 (2002).

5. Reck-Peterson, S.L., Redwine, W.B., Vale, R.D. & Carter, A.P. The cytoplasmic dynein transport machinery and its many cargoes. Nat Rev Mol Cell Biol 19, 382–398 (2018).

6. Roberts, A.J., Kon, T., Knight, P.J., Sutoh, K. & Burgess, S.A. Functions and mechanics of dynein motor proteins. Nat Rev Mol Cell Biol 14, 713–726 (2013).

7. Olenick, M.A. & Holzbaur, E.L.F. Dynein activators and adaptors at a glance. J Cell Sci 132 (2019).

8. Carter, A.P., Diamant, A.G. & Urnavicius, L. How dynein and dynactin transport cargos: a structural perspective. Curr Opin Struct Biol 37, 62–70 (2016).

9. Hoogenraad, C.C. et al. Bicaudal D induces selective dynein-mediated microtubule minus end-directed transport. EMBO J 22, 6004–6015 (2003).

10. Egan, M.J., Tan, K. & Reck-Peterson, S.L. Lis1 is an initiation factor for dynein-driven organelle transport. J Cell Biol 197, 971–982 (2012).

11. Markus, S.M., Marzo, M.G. & McKenney, R.J. New insights into the mechanism of dynein motor regulation by lissencephaly-1. Elife 9 (2020).

12. Singh, K. et al. Molecular mechanism of dynein-dynactin complex assembly by LIS1. Science 383, eadk8544 (2024).

13. Reimer, J.M., DeSantis, M.E., Reck-Peterson, S.L. & Leschziner, A.E. Structures of human dynein in complex with the lissencephaly 1 protein, LIS1. Elife 12 (2023).

14. Gutierrez, P.A., Ackermann, B.E., Vershinin, M. & McKenney, R.J. Differential effects of the dynein-regulatory factor Lissencephaly-1 on processive dynein-dynactin motility. J Biol Chem 292, 12245–12255 (2017).

15. Toropova, K. et al. Lis1 regulates dynein by sterically blocking its mechanochemical cycle. Elife 3 (2014).

16. Fry, A.E., Cushion, T.D. & Pilz, D.T. The genetics of lissencephaly. Am J Med Genet C Semin Med Genet 166C, 198–210 (2014).

17. Mochida, G.H. Genetics and biology of microcephaly and lissencephaly. Semin Pediatr Neurol 16, 120–126 (2009).

18. Dobyns, W.B., Reiner, O., Carrozzo, R. & Ledbetter, D.H. Lissencephaly. A human brain malformation associated with deletion of the LIS1 gene located at chromosome 17p13. JAMA 270, 2838–2842 (1993).

19. Wynshaw-Boris, A. Lissencephaly and LIS1: insights into the molecular mechanisms of neuronal migration and development. Clin Genet 72, 296–304 (2007).

20. Splinter, D. et al. Bicaudal D2, dynein, and kinesin-1 associate with nuclear pore complexes and regulate centrosome and nuclear positioning during mitotic entry. PLoS Biol 8, e1000350 (2010).

21. Raaijmakers, J.A., Tanenbaum, M.E. & Medema, R.H. Systematic dissection of dynein regulators in mitosis. J Cell Biol 201, 201–215 (2013).

22. Markus, S.M., Punch, J.J. & Lee, W.L. Motor- and tail-dependent targeting of dynein to microtubule plus ends and the cell cortex. Curr Biol 19, 196–205 (2009).

23. Faulkner, N.E. et al. A role for the lissencephaly gene LIS1 in mitosis and cytoplasmic dynein function. Nat Cell Biol 2, 784–791 (2000).

24. Moon, H.M. et al. LIS1 controls mitosis and mitotic spindle organization via the LIS1-NDEL1-dynein complex. Hum Mol Genet 23, 449–466 (2014).

25. Huang, J., Roberts, A.J., Leschziner, A.E. & Reck-Peterson, S.L. Lis1 acts as a “clutch” between the ATPase and microtubule-binding domains of the dynein motor. Cell 150, 975–986 (2012).

26. Lammers, L.G. & Markus, S.M. The dynein cortical anchor Num1 activates dynein motility by relieving Pac1/LIS1-mediated inhibition. J Cell Biol 211, 309–322 (2015).

27. Torisawa, T. et al. Functional dissection of LIS1 and NDEL1 towards understanding the molecular mechanisms of cytoplasmic dynein regulation. J Biol Chem 286, 1959–1965 (2011).

28. Kusakci, E. et al. Lis1 slows force-induced detachment of cytoplasmic dynein from microtubules. Nat Chem Biol 20, 521–529 (2024).

29. Yamada, M. et al. LIS1 and NDEL1 coordinate the plus-end-directed transport of cytoplasmic dynein. EMBO J 27, 2471–2483 (2008).

30. Lenz, J.H., Schuchardt, I., Straube, A. & Steinberg, G. A dynein loading zone for retrograde endosome motility at microtubule plus-ends. EMBO J 25, 2275–2286 (2006).

31. Elshenawy, M.M. et al. Lis1 activates dynein motility by modulating its pairing with dynactin. Nat Cell Biol 22, 570–578 (2020).

32. Htet, Z.M. et al. LIS1 promotes the formation of activated cytoplasmic dynein-1 complexes. Nat Cell Biol 22, 518–525 (2020).

33. Marzo, M.G., Griswold, J.M. & Markus, S.M. Pac1/LIS1 stabilizes an uninhibited conformation of dynein to coordinate its localization and activity. Nat Cell Biol 22, 559–569 (2020).

34. Qiu, R., Zhang, J. & Xiang, X. LIS1 regulates cargo-adapter-mediated activation of dynein by overcoming its autoinhibition in vivo. J Cell Biol 218, 3630–3646 (2019).

35. Qiu, R., Zhang, J., Rotty, J.D. & Xiang, X. Dynein activation in vivo is regulated by the nucleotide states of its AAA3 domain. Curr Biol 31, 4486–4498 e4486 (2021).

36. Karasmanis, E.P. et al. Lis1 relieves cytoplasmic dynein-1 autoinhibition by acting as a molecular wedge. Nat Struct Mol Biol 30, 1357–1364 (2023).

37. Zhao, Y., Oten, S. & Yildiz, A. Nde1 promotes Lis1-mediated activation of dynein. Nat Commun 14, >7221 (2023).

38. Gillies, J.P. et al. Structural basis for cytoplasmic dynein-1 regulation by Lis1. Elife 11 (2022).

39. Kendrick, A.A., et al. Cryo-EM visualizes multiple steps of dynein’s activation pathway. bioRxiv (2024).

40. Baumbach, J. et al. Lissencephaly-1 is a context-dependent regulator of the human dynein complex. Elife 6 (2017).

41. DeSantis, M.E. et al. Lis1 Has Two Opposing Modes of Regulating Cytoplasmic Dynein. Cell 170, 1197–1208 e1112 (2017).

42. Wei, N. & Deng, X.W. The COP9 signalosome. Annu Rev Cell Dev Biol 19, 261–286 (2003).

43. Wolf, D.A., Zhou, C. & Wee, S. The COP9 signalosome: an assembly and maintenance platform for cullin ubiquitin ligases? Nat Cell Biol 5, 1029–1033 (2003).

44. Cope, G.A. & Deshaies, R.J. COP9 signalosome: a multifunctional regulator of SCF and other cullin-based ubiquitin ligases. Cell 114, 663–671 (2003).

45. Schulze-Niemand, E. & Naumann, M. The COP9 signalosome: A versatile regulatory hub of Cullin-RING ligases. Trends Biochem Sci 48, 82–95 (2023).

46. Hu, J., McCall, C.M., Ohta, T. & Xiong, Y. Targeted ubiquitination of CDT1 by the DDB1-CUL4A-ROC1 ligase in response to DNA damage. Nat Cell Biol 6, 1003–1009 (2004).

47. Nishitani, H. et al. Two E3 ubiquitin ligases, SCF-Skp2 and DDB1-Cul4, target human Cdt1 for proteolysis. EMBO J 25, 1126–1136 (2006).

48. Higa, L.A., Mihaylov, I.S., Banks, D.P., Zheng, J. & Zhang, H. Radiation-mediated proteolysis of CDT1 by CUL4-ROC1 and CSN complexes constitutes a new checkpoint. Nat Cell Biol 5, 1008–1015 (2003).

49. Lam, C., Vergnolle, M.A., Thorpe, L., Woodman, P.G. & Allan, V.J. Functional interplay between LIS1, NDE1 and NDEL1 in dynein-dependent organelle positioning. J Cell Sci 123, 202–212 (2010).

50. Pu, J., Guardia, C.M., Keren-Kaplan, T. & Bonifacino, J.S. Mechanisms and functions of lysosome positioning. J Cell Sci 129, 4329–4339 (2016).

51. Kapitein, L.C. et al. Probing intracellular motor protein activity using an inducible cargo trafficking assay. Biophys J 99, 2143–2152 (2010).

52. Shailesh, H., Zakaria, Z.Z., Baiocchi, R. & Sif, S. Protein arginine methyltransferase 5 (PRMT5) dysregulation in cancer. Oncotarget 9, 36705–36718 (2018).

53. Vinet, M. et al. Protein arginine methyltransferase 5: A novel therapeutic target for triple-negative breast cancers. Cancer Med 8, 2414–2428 (2019).

54. Mulvaney, K.M. et al. Molecular basis for substrate recruitment to the PRMT5 methylosome. Mol Cell 81, 3481–3495 e3487 (2021).

55. Owens, J.L. et al. PRMT5 Cooperates with pICln to Function as a Master Epigenetic Activator of DNA Double-Strand Break Repair Genes. iScience 23, 100750 (2020).

56. Guderian, G. et al. RioK1, a new interactor of protein arginine methyltransferase 5 (PRMT5), competes with pICln for binding and modulates PRMT5 complex composition and substrate specificity. J Biol Chem 286, 1976–1986 (2011).

57. Lacroix, M. et al. The histone-binding protein COPR5 is required for nuclear functions of the protein arginine methyltransferase PRMT5. EMBO Rep 9, 452–458 (2008).

58. Moon, H.M. & Wynshaw-Boris, A. Cytoskeleton in action: lissencephaly, a neuronal migration disorder. Wiley Interdiscip Rev Dev Biol 2, 229–245 (2013).

59. Cardoso, C. et al. Clinical and molecular basis of classical lissencephaly: Mutations in the LIS1 gene (PAFAH1B1). Hum Mutat 19, 4–15 (2002).

60. Stenson, P.D. et al. The Human Gene Mutation Database (HGMD((R))): optimizing its use in a clinical diagnostic or research setting. Hum Genet 139, 1197–1207 (2020).

61. Caspi, M. et al. LIS1 missense mutations: variable phenotypes result from unpredictable alterations in biochemical and cellular properties. J Biol Chem 278, 38740–38748 (2003).

62. Niekamp, S., Stuurman, N., Zhang, N. & Vale, R.D. Three-color single-molecule imaging reveals conformational dynamics of dynein undergoing motility. Proc Natl Acad Sci U S A 118 (2021).

63. Carter, A.P. Crystal clear insights into how the dynein motor moves. J Cell Sci 126, 705–713 (2013).

64. Gao, F.J. et al. GSK-3beta Phosphorylation of Cytoplasmic Dynein Reduces Ndel1 Binding to Intermediate Chains and Alters Dynein Motility. Traffic 16, 941–961 (2015).

65. Vaughan, P.S., Leszyk, J.D. & Vaughan, K.T. Cytoplasmic dynein intermediate chain phosphorylation regulates binding to dynactin. J Biol Chem 276, 26171–26179 (2001).

66. Chanduri, M. et al. Inositol hexakisphosphate kinase 1 (IP6K1) activity is required for cytoplasmic dynein-driven transport. Biochem J 473, 3031–3047 (2016).

67. Sapir, T., Cahana, A., Seger, R., Nekhai, S. & Reiner, O. LIS1 is a microtubule-associated phosphoprotein. Eur J Biochem 265, 181–188 (1999).

68. McKenney, R.J., Vershinin, M., Kunwar, A., Vallee, R.B. & Gross, S.P. LIS1 and NudE induce a persistent dynein force-producing state. Cell 141, 304–314 (2010).

69. Brown, J.S. et al. Neddylation promotes ubiquitylation and release of Ku from DNA-damage sites. Cell Rep 11, 704–714 (2015).

70. Rabut, G. & Peter, M. Function and regulation of protein neddylation. ‘Protein modifications: beyond the usual suspects’ review series. EMBO Rep 9, 969–976 (2008).

71. Kumar, P. et al. A Human Tyrosine Phosphatase Interactome Mapped by Proteomic Profiling. J Proteome Res 16, 2789–2801 (2017).

72. Perez-Riverol, Y. et al. The PRIDE database resources in 2022: a hub for mass spectrometry-based proteomics evidences. Nucleic Acids Res 50, D543–D552 (2022).

73. Coleman, K.E. et al. SENP8 limits aberrant neddylation of NEDD8 pathway components to promote cullin-RING ubiquitin ligase function. Elife 6 (2017).

74. Sato, Y., Murakami, Y. & Takahashi, M. Semi-retentive cytoskeletal fractionation (SERCYF): A novel method for the biochemical analysis of the organization of microtubule and actin cytoskeleton networks. Biochem Biophys Res Commun 488, 614–620 (2017).

75. Kumar, G. et al. RUFY3 links Arl8b and JIP4-Dynein complex to regulate lysosome size and positioning. Nat Commun 13, 1540 (2022).

