## Supplementary data for "COP9 signalosome and PRMT5 methylosome complexes are essential regulators of Lis1-dynein based transport"

**This file contains Supplementary figures S1 – S13, and supplementary videos 1 – 8, and Supplementary table 1.**

#### Supplementary Figures

**Figure S1: (A)** HEK 293T cells were co-transfected with SFB-CSN1, SFB-CSN2, SFB-CSN3, SFB-CSN4, SFB-CSN5, SFB-CSN6, SFB-CSN7A, SFB-CSN8, and SFB-CSNAP in combination with Myc-Lis1 plasmids. Cells were then lysed, and incubated with S-protein beads. Interaction was detected by immunoblotting with anti-Myc antibody. **(B)** Bacterially purified GST-CSN1, GST-CSN2, GST-CSN3, GST-CSN4, GST-CSN5, GST-CSN6, GST-CSN7A, and GST-CSN8 were incubated with MBP-Lis1 purified protein. Interaction was detected by immunoblotting with anti-MBP antibody. A pictorial representation of the arrangement of CSN subunits in the complex is shown. CBB: Coomassie Brilliant Blue. **(C)** Pictorial representation of the domain architecture of Lis1 protein is shown. **(D)** HEK 293T cells were co-transfected with SFB-Lis1 WT, SFB-Lis1 ΔLisH, and SFB-Lis1 ΔWD in combination with Myc-CSN5 plasmids. Cells were then lysed, and incubated with S-protein beads. Interaction was detected by immunoblotting with anti-Myc antibody. **(E)** Bacterially purified GST-CSN1, GST-CSN2, GST-CSN3 and GST-CSN4 were incubated with MBP-Lis1 ΔLisH and MBP-Lis1 ΔWD purified proteins. Interaction was detected by immunoblotting with anti-MBP antibody. CBB: Coomassie Brilliant Blue. **(F)** HEK 293T cells were co-transfected with SFB-CSN1, SFB-CSN2, SFB-CSN3, SFB-CSN4, SFB-CSN5, SFB-CSN6, SFB-CSN7A, and SFB-CSN8 in combination with Myc-DIC1 plasmids. Cells were then lysed, and incubated with S-protein beads. Interaction was detected by immunoblotting with anti-Myc antibody. **(G)** Control and CSN5 depleted HEK 293T cells were immunoprecipiated with DIC antibody. Interaction was tested by immunoblotting with Lis1 antibody. **(H)** HEK 293T cells were co-transfected with SFB-DIC1 and Myc-Lis1 plasmids. Cells were treated with 1 uM

CSN5i3 for 12 hours. 24 hours after transfection, cells were lysed and incubated with S-protein beads. Interaction was tested by immunoblotting with anti-Myc antibody.

**Figure S2: (A)** U2OS cells were transfected with SFB-DIC1 and eGFP-CSN7A plasmids. 4 hours before fixation, cells were treated with 1 uM Nocodazole and 2 uM Paclitaxel. Cells were labelled with α-tubulin, and anti-Flag antibodies, and imaged using a super-resolution microscope with SIM module to determine their colocalization. Scale bars: 10 μm. **(B)** Quantification of the Pearson correlation coefficient (n = 27 cells) across three biological replicates for the experiment shown in (A). Data are median ± interquartile range. (***p < 0.0001; One way ANOVA followed by Dunnett's multiple comparisons test). **(C)** U2OS cells were transfected with SFB-DIC1 and eGFP-CSN7A plasmids. Cells were given a cold shock with cold media kept at 4°C and keeping the dish on ice for 5 minutes with immediate fixation. For recovery, warm media kept at 37°C was added, and dish was kept in 37°C incubator for 5 minutes with immediate fixation. Cells were labelled with α-tubulin, and anti-Flag antibodies, and imaged using a super-resolution microscope with SIM module to determine their colocalization. Scale bars: 10 μm. **(D)** Quantification of the Pearson correlation coefficient (n = 22 cells) across three biological replicates for the experiment shown in (C). Data are median ± interquartile range. (***p < 0.0001; One way ANOVA followed by Dunnett's multiple comparisons test). **(E)** U2OS cells transfected with SFB-CSN1 plasmid was fixed and labelled with anti-DHC, anti- α-tubulin, anti-p150, anti-BicD2 along with anti-flag antibodies and imaged using a super-resolution microscope with SIM module to determine their colocalization. Scale bars: 10 μm. **(F)** Quantification of the Pearson correlation coefficient (n = 60 cells) for the experiment shown in (E). **(G)** HEK 293T cells were transfected with SFB-vector, SFB-CSN1, and SFB-DIC1 plasmids. Cells were then lysed, and incubated with S-protein beads. Interaction was detected by immunoblotting with indicated antibodies. The densitometric quantifications from three biological replicates are shown under each blot. **(H)**

Quantification of the speed of the moving dynein particles in mCherry-DIC1 alone and in combination with eGFP-CSN1 samples (110 dynein particles from n = 8 cells) across two biological replicates. (***$p < 0.0001$; Two tailed t test).

**Figure S3: (A)** HEK 293T cells were transfected with SFB-Cul1, SFB-Cul4A, SFB-Cul7, and SFB-Lis1, in combination with Myc-DIC1. Cells were lysed and incubated with S-protein beads. Interaction was detected by probing with anti-Myc antibody. **(B)** HEK 293T cells were transfected with SFB-vector, SFB-CSN1, SFB-Cul1, SFB-Cul4A, and SFB-Cul7 plasmids. The lysates were probed with indicated antibodies. **(C)** HEK 293T cells were transfected with dynein subunits SFB-DIC1, SFB-DIC2, SFB-LIC1, SFB-LIC2, SFB-LC8, SFB-Tctex1, SFB-LC7 with or without Myc-CSN1 plasmids. The lysates were probed with indicated antibodies. **(D)** HEK 293T cells were transfected with increasing concentrations (0, 0.5, 1, 2, 4 µg) of SFB-Cul1, SFB-Cul4A, and SFB-Cul7 plasmids. The samples were immunoblotted with indicated antibodies. **(E)** Bacterially purified GST-DIC1 was incubated with NAE1, UBC12 and Nedd8 for in vitro neddylation for 60 mins, and with purified CSN for in vitro deneddylation for 30 mins. His-Lis1 (2 ug) or an unrelated protein His-CDC25A as a negative control (2 ug) were added during deneddylation to see the kinetics of the reaction. Neddylated protein was visualized by immunoblotting with anti-Nedd8 antibody. CBB: Coomassie Brilliant Blue.

**Figure S4: (A)** A schematic of lysosome and endosome distribution pattern in cells is shown. **(B)** A schematic of golgi distribution pattern in cells is shown. **(C)** Control and Lis1 depleted stable U2OS cells were fixed and labelled with LAMP2 (lysosomes), EEA1 (early endosome), GM130 (cis-Golgi) and p230 (trans-Golgi) antibodies to visualise their distribution under confocal microscope. Scale bars: 10 µm. **(D)** Quantification of the distribution of LAMP2-positive compartments (n=50 cells), EEA1-positive compartments (n=15 cells) across three biological replicates for the experiments shown in (C) (***$p < 0.0001$; ns not significant; two-tailed Student's *t*-test). The golgi distribution has been categorized into 3 types: ribbon, broken ribbon and scatter. The values plotted are the mean ± SE from three independent experiments. (***p < 0.0004; **p < 0.0063; *p < 0.0141; Two-way ANOVA followed by Dunnett's multiple comparisons test). **(E)** Control and Lis1 depleted stable IMR32 cells were fixed and labelled with LAMP2 (lysosomes), EEA1 (early endosome), GM130 (cis-Golgi) and p230 (trans-Golgi) antibodies to visualise their distribution under confocal microscope. Scale bars: 10 μm. **(F)** Quantification of the distribution of LAMP2-positive compartments (n=20 cells), EEA1-positive compartments (n=26 cells) across three biological replicates for the experiments shown in (E) (***p < 0.0001; two-tailed Student's *t*-test). The golgi distribution has been categorized into 3 types: ribbon, broken ribbon and scatter. The values plotted are the mean ± SE from three independent experiments. ((***p < 0.0001, **p 0.0010, *p < 0.0107; ns not significant; Two-way ANOVA followed by Dunnett's multiple comparisons test).

**Figure S5: (A)** IMR32 cells were treated with DMSO, 3 uM MLN4924, and 3 uM CSN5i3 for 4 hours before fixation, and labelled with LAMP2 (lysosomes), EEA1 (early endosome), GM130 (cis-Golgi) and p230 (trans-Golgi) antibodies to visualise their distribution under confocal microscope. Scale bars: 10 μm. **(B)** Quantification of the distribution of LAMP2-positive compartments (n=31 cells), EEA1-positive compartments (n=26 cells) across three biological replicates for the experiments shown in (A) (***p < 0.0001; ns not significant; One way ANOVA followed by Dunnett's multiple comparisons test). The golgi distribution has been categorized into 3 types: ribbon, broken ribbon and scatter. The values plotted are the mean ± SE from three independent experiments. (**p 0.0094, *p 0.0364, ns not significant; Two-way ANOVA followed by Dunnett's multiple comparisons test). **(C)** Video stills of U2OS cells transfected with Pex3-mEmerald-FKBP and FRB-BicD2-S, treated with DMSO, MLN4924, and CSN5i3. Individual moving peroxisomes are highlighted with arrows with their corresponding kymographs shown in **(D)**. The scale bar of video stills and kymographs is 1 μm. **(E)** U2OS cells were transfected with Pex3-mEmerald-FKBP and FRB-BicD2-S along with mCherry-CSN1. Cells were treated with 1 uM Rapamycin just before live-cell imaging. Video stills and the **(F)** kymographs of the arrow-marked individual moving peroxisomes are shown. Scale: 1 µm. **(G)** Quantification of the speed of peroxisome in U2OS cells for the experiment shown in (E). Each color represents a replicate. Bigger circle is the mean of the data points represented as smaller circles from each biological replicate (n = 25, 40, 40 cells). Data are mean ± SE range. (***p < 0.0001; One way ANOVA). **(H)** Quantification of the net displacement of peroxisome in U2OS cells for the experiment shown in (E). Each color represents a replicate. Bigger circle is the mean of the data points represented as smaller circles from each biological replicate (n = 25, 40, 40 cells). Data are mean ± SE (***p < 0.0001; One way ANOVA).

**Figure S6: (A)** U2OS cells were transiently treated with DIC shRNA and the knockdown was tested by probing with respective antibodies. **(B)** Video stills of DIC depleted U2OS cells transfected with Pex3-mEmerald-FKBP and FRB-BicD2-S, in addition to DIC1 WT and DIC1 K42R. The individual moving peroxisomes are highlighted with arrows with their corresponding kymographs are shown in **(C)**. The scale bar of video stills and kymographs is 1 µm. **(D)** Control and DIC depleted U2OS cells were fixed and labelled with LAMP2 (lysosomes), EEA1 (early endosome), GM130 (cis-Golgi) and p230 (trans-Golgi) antibodies to visualise their distribution under confocal microscope. Scale bars: 10 µm. **(E)** Quantification of the distribution of LAMP2-positive compartments (n=17 cells), EEA1-positive compartments (n=25 cells) across three biological replicates for the experiments shown in (D) (***p < 0.0001; ** p 0.0020; One way ANOVA followed by Dunnett's multiple comparisons test). The golgi distribution has been categorized into 3 types: ribbon, broken ribbon and scatter. The values plotted are the mean ± SE from three independent experiments. (***p 0.0004, **p 0.0014, *p 0.0142, ns not significant; Two-way ANOVA followed by Dunnett's multiple comparisons test).

**Figure S7: (A)** Control and DIC depleted U2OS cells were transfected with SFB-DIC1 WT and SFB-DIC1 K42R plasmids. Cells were fixed and labelled with EEA1 (early endosomes), GM130 (cis-Golgi), p230 (trans-Golgi) antibodies to visualise their distribution under confocal microscope. Scale bars: 10 μm. **(B)** Quantification of the distribution of EEA1-positive compartments (n = 26 cells) across three biological replicates for the experiments shown in (A). Data are median ± interquartile range. (***p < 0.0001; **p 0.0056; two-tailed Student's t-test). The values plotted for golgi distribution are mean ± SE from three independent experiments shown in (A). (***p < 0.0003, **p 0.0091, *p 0.0251, ns not significant; Two-way ANOVA followed by Dunnett's multiple comparisons test).

**Figure S8: (A)** HEK 293T cells were transfected with SFB-DIC1 and SFB-DIC1 K42R plasmids. Cells were then lysed, and incubated with Dynein Heavy Chain (DHC)-bound Protein G beads. Interaction was detected by immunoblotting with anti-Flag antibody. CBB: Coomassie Brilliant Blue. **(B)** HEK 293T cells were transfected with SFB-DIC1 WT and SFB-DIC1 K42R and Myc-LIC1 plasmids. Cells were then lysed, and incubated with S-protein beads. Interaction was detected by immunoblotting with anti-Myc antibody. **(C)** HEK 293T cells were transfected with SFB-LC8, Tctex1, and LC7 with Myc-DIC1 plasmids. Cells were treated with 1 uM MLN4924 (NAE1 inhibitor) or DMSO for 12 hours. Cells were lysed and further incubated with S-protein beads. Interaction was detected by immunoblotting with anti-Myc antibody. **(D)** Bacterially purified recombinant GST-DIC1 WT and GST-DIC1 K42R were incubated with purified MBP-Nde1. The eluates were immunoblotted with anti-MBP antibody. CBB: Coomassie Brilliant Blue. **(E)** HEK 293T cells were transfected with SFB DIC1 WT and K42R plasmids. 24 hours after transfection, cycloheximide (50 μg/ml) was added, and cells were collected at the indicated time points. The protein levels were detected by immunoblotting with respective antibodies. **(F)** HEK 293T cells were transfected with SFB DIC1 and K42R in combination with Myc DIC1 WT and K42R. Cells were lysed, and incubated with S-protein beads. Interaction was detected by immunoblotting with anti-Myc antibody.

**Figure S9: (A)** HEK 293T cells were co-transfected with SFB-PRMT5, SFB-WDR77, and SFB-pICln in combination with Myc-Lis1 plasmids. Cells were then lysed, and incubated with streptavidin-sepharose beads. Interaction was detected by immunoblotting with anti-Myc antibody. **(B)** IMR32 cell lysates were subjected to immunoprecipitation with either IgG or PRMT5 antibody. Presence of Lis1 immunoprecipitates was detected by western blotting with Lis1 antibody. **(C)** HEK 293T cells were co-transfected with SFB-PRMT5, SFB-WDR77, and SFB-pICln in combination with DIC1-HA, and DIC2-HA plasmids. Cells were then lysed, and incubated with streptavidin-sepharose beads. Interaction was detected by immunoblotting with anti-HA antibody. **(D)** Control and PRMT5 depleted stable IMR32 cells were fixed and labelled with LAMP2 (lysosomes), EEA1 (early endosome), GM130 (cis-Golgi) and p230 (trans-Golgi) antibodies to visualise their distribution under confocal microscope. Scale bars: 10 μm. **(E)** Quantification of the distribution of LAMP2-positive compartments (n=31 cells), EEA1-positive compartments (n=25 cells) across three biological replicates for the experiments shown in (D) (***$p < 0.0001$; One way ANOVA followed by Dunnett's multiple comparisons test). The golgi distribution has been categorized into 3 types: ribbon, broken ribbon and scatter. The values plotted are the mean ± SE from three independent experiments. (***$p < 0.0002$, **p 0.0014, *p 0.0140, ns not significant; Two-way ANOVA followed by Dunnett's multiple comparisons test). **(F)** Video stills of Control and PRMT5 depleted U2OS cells transfected with Pex3-mEmerald-FKBP and FRB-BicD2S plasmids are shown. The individual moving peroxisomes are highlighted with arrows with their corresponding kymographs are shown in **(G)**. The scale bar of video stills and kymographs is 1 μm.

**Figure S10: (A)** U2OS cells were depleted of WDR77 using two WDR77 shRNA targeting at different sites and the knockdown was tested by probing with respective antibodies. **(B)** Control and WDR77 depleted stable U2OS cells were fixed and labelled with LAMP2 (lysosomes), EEA1 (early endosome), GM130 (cis-Golgi) and p230 (trans-Golgi) antibodies to visualise their distribution under confocal microscope. Scale bars: 10 µm. **(C)** Quantification of the distribution of LAMP2-positive compartments (n=40 cells), EEA1-positive compartments (n=32 cells) across three biological replicates for the experiments shown in (B) (***p < 0.0001; two-tailed Student's *t*-test). The golgi distribution has been categorized into 3 types: ribbon, broken ribbon and scatter. The values plotted are the mean ± SE from three independent experiments. (***p < 0.0006; **p < 0.0050; *p < 0.0157; Two-way ANOVA followed by Dunnett's multiple comparisons test). **(D)** U2OS cells were treated with EPZ015666 at indicated concentrations for 4 hours before fixation. Cells were labelled with LAMP2 (lysosomes), EEA1 (early endosome), GM130 (cis-Golgi) and p230 (trans-Golgi) antibodies to visualise their distribution under confocal microscope. Scale bars: 10 µm. **(E)** Quantification of the distribution of LAMP2-positive compartments (n=80,78,56 cells for each sample), EEA1-positive compartments (n=24 cells) across three replicates for the experiments shown in (D) (***p < 0.0001; One way ANOVA followed by Dunnett's multiple comparisons test). The golgi distribution has been categorized into 3 types: ribbon, broken ribbon and scatter. The values plotted are the mean ± SE from three independent experiments. (**p < 0.0028, *p 0.0281, ns not significant; Two-way ANOVA followed by Dunnett's multiple comparisons test).

**Figure S11: (A)** Control and PRMT5 depleted stable U2OS cells were transfected with shRNA resistant PRMT5 WT and its catalytic dead G367A:R368A plasmids. Cells were fixed and labelled with EEA1 (early endosomes), GM130 (cis-Golgi), and p230 (trans-Golgi) antibody to visualize their distribution under confocal microscope. Scale bars: 10 µm. **(B)** Quantification of the distribution of EEA1-positive compartments (n = 30 cells) across three biological replicates for the experiments shown in (A). Data are median ± interquartile range.

(***p < 0.0001; ns not significant; two-tailed Student's t-test). The values plotted for GM130 and p230 distribution are the mean ± SE from three independent experiments shown in (C). (**p 0.0077, ns not significant; Two-way ANOVA followed by Dunnett's multiple comparisons test), and for p230 (***p < 0.0006, **p 0.0041, *p 0.0111; ns not significant; Two-way ANOVA followed by Dunnett's multiple comparisons test).

**Figure S12: (A)** Control and WDR77 depleted stable HEK293T cells were transfected with SFB-Lis1 plasmid. 24 hours after transfection, cells were lysed, denatured and incubated with streptavidin-sepharose beads. Methylated proteins were detected with immunoblotting using anti-SDMA antibody. **(B)** Control and Lis1 depleted U2OS cells were transfected with shRNA resistant Lis1 WT, Lis1 R238A, and R342A plasmids. Cells were fixed and labelled with LAMP2 (lysosomes) antibody to visualise their distribution under confocal microscope. Scale bars: 10 μm. **(C)** HEK 293T cells were transfected with SFB-Lis1 WT or SFB-Lis1 R238A plasmids. 24 hours after transfection, cells were lysed and immunoprecipitated with DHC antibody. The interaction was detected by immunoblotting with anti-flag antibody. CBB: Coomassie Brilliant Blue. The densitometric quantifications from three biological replicates are shown under each blot. **(D)** HEK 293T cells were transfected with SFB Lis1 WT and R238A plasmids. 24 hours after transfection, cycloheximide (50 μg/ml) was added, and cells were collected at the indicated time points. The protein levels were detected by immunoblotting with respective antibodies. **(E)** HEK 293T cells were transfected with SFB Lis1 WT and R238A in combination with Myc Lis1 WT and R238A plasmids. Cells were lysed, and incubated with S-protein beads. Interaction was detected by immunoblotting with anti-Myc antibody.

**Figure S13: (A)** Control and Lis1 depleted stable U2OS cells were transfected with shRNA resistant Lis1 WT, Lis1 R238K, and R238F plasmids. Cells were fixed and labelled with EEA1 (early endosomes), GM130 (cis-Golgi), and p230 (trans-Golgi) antibody to visualize their distribution under confocal microscope. Scale bars: 10 μm. **(B)** Quantification of the distribution of EEA1-positive compartments (n = 40 cells) across three biological replicates for the experiments shown in (A). Data are median ± interquartile range. (***$p < 0.0001$; ns not significant; two-tailed Student's t-test). The values plotted for the golgi distribution are the mean ± SE from three independent experiments shown in (***$p < 0.0007$, *$p < 0.0432$, ns not significant; Two-way ANOVA followed by Dunnett's multiple comparisons test). **(C)** Video stills of Lis1 depleted U2OS cells transfected with Pex3-mEmerald-FKBP and FRB-BicD2-S, in addition to shRNA resistant Lis1 WT, Lis1 RK, and Lis1 RF plasmids. The individual moving peroxisomes are highlighted with arrows with their corresponding kymographs are shown in **(D)**. The scale bar of video stills and kymographs is 1 µm. **(E)** HEK 293T cells were transfected with SFB-Lis1 plasmid. Cells were treated with 1 uM MLN4924 (NAE1 inhibitor) for 12 hours. Cells were lysed under denaturing conditions, and further incubated with streptavidin-sepharose beads. Methylated proteins were detected by immunoblotting with anti-symmetric dimethyl arginine antibody. **(F)** HEK 293T cells were co-transfected with SFB-LIC1 and Myc-DIC1 plasmids. Cells were treated with EPZ015666 (PRMT5 inhibitor) for 12 hours. Cells were lysed and further incubated with S-protein beads. Interaction was detected by immunoblotting with anti-Myc antibody.

#### Supplementary videos

**Video S1:** Tracking the colocalization of mCherry-DIC1 and eGFP-CSN1 in U2OS cells using live-cell super-resolution microscopy with SIM module. Scale bars: 5 µm.

**Video S2:** Tracking the movement of mCherry-DIC1 alone or in combination with eGFP-CSN1 in U2OS cells stained with Tubulin DeepRed using live-cell super-resolution microscopy with SIM module. Scale bars: 5 µm.

**Video S3:** Peroxisome motility assay in U2OS cells using Pex3-mEmerald-FKBP and FRB-BicD2S induced by 1 uM rapamycin in the presence of DMSO, MLN4924, and CSN5i3 using live-cell super-resolution microscopy with SIM module. Scale bars: 5 μm.

**Video S4:** Peroxisome motility assay in U2OS cells using Pex3-mEmerald-FKBP and FRB-BicD2S induced by 1 uM rapamycin in the presence of exogenously expressed mCherry-CSN1 using live-cell super-resolution microscopy with SIM module. Scale bars: 5 μm.

**Video S5:** Rescue of peroxisome motility assay in U2OS cells using Pex3-mEmerald-FKBP and FRB-BicD2S induced by 1 uM rapamycin in DIC knockdown U2OS cells with exogenously expressed mCherry-DIC1 WT and mCherry-DIC1 K42R using live-cell super-resolution microscopy with SIM module. Scale bars: 5 μm.

**Video S6:** Peroxisome motility assay in U2OS cells using Pex3-mEmerald-FKBP and FRB-BicD2S induced by 1 uM rapamycin in Control and PRMT5 knockdown stable U2OS cells using live-cell super-resolution microscopy with SIM module. Scale bars: 5 μm.

**Video S7:** Rescue of Peroxisome motility assay in U2OS cells using Pex3-mEmerald-FKBP and FRB-BicD2S induced by 1 uM rapamycin in Lis1 knockdown stable U2OS cells with exogenously expressed shRNA resistant mCherry-Lis1 WT, mCherry-Lis1 R238K and mCherry-Lis1 R238F using live-cell super-resolution microscopy with SIM module. Scale bars: 5 μm.

**Video S8:** Tracking the movement of mCherry-DIC1 and eGFP-Lis1 in U2OS cells using live-cell super-resolution microscopy with SIM module. Scale bars: 5 μm. The zoom clip of the moving dynein and Lis1 is shown on the right. Scale bars: 1 μm

### Figure S1

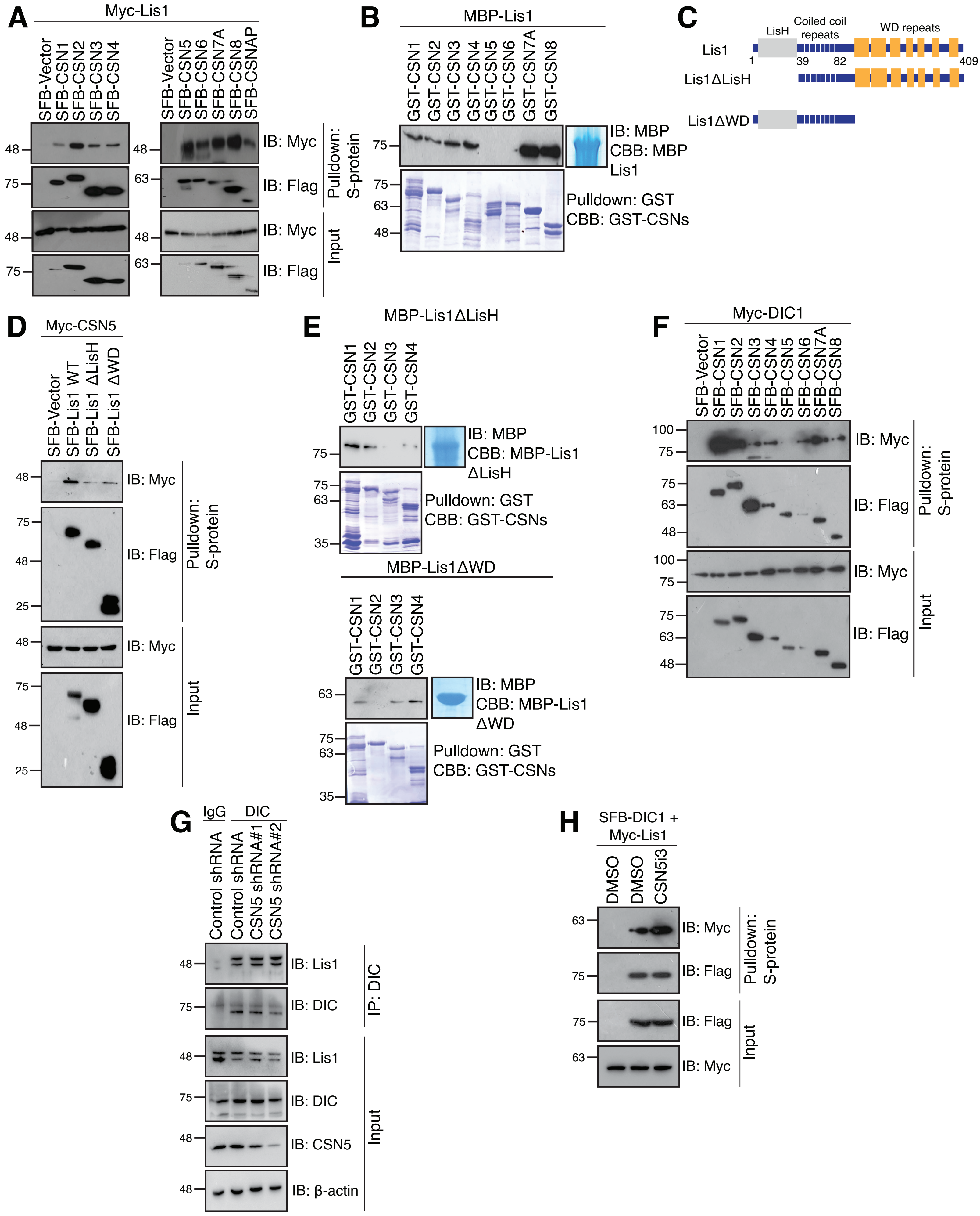

A
Myc-Lis1
SFB-Vector
SFB-CSN1
SFB-CSN2
SFB-CSN3
SFB-CSN4
SFB-CSN5
SFB-CSN6
SFB-CSN7A
SFB-CSN8
SFB-CSNAP
IB: Myc
IB: Flag
Pulldown: S-protein
Input
B
MBP-Lis1
GST-CSN1
GST-CSN2
GST-CSN3
GST-CSN4
GST-CSN5
GST-CSN6
GST-CSN7A
GST-CSN8
IB: MBP
CBB: MBP Lis1
Pulldown: GST
CBB: GST-CSNs
C
LisH
Coiled coil repeats
WD repeats
Lis1
Lis1ΔLisH
Lis1ΔWD
1
39
82
409
D
Myc-CSN5
SFB-Vector
SFB-Lis1 WT
SFB-Lis1 ΔLisH
SFB-Lis1 ΔWD
E
MBP-Lis1ΔLisH
CBB: MBP-Lis1 ΔLisH
MBP-Lis1ΔWD
CBB: MBP-Lis1 ΔWD
F
Myc-DIC1
G
IgG
DIC
Control shRNA
CSN5 shRNA#1
CSN5 shRNA#2
IB: Lis1
IB: DIC
IP: DIC
IB: CSN5
IB: β-actin
H
SFB-DIC1 + Myc-Lis1
DMSO
CSN5i3

### Figure S2

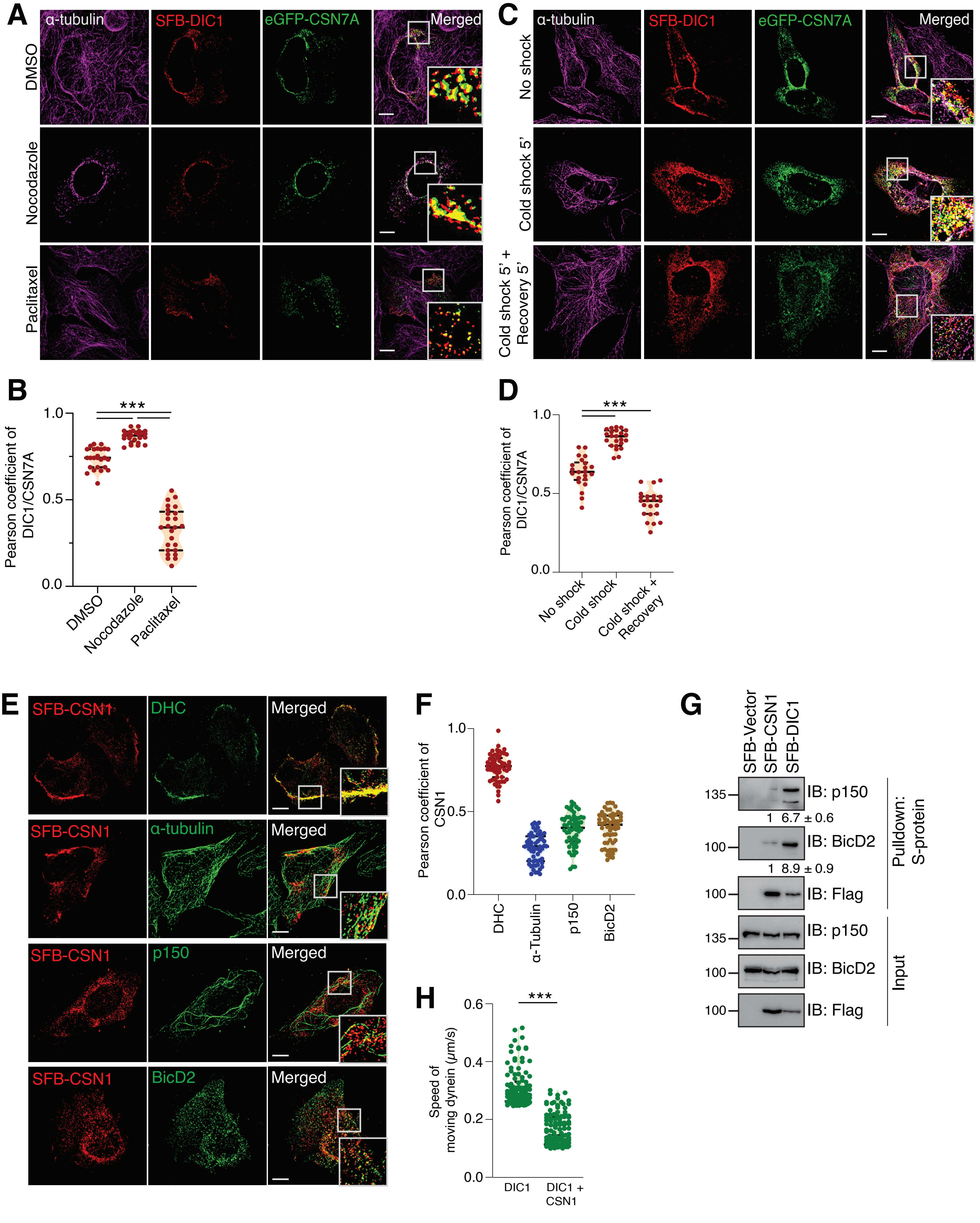

A
α-tubulin
SFB-DIC1
eGFP-CSN7A
Merged
DMSO
Nocodazole
Paclitaxel
C
α-tubulin
SFB-DIC1
eGFP-CSN7A
Merged
No shock
Cold shock 5'
Cold shock 5' + Recovery 5'
B
Pearson coefficient of DIC1/CSN7A
***
1.0
0.5
0.0
DMSO
Nocodazole
Paclitaxel
D
Pearson coefficient of DIC1/CSN7A
***
1.0
0.5
0.0
No shock
Cold shock
Cold shock + Recovery
E
SFB-CSN1
DHC
Merged
SFB-CSN1
α-tubulin
Merged
SFB-CSN1
p150
Merged
SFB-CSN1
BicD2
Merged
F
Pearson coefficient of CSN1
1.0
0.5
0.0
DHC
α-Tubulin
p150
BicD2
G
SFB-Vector
SFB-CSN1
SFB-DIC1
135
IB: p150
1
6.7 ± 0.6
100
IB: BicD2
1
8.9 ± 0.9
100
IB: Flag
Pulldown: S-protein
135
IB: p150
100
IB: BicD2
100
IB: Flag
Input
H
Speed of moving dynein (μm/s)
***
0.6
0.4
0.2
0.0
DIC1
DIC1 + CSN1

### Figure S3

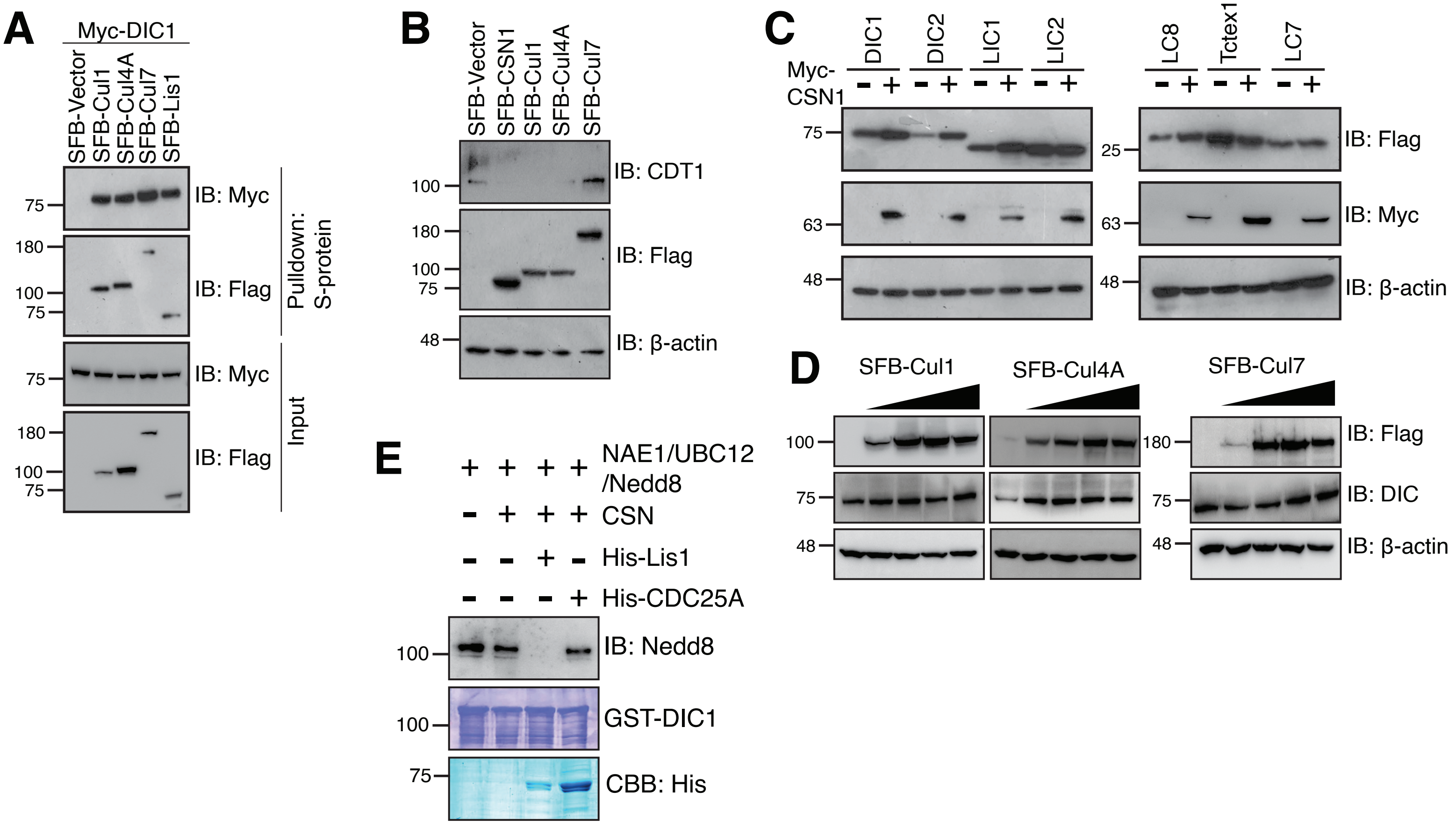
A
Myc-DIC1
SFB-Vector
SFB-Cul1
SFB-Cul4A
SFB-Cul7
SFB-Lis1
IB: Myc
IB: Flag
Pulldown: S-protein
Input
B
SFB-Vector
SFB-CSN1
SFB-Cul1
SFB-Cul4A
SFB-Cul7
IB: CDT1
IB: β-actin
C
Myc-CSN1
DIC1
DIC2
LIC1
LIC2
LC8
Tctex1
LC7
D
SFB-Cul1
SFB-Cul4A
SFB-Cul7
IB: DIC
E
NAE1/UBC12/Nedd8
CSN
His-Lis1
His-CDC25A
IB: Nedd8
GST-DIC1
CBB: His

### Figure S4

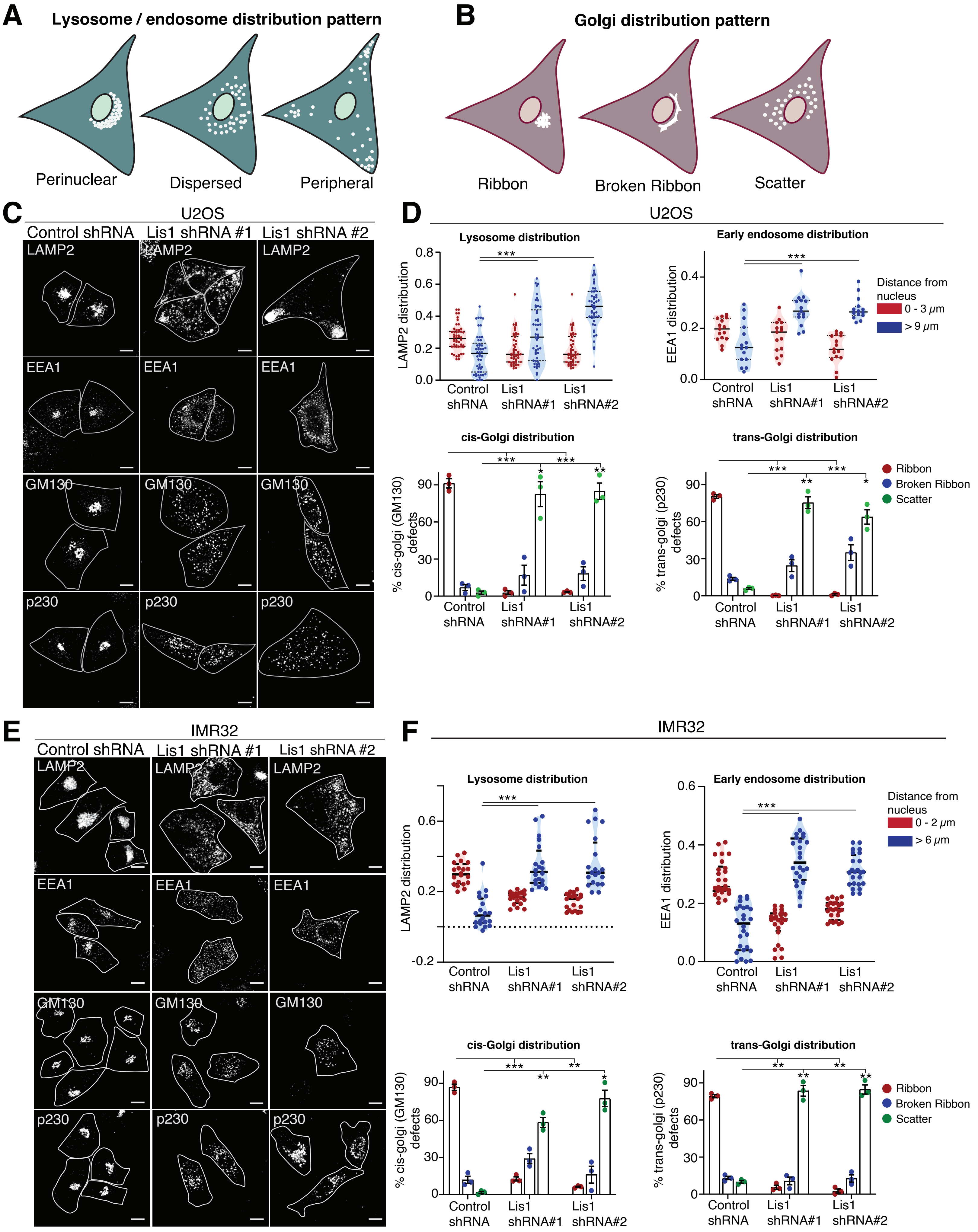
A
Lysosome / endosome distribution pattern
Perinuclear
Dispersed
Peripheral
B
Golgi distribution pattern
Ribbon
Broken Ribbon
Scatter
C
U2OS
Control shRNA
Lis1 shRNA #1
Lis1 shRNA #2
LAMP2
EEA1
GM130
p230
D
U2OS
Lysosome distribution
LAMP2 distribution
Early endosome distribution
EEA1 distribution
Distance from nucleus
0 - 3 μm
> 9 μm
Control shRNA
Lis1 shRNA#1
Lis1 shRNA#2
cis-Golgi distribution
% cis-golgi (GM130) defects
trans-Golgi distribution
% trans-golgi (p230) defects
Ribbon
Broken Ribbon
Scatter
E
IMR32
Control shRNA
Lis1 shRNA #1
Lis1 shRNA #2
LAMP2
EEA1
GM130
p230
F
IMR32
Lysosome distribution
LAMP2 distribution
Early endosome distribution
EEA1 distribution
Distance from nucleus
0 - 2 μm
> 6 μm
Control shRNA
Lis1 shRNA#1
Lis1 shRNA#2
cis-Golgi distribution
% cis-golgi (GM130) defects
trans-Golgi distribution
% trans-golgi (p230) defects
Ribbon
Broken Ribbon
Scatter

### Figure S5

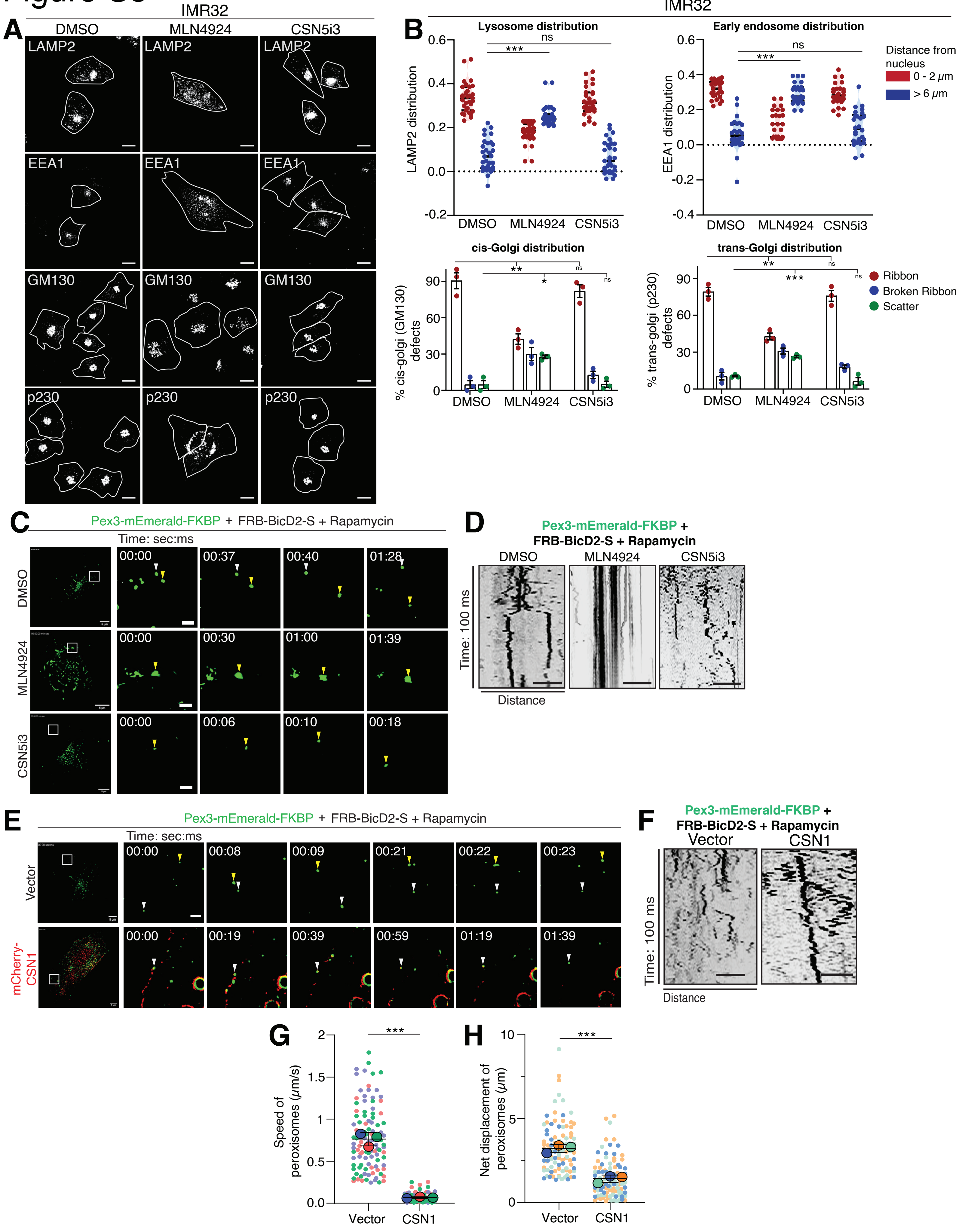
IMR32
A
DMSO
MLN4924
CSN5i3
LAMP2
EEA1
GM130
p230
B
Lysosome distribution
Early endosome distribution
LAMP2 distribution
EEA1 distribution
Distance from nucleus
0 - 2 μm
> 6 μm
cis-Golgi distribution
trans-Golgi distribution
% cis-golgi (GM130) defects
% trans-golgi (p230) defects
Ribbon
Broken Ribbon
Scatter
ns
C
Pex3-mEmerald-FKBP + FRB-BicD2-S + Rapamycin
Time: sec:ms
D
Time: 100 ms
Distance
E
Vector
mCherry-CSN1
F
CSN1
G
Speed of peroxisomes (μm/s)
H
Net displacement of peroxisomes (μm)

### Figure S6

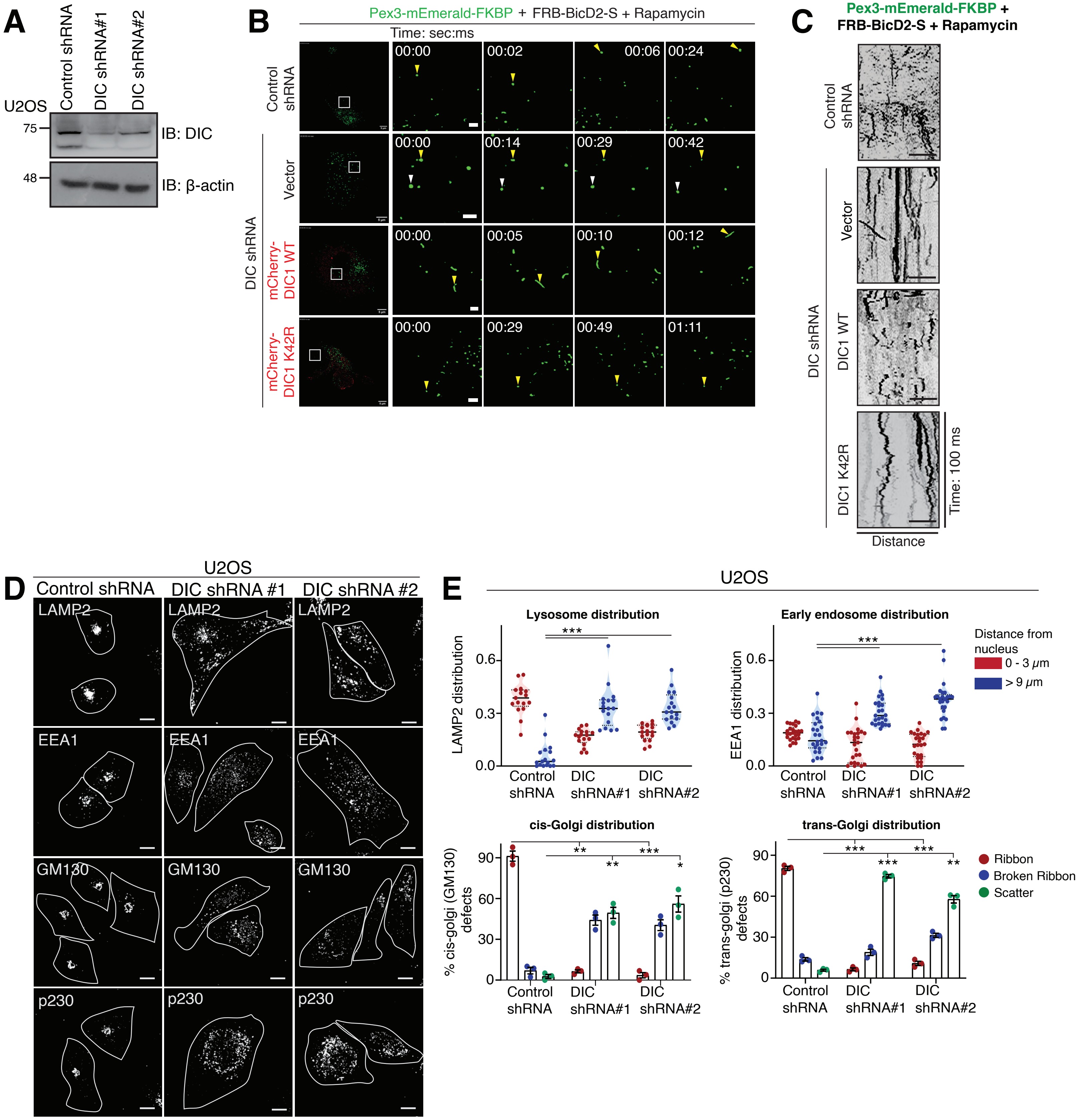

A
U2OS
Control shRNA
DIC shRNA#1
DIC shRNA#2
75
48
IB: DIC
IB: β-actin
B
Pex3-mEmerald-FKBP + FRB-BicD2-S + Rapamycin
Time: sec:ms
Control shRNA
DIC shRNA
Vector
mCherry-DIC1 WT
mCherry-DIC1 K42R
00:00
00:02
00:06
00:24
00:14
00:29
00:42
00:05
00:10
00:12
00:49
01:11
C
Pex3-mEmerald-FKBP + FRB-BicD2-S + Rapamycin
Control shRNA
DIC shRNA
Vector
DIC1 WT
DIC1 K42R
Time: 100 ms
Distance
D
U2OS
Control shRNA
DIC shRNA #1
DIC shRNA #2
LAMP2
EEA1
GM130
p230
E
U2OS
Lysosome distribution
LAMP2 distribution
Early endosome distribution
EEA1 distribution
Distance from nucleus
0 - 3 μm
> 9 μm
cis-Golgi distribution
% cis-golgi (GM130) defects
trans-Golgi distribution
% trans-golgi (p230) defects
Ribbon
Broken Ribbon
Scatter
Control shRNA
DIC shRNA#1
DIC shRNA#2

### Figure S7

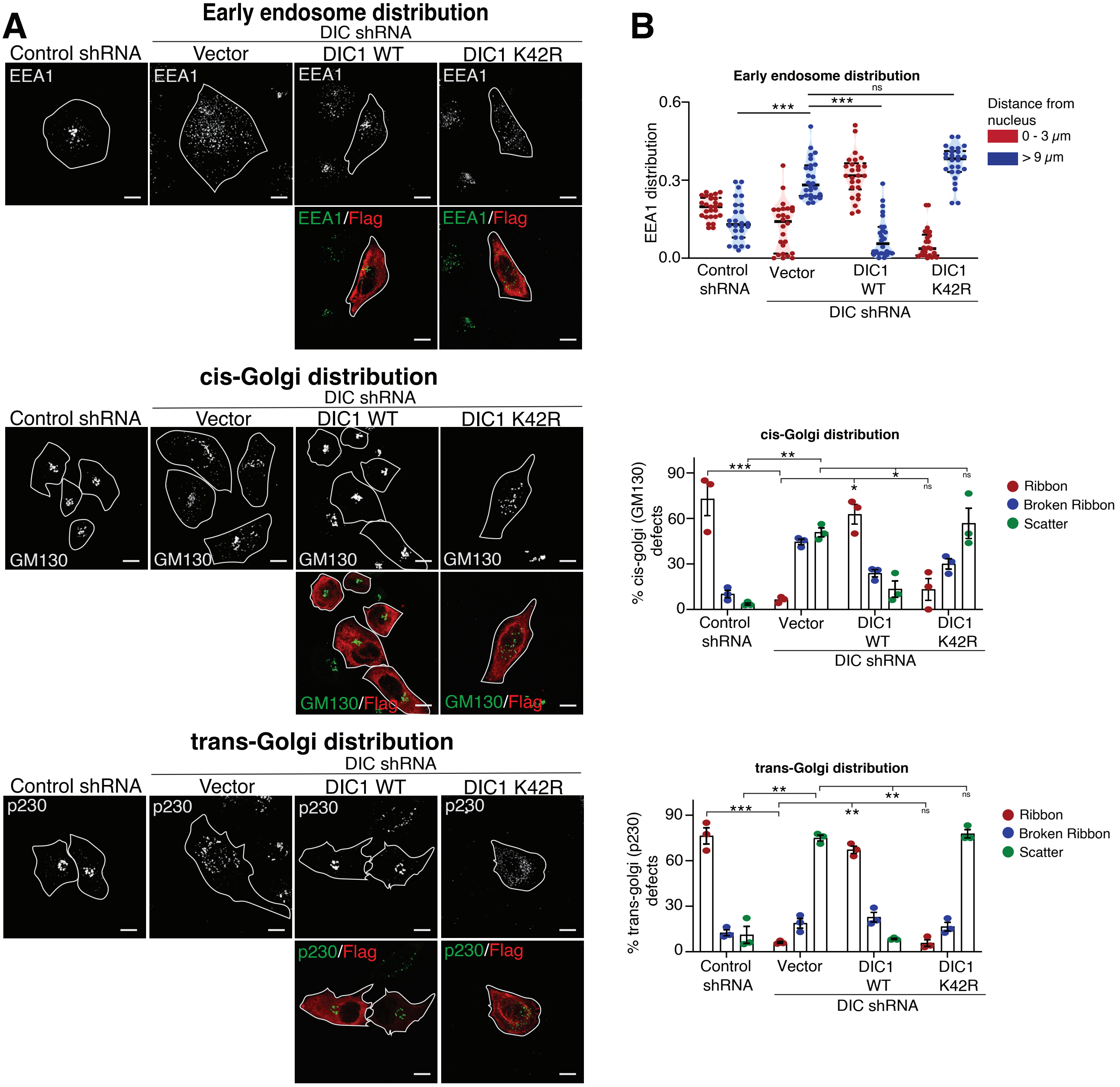

A
Early endosome distribution
DIC shRNA
Control shRNA
Vector
DIC1 WT
DIC1 K42R
EEA1
EEA1/Flag
cis-Golgi distribution
GM130
GM130/Flag
trans-Golgi distribution
p230
p230/Flag
B
Early endosome distribution
EEA1 distribution
0.0
0.3
0.6
ns
***
Distance from nucleus
0 - 3 μm
> 9 μm
Control shRNA
Vector
DIC1 WT
DIC1 K42R
DIC shRNA
cis-Golgi distribution
% cis-golgi (GM130) defects
0
30
60
90
**
*
ns
Ribbon
Broken Ribbon
Scatter
trans-Golgi distribution
% trans-golgi (p230) defects

### Figure S8

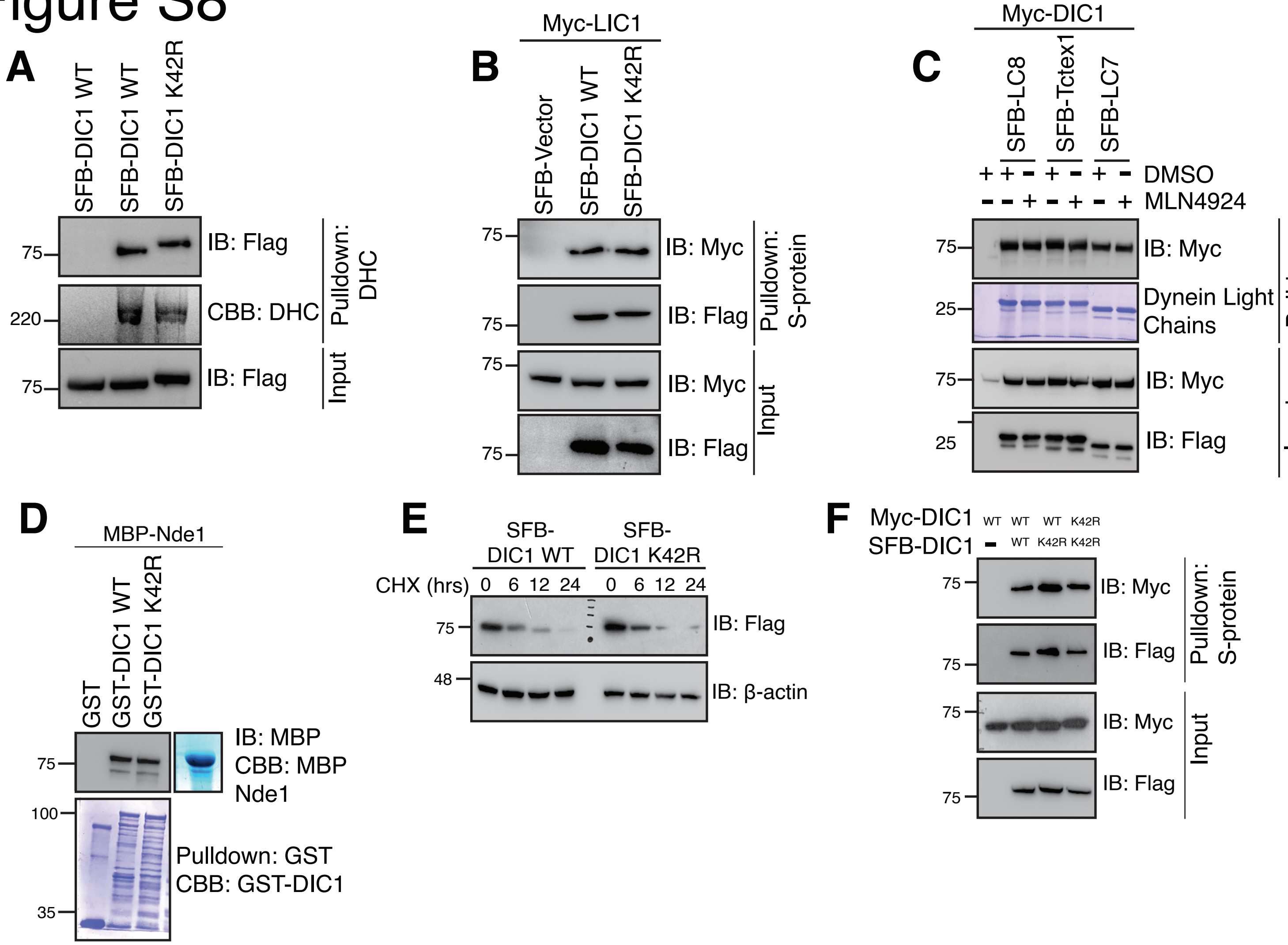

A
SFB-DIC1 WT
SFB-DIC1 WT
SFB-DIC1 K42R
75
IB: Flag
220
CBB: DHC
Pulldown: DHC
75
IB: Flag
Input
B
Myc-LIC1
SFB-Vector
SFB-DIC1 WT
SFB-DIC1 K42R
75
IB: Myc
75
IB: Flag
Pulldown: S-protein
75
IB: Myc
75
IB: Flag
Input
C
Myc-DIC1
SFB-LC8
SFB-Tctex1
SFB-LC7
DMSO
MLN4924
75
IB: Myc
25
Dynein Light Chains
Pulldown: S-protein
75
IB: Myc
25
IB: Flag
Input
D
MBP-Nde1
GST
GST-DIC1 WT
GST-DIC1 K42R
75
IB: MBP
CBB: MBP Nde1
100
35
Pulldown: GST
CBB: GST-DIC1
E
SFB-DIC1 WT
SFB-DIC1 K42R
CHX (hrs)
0 6 12 24
0 6 12 24
75
IB: Flag
48
IB: β-actin
F
Myc-DIC1
SFB-DIC1
WT WT WT K42R
WT K42R K42R
75
IB: Myc
75
IB: Flag
Pulldown: S-protein
75
IB: Myc
75
IB: Flag
Input

### Figure S9

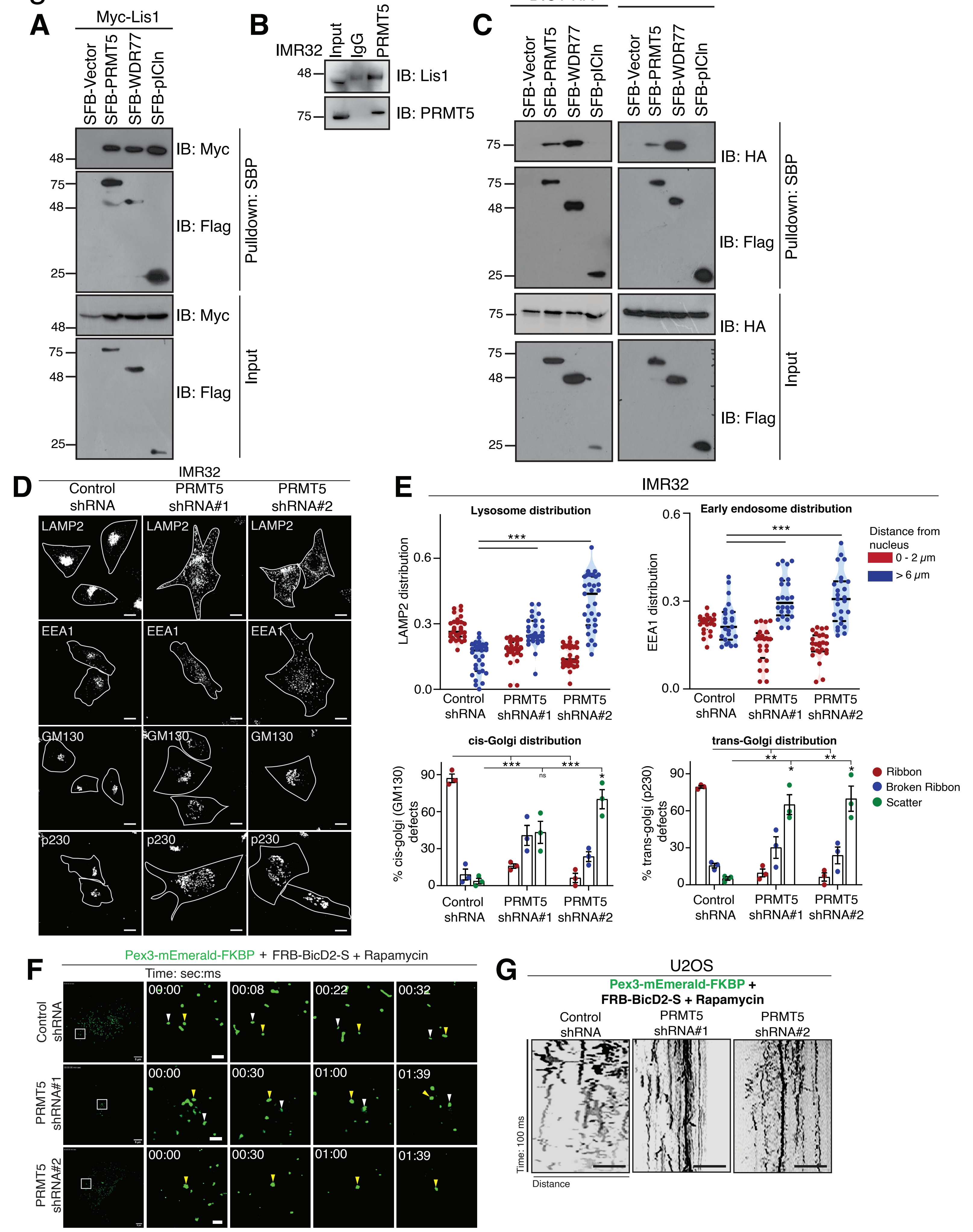

A
Myc-Lis1
SFB-Vector
SFB-PRMT5
SFB-WDR77
SFB-pICln
IB: Myc
IB: Flag
Pulldown: SBP
Input
B
IMR32
Input
IgG
PRMT5
IB: Lis1
IB: PRMT5
C
DIC1-HA
DIC2-HA
IB: HA
IB: Flag
D
IMR32
Control shRNA
PRMT5 shRNA#1
PRMT5 shRNA#2
LAMP2
EEA1
GM130
p230
E
Lysosome distribution
Early endosome distribution
LAMP2 distribution
EEA1 distribution
Distance from nucleus
0 - 2 μm
> 6 μm
cis-Golgi distribution
trans-Golgi distribution
% cis-golgi (GM130) defects
% trans-golgi (p230) defects
Ribbon
Broken Ribbon
Scatter
F
Pex3-mEmerald-FKBP + FRB-BicD2-S + Rapamycin
Time: sec:ms
G
U2OS
Time: 100 ms
Distance

### Figure S10

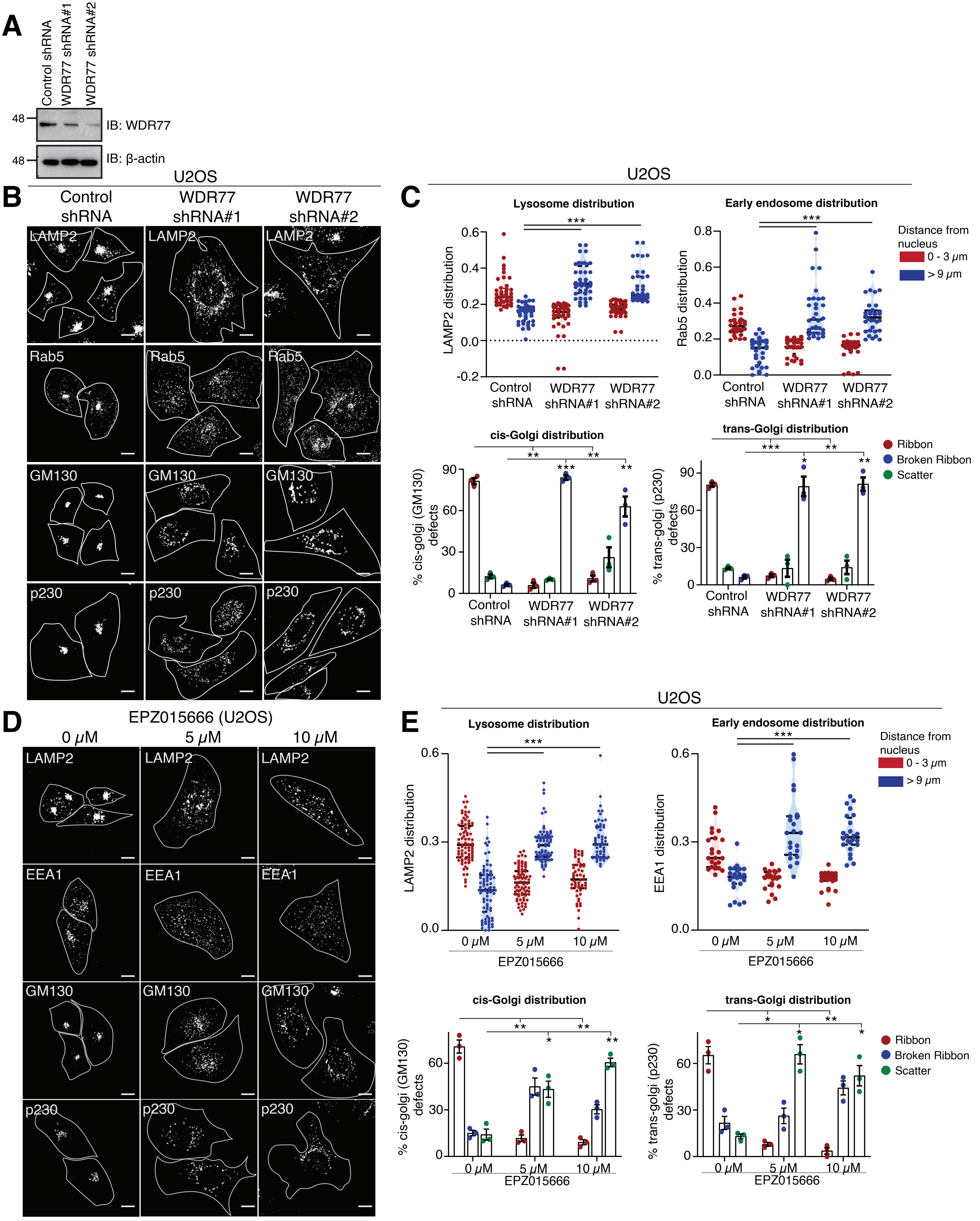

A
Control shRNA
WDR77 shRNA#1
WDR77 shRNA#2
48
IB: WDR77
48
IB: β-actin
B
U2OS
Control shRNA
WDR77 shRNA#1
WDR77 shRNA#2
LAMP2
Rab5
GM130
p230
C
U2OS
Lysosome distribution
LAMP2 distribution
Early endosome distribution
Rab5 distribution
Distance from nucleus
0 - 3 μm
> 9 μm
cis-Golgi distribution
% cis-golgi (GM130) defects
trans-Golgi distribution
% trans-golgi (p230) defects
Ribbon
Broken Ribbon
Scatter
D
EPZ015666 (U2OS)
0 μM
5 μM
10 μM
LAMP2
EEA1
GM130
p230
E
U2OS
Lysosome distribution
LAMP2 distribution
Early endosome distribution
EEA1 distribution
EPZ015666
Distance from nucleus
0 - 3 μm
> 9 μm
cis-Golgi distribution
% cis-golgi (GM130) defects
trans-Golgi distribution
% trans-golgi (p230) defects
Ribbon
Broken Ribbon
Scatter

### Figure S11

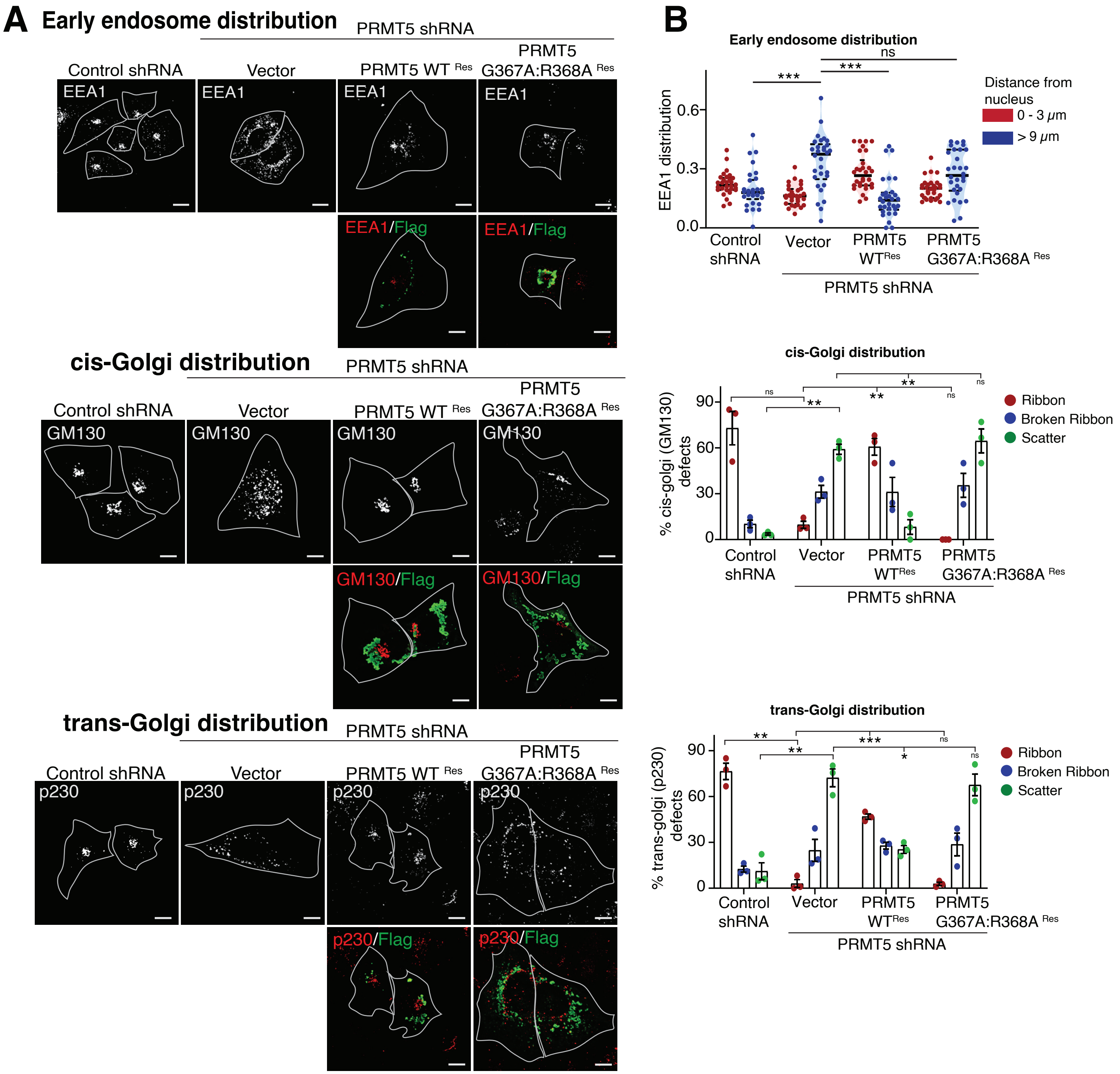
A
Early endosome distribution
PRMT5 shRNA
Control shRNA
Vector
PRMT5 WT Res
PRMT5 G367A:R368A Res
EEA1
EEA1/Flag
cis-Golgi distribution
GM130
GM130/Flag
trans-Golgi distribution
p230
p230/Flag
B
Early endosome distribution
EEA1 distribution
Distance from nucleus
0 - 3 μm
> 9 μm
cis-Golgi distribution
% cis-golgi (GM130) defects
Ribbon
Broken Ribbon
Scatter
trans-Golgi distribution
% trans-golgi (p230) defects
Control shRNA
Vector
PRMT5 WTRes
PRMT5 G367A:R368A Res
PRMT5 shRNA

### Figure S12

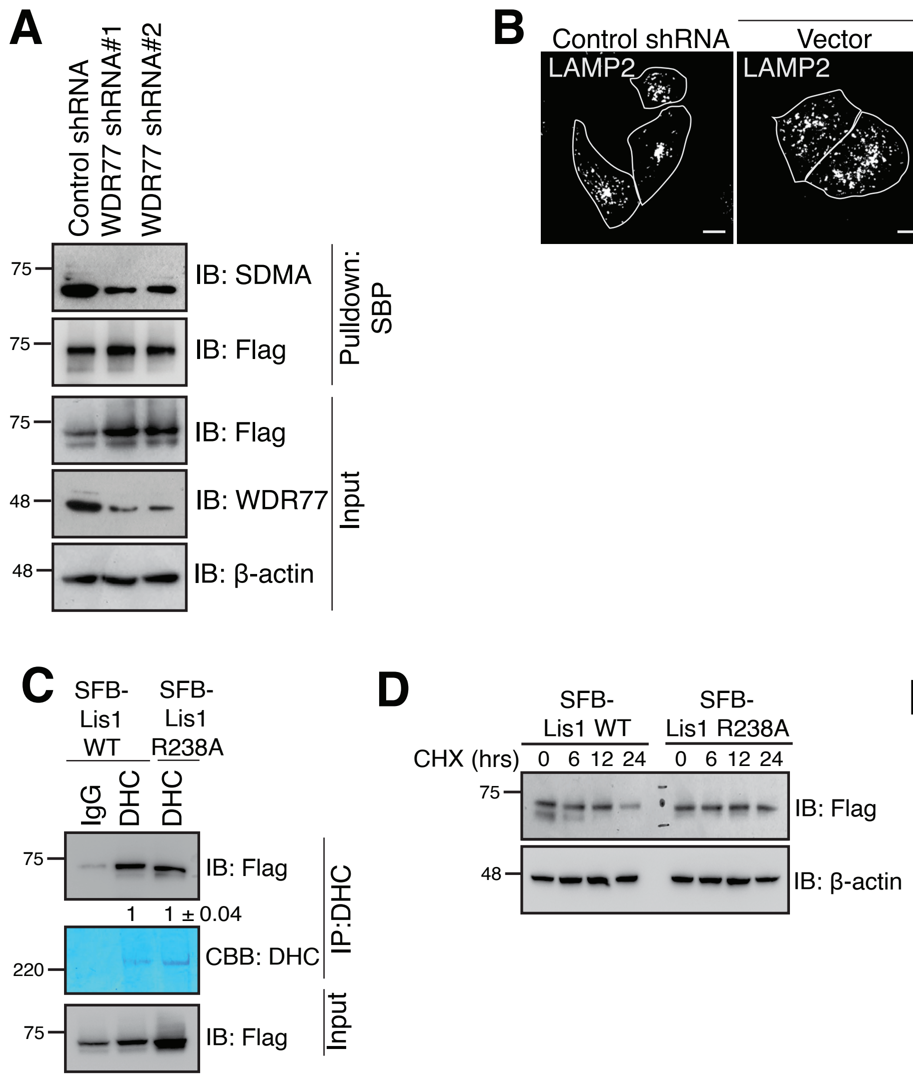

A
Control shRNA
WDR77 shRNA#1
WDR77 shRNA#2
75
IB: SDMA
75
IB: Flag
Pulldown: SBP
75
IB: Flag
48
IB: WDR77
48
IB: β-actin
Input
B
Lis1 shRNA
Control shRNA
Vector
Lis1 WTRes
Lis1 R238ARes
Lis1 R342ARes
LAMP2
LAMP2/Flag
C
SFB-Lis1 WT
SFB-Lis1 R238A
IgG
DHC
DHC
75
IB: Flag
1
1 ± 0.04
CBB: DHC
220
IP:DHC
75
IB: Flag
Input
D
SFB-Lis1 WT
SFB-Lis1 R238A
CHX (hrs)
0 6 12 24
0 6 12 24
75
IB: Flag
48
IB: β-actin
E
Myc-Lis1
WT WT WT RA
SFB-Lis1
- WT RA RA
63
IB: Myc
75
IB: Flag
Pulldown: S-protein
63
IB: Myc
75
IB: Flag
Input

### Figure S13

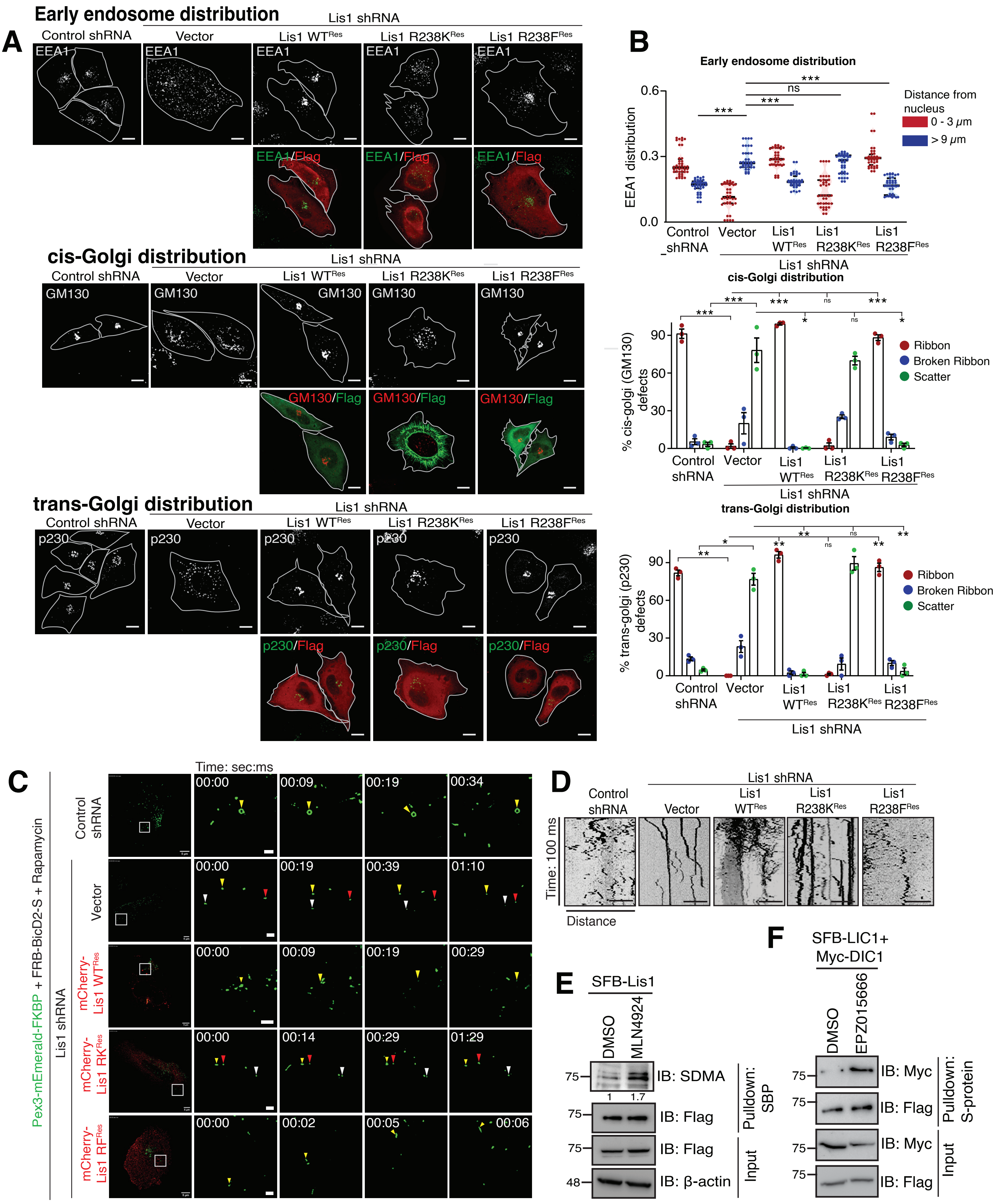

A
Early endosome distribution
Lis1 shRNA
Control shRNA
Vector
Lis1 WTRes
Lis1 R238KRes
Lis1 R238FRes
EEA1
EEA1/Flag
B
Early endosome distribution
EEA1 distribution
Distance from nucleus
0 - 3 μm
> 9 μm
ns
***
cis-Golgi distribution
GM130
GM130/Flag
% cis-golgi (GM130) defects
Ribbon
Broken Ribbon
Scatter
trans-Golgi distribution
p230
p230/Flag
% trans-golgi (p230) defects
C
Time: sec:ms
Pex3-mEmerald-FKBP + FRB-BicD2-S + Rapamycin
Control shRNA
Lis1 shRNA
Vector
mCherry-Lis1 WTRes
mCherry-Lis1 RKRes
mCherry-Lis1 RFRes
D
Time: 100 ms
Distance
E
SFB-Lis1
DMSO
MLN4924
IB: SDMA
IB: Flag
IB: β-actin
Pulldown: SBP
Input
F
SFB-LIC1+ Myc-DIC1
EPZ015666
IB: Myc
Pulldown: S-protein

**Table S1**

| S. No. | List of proteins in Lis1 purification (Gene names) | Unique peptides |
|---|---|---|
| 1 | PAFAH1B1 | 50 |
| 2 | DYNC1H1 | 25 |
| 3 | DYNC1I1 | 1 |
| 4 | DYNC1I2 | 18 |
| 5 | DYNC1LI1 | 17 |
| 6 | DYNC1LI2 | 14 |
| 7 | DYNLRB2 | 1 |
| 8 | DYNLT1 | 1 |
| 9 | DCTN1 | 27 |
| 10 | DCTN2 | 15 |
| 11 | DCTN3 | 2 |
| 12 | DCTN4 | 7 |
| 13 | NUDC | 27 |
| 14 | NDE1 | 27 |
| 15 | GPS1 | 8 |
| 16 | COPS2 | 20 |
| 17 | COPS3 | 9 |
| 18 | COPS4 | 10 |
| 19 | COPS5 | 12 |
| 20 | COPS6 | 5 |
| 21 | COPS7A | 5 |
| 22 | COPS7B | 2 |
| 23 | COPS8 | 4 |
| 24 | PRMT5 | 11 |
| 25 | WDR77 | 3 |
| 26 | CLNS1A | 6 |
| 27 | PAFAH1B2 | 13 |
| 28 | PAFAH1B3 | 12 |
| 29 | CUL4A | 8 |
| 30 | CUL7 | 4 |
| 31 | HSPA1B | 24 |
| 32 | HNRNPM | 22 |
| 33 | HADHB | 21 |
| 34 | HSPA5 | 21 |
| 35 | HSPD1 | 19 |
| 36 | DNAJA1 | 17 |
| 37 | DYNC1LI1 | 17 |
| 38 | PPM1G | 16 |
| 39 | GRIPAP1 | 16 |
| 40 | AIFM1 | 16 |
| 41 | USP47 | 16 |
| 42 | DISC1 | 15 |
| 43 | GTF2I | 15 |
| 44 | DHX9 | 14 |

| | | |
|---|---|---|
| 45 | NCL | 14 |
| 46 | TUFM | 14 |
| 47 | DDX21 | 14 |
| 48 | DNAJA2 | 13 |
| 49 | HSP90AA1 | 13 |
| 50 | USP7 | 13 |
| 51 | DNAJC7 | 13 |
| 52 | EEF2 | 13 |
| 53 | HSPA1L | 12 |
| 54 | RPS3 | 12 |
| 55 | HUWE1 | 12 |
| 56 | ATP5A1 | 12 |
| 57 | IRS4 | 12 |
| 58 | PPP6R3 | 12 |
| 59 | PABPC1 | 12 |
| 60 | DDX17 | 12 |
| 61 | SART3 | 12 |
| 62 | EEF1A1 | 12 |
| 63 | RPL3 | 11 |
| 64 | BAG5 | 11 |
| 65 | RUVBL2 | 11 |
| 66 | HSPH1 | 11 |
| 67 | HSPB1 | 10 |
| 68 | RUVBL1 | 10 |
| 69 | HSPA4 | 10 |
| 70 | DHX30 | 10 |
| 71 | SLC25A13 | 10 |
| 72 | RUVBL1 | 10 |
| 73 | HSP90AB1 | 10 |
| 74 | RPL7 | 10 |
| 75 | CCT6B | 9 |
| 76 | SLC25A6 | 9 |
| 77 | RPL4 | 9 |
| 78 | RPL3 | 9 |
| 79 | NPM1 | 9 |
| 80 | AP4B1 | 5 |
| 81 | RPS7 | 5 |
| 82 | CKAP4 | 5 |
| 83 | GTF3C5 | 5 |
| 84 | DNAJC10 | 5 |
| 85 | ILF3 | 5 |
| 86 | RPS27 | 5 |
| 87 | HSP90AB3P | 5 |
| 88 | ILF2 | 5 |
| 89 | AHCYL1 | 5 |
| 90 | HNRNPA1L2 | 4 |
| 91 | ACTA2 | 4 |
| 92 | RPL17 | 4 |

| | | |
|---|---|---|
| 93 | RPL31 | 4 |
| 94 | IGF2BP3 | 4 |
| 95 | SLC25A3 | 4 |
| 96 | DHX15 | 3 |
| 97 | GNL3 | 3 |
| 98 | PCBP1 | 3 |
| 99 | RPL28 | 3 |
